# The complete sequence of a human genome

**DOI:** 10.1101/2021.05.26.445798

**Authors:** Sergey Nurk, Sergey Koren, Arang Rhie, Mikko Rautiainen, Andrey V. Bzikadze, Alla Mikheenko, Mitchell R. Vollger, Nicolas Altemose, Lev Uralsky, Ariel Gershman, Sergey Aganezov, Savannah J. Hoyt, Mark Diekhans, Glennis A. Logsdon, Michael Alonge, Stylianos E. Antonarakis, Matthew Borchers, Gerard G. Bouffard, Shelise Y. Brooks, Gina V. Caldas, Haoyu Cheng, Chen-Shan Chin, William Chow, Leonardo G. de Lima, Philip C. Dishuck, Richard Durbin, Tatiana Dvorkina, Ian T. Fiddes, Giulio Formenti, Robert S. Fulton, Arkarachai Fungtammasan, Erik Garrison, Patrick G.S. Grady, Tina A. Graves-Lindsay, Ira M. Hall, Nancy F. Hansen, Gabrielle A. Hartley, Marina Haukness, Kerstin Howe, Michael W. Hunkapiller, Chirag Jain, Miten Jain, Erich D. Jarvis, Peter Kerpedjiev, Melanie Kirsche, Mikhail Kolmogorov, Jonas Korlach, Milinn Kremitzki, Heng Li, Valerie V. Maduro, Tobias Marschall, Ann M. McCartney, Jennifer McDaniel, Danny E. Miller, James C. Mullikin, Eugene W. Myers, Nathan D. Olson, Benedict Paten, Paul Peluso, Pavel A. Pevzner, David Porubsky, Tamara Potapova, Evgeny I. Rogaev, Jeffrey A. Rosenfeld, Steven L. Salzberg, Valerie A. Schneider, Fritz J. Sedlazeck, Kishwar Shafin, Colin J. Shew, Alaina Shumate, Yumi Sims, Arian F. A. Smit, Daniela C. Soto, Ivan Sović, Jessica M. Storer, Aaron Streets, Beth A. Sullivan, Françoise Thibaud-Nissen, James Torrance, Justin Wagner, Brian P. Walenz, Aaron Wenger, Jonathan M. D. Wood, Chunlin Xiao, Stephanie M. Yan, Alice C. Young, Samantha Zarate, Urvashi Surti, Rajiv C. McCoy, Megan Y. Dennis, Ivan A. Alexandrov, Jennifer L. Gerton, Rachel J. O’Neill, Winston Timp, Justin M. Zook, Michael C. Schatz, Evan E. Eichler, Karen H. Miga, Adam M. Phillippy

## Abstract

In 2001, Celera Genomics and the International Human Genome Sequencing Consortium published their initial drafts of the human genome, which revolutionized the field of genomics. While these drafts and the updates that followed effectively covered the euchromatic fraction of the genome, the heterochromatin and many other complex regions were left unfinished or erroneous. Addressing this remaining 8% of the genome, the Telomere-to-Telomere (T2T) Consortium has finished the first truly complete 3.055 billion base pair (bp) sequence of a human genome, representing the largest improvement to the human reference genome since its initial release. The new T2T-CHM13 reference includes gapless assemblies for all 22 autosomes plus Chromosome X, corrects numerous errors, and introduces nearly 200 million bp of novel sequence containing 2,226 paralogous gene copies, 115 of which are predicted to be protein coding. The newly completed regions include all centromeric satellite arrays and the short arms of all five acrocentric chromosomes, unlocking these complex regions of the genome to variational and functional studies for the first time.

## Introduction

The latest major update to the human reference genome was released by the Genome Reference Consortium (GRC) in 2013 and most recently patched in 2019 (GRCh38.p13) (*1*). This assembly traces its origin to the publicly funded Human Genome Project (*2*) and has been continually improved over the past two decades. Unlike the competing Celera assembly (*3*), and most modern genome projects that are also based on shotgun sequence assembly (*4*), the GRC human reference assembly is primarily based on Sanger sequencing data derived from bacterial artificial chromosome (BAC) clones that were ordered and oriented along the genome via radiation hybrid, genetic linkage, and fingerprint maps (*5*). This laborious approach resulted in what remains one of the most continuous and accurate reference genomes today. However, reliance on these technologies limited the assembly to only the euchromatic regions of the genome that could be reliably cloned into BACs, mapped, and assembled. Restriction enzyme biases led to the underrepresentation of many long, tandem repeats in the resulting BAC libraries, and the opportunistic assembly of BACs derived from multiple different individuals resulted in a mosaic assembly that does not represent a continuous haplotype. As such, the current GRC assembly contains several unsolvable gaps, where a correct genomic reconstruction is impossible due to incompatible structural polymorphisms associated with segmental duplications on either side of the gap (*6*). As a result of these shortcomings, many repetitive and polymorphic regions of the genome have been left unfinished or incorrectly assembled for over 20 years.

The current GRCh38.p13 reference genome contains 151 Mbp of unknown sequence distributed throughout the genome, including pericentromeric and subtelomeric regions, recent segmental duplications, ampliconic gene arrays, and ribosomal DNA (rDNA) arrays, all of which are necessary for fundamental cellular processes (**Fig. 1A**). Some of the largest reference gaps include the entire p-arms (short arms) of all five acrocentric chromosomes (Chr13, Chr14, Chr15, Chr21, and Chr22), and large human satellite arrays (e.g., Chr1, Chr9, and Chr16), which are currently represented in the reference simply as multi-megabase stretches of unknown bases (‘N’s). In addition to these apparent gaps, other regions of the current reference are artificial or are otherwise incorrect. The centromeric alpha satellite arrays, for example, are represented in GRCh38 as computationally generated models of alpha satellite monomers to serve as decoys for resequencing analyses (*7*). In the case of the acrocentrics, some sequence is included for the p-arm of Chromosome 21 but appears incorrectly localized and poorly assembled, resulting in false gene duplications that complicate downstream analyses (*8*). When compared to other human genomes, the current reference also shows a genome-wide deletion bias, suggesting the systematic collapse of repeats during its initial cloning and/or assembly (*9*).

**Fig. 1.**
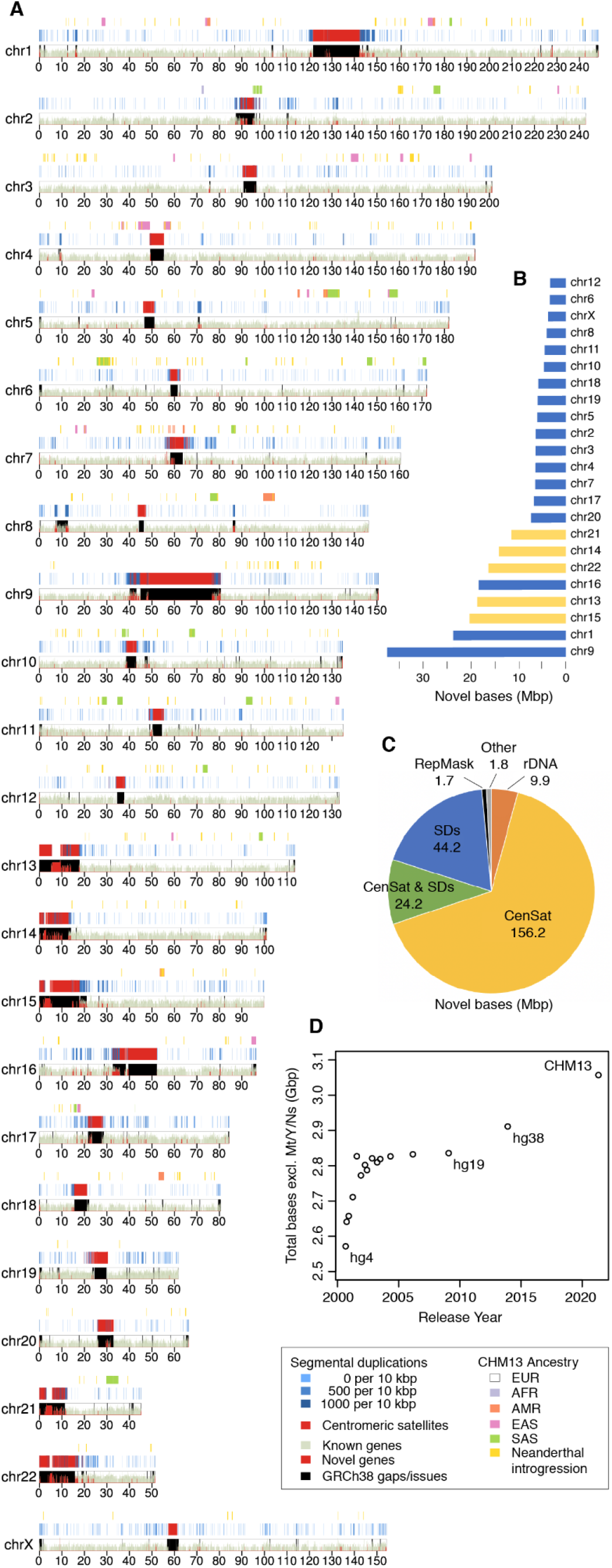
Summary of the complete T2T-CHM13 human genome assembly. **(A)** karyoploteR (*25*) ideogram of the T2T-CHM13v1.1 assembly improvements. The bottom track shows the density of known genes in green and new paralogs in red. GRCh38 gaps and issues that are resolved by the CHM13 assembly are highlighted by black rectangles. Above, the density of segmental duplications is given in blue (*26*) and centromeric satellites (CenSat) in red (*27*). The top track is a local ancestry analysis where the majority of the genome is predicted to be of European ancestry (1000 Genomes EUR), with regions of admixture colored as specified in the legend. **(B)** New bases in the CHM13 assembly relative to GRCh38 per chromosome, with the acrocentrics highlighted in yellow. **(C)** New or structurally variable bases added by sequence type (“CenSat & SDs” is the overlap between these two annotations). **(D)** Total non-gap bases in UCSC reference genome releases dating back to September 2000 (hg4) and ending with T2T-CHM13 in 2021.

Despite the functional importance of these missing or erroneous regions, the Human Genome Project was officially declared complete in 2003 (*10*), and there was limited progress towards closing the remaining gaps in the years that followed. This was largely due to limitations of its construction discussed above, but also due to the sequencing technologies of the time, which were dominated by low-cost, high-throughput methods capable of sequencing only a few hundred bases per read. Thus, shotgun-based assembly methods were unable to surpass the quality of the existing reference. However, recent advances in long-read genome sequencing and assembly methods have enabled the complete assembly of individual human chromosomes from telomere to telomere without gaps (*11, 12*). In addition to using long reads, these T2T projects have targeted the genomes of clonal, complete hydatidiform mole (CHM) cell lines, which are almost completely homozygous and therefore easier to assemble than heterozygous diploid genomes (*13*). This single-haplotype, *de novo* strategy overcomes the limitations of the GRC’s mosaic BAC-based legacy, bypasses the challenges of structural polymorphism, and allows the use of modern genome sequencing and assembly methods.

Application of long-read sequencing for the improvement of the human reference genome followed the introduction of PacBio’s single-molecule, polymerase-based technology (*14*). This was the first commercial sequencing technology capable of producing multi-kilobase sequence reads, which, even with a 15% error rate, proved capable of resolving complex forms of structural variation and gaps in GRCh38 (*9, 15*). The next major advance in sequencing read lengths came from Oxford Nanopore’s single-molecule, nanopore-based technology, capable of sequencing “ultra-long” reads in excess of 1 Mbp (*16*), but again with an error rate of 15%. By spanning most genomic repeats, these ultra-long reads enabled highly continuous *de novo* assembly (*17*), including the first complete assemblies of a human centromere (ChrY) (*18*) and a human chromosome (ChrX) (*11*). However, due to their high error rate, these long-read technologies have posed considerable algorithmic challenges, especially for the reliable assembly of long, highly similar repeat arrays (*19*). Improved sequencing accuracy simplifies the problem, but past technologies have excelled at either accuracy or length, not both. PacBio’s recent “HiFi” circular consensus sequencing offers a compromise of 20 kbp read lengths and a median accuracy of 99.9% (*20, 21*), which has resulted in unprecedented assembly accuracy with relatively minor adjustments to standard assembly approaches (*22, 23*). Whereas ultra-long nanopore sequencing excels at spanning long, identical repeats, HiFi sequencing excels at differentiating subtly diverged repeat copies or haplotypes.

In order to create a complete and gapless human genome assembly, we leveraged the complementary aspects of PacBio HiFi and Oxford Nanopore ultra-long read sequencing, combined with the essentially haploid nature of the CHM13hTERT cell line (hereafter, CHM13) (*24*). The resulting T2T-CHM13 reference assembly removes a 20-year-old barrier that has hidden 8% of the genome from sequence-based analysis, including all centromeric regions and the entire short arms of five human chromosomes. Here we describe the construction, validation, and initial analysis of the first truly complete human reference genome and discuss its potential impact on the field.

## Cell line and sequencing

As with many prior reference genome improvement efforts (*1, 9, 13, 24, 28, 29*), including the T2T assemblies of human chromosomes X (*11*) and 8 (*12*), we utilized a complete hydatidiform mole for sequencing. CHM genomes arise from the loss of the maternal complement and duplication of the paternal complement postfertilization and are, therefore, homozygous for one set of alleles. This simplifies the genome assembly problem by removing the confounding effect of heterozygous variation. We selected CHM13 for its stable 46,XX karyotype compared to other CHMs (*11*), but later found that CHM13 does possess a low level of heterozygosity, notably including a megabase-scale heterozygous deletion within the rDNA array on Chromosome 15, which was revealed by both FISH and nanopore sequencing (Figs. S1-2, Note S1). This and other identified heterozygous variants appear fixed in CHM13 and may have arisen during growth of the mole or passaging of the cell line. Local ancestry analysis shows the majority of the CHM13 genome is of European origin, including regions of Neanderthal introgression, with some predicted admixture from other populations (*30*) (**Fig. 1A**, Note S2).

Over the past 6 years, we have extensively sequenced CHM13 with multiple technologies (Note S3), including 30× PacBio circular consensus sequencing (HiFi) (*29*), 120× Oxford Nanopore ultra-long read sequencing (ONT) (*11, 12*), 100× Illumina PCR-Free sequencing (ILMN) (*1*), 70× Illumina / Arima Genomics Hi-C (Hi-C) (*11*), BioNano optical maps (*11*), and Strand-seq (*29*). Here we developed new methods for assembly, polishing, and validation that better utilize these datasets. In contrast to the first T2T assembly of Chromosome X (*11*)—which relied on ONT sequencing to create a backbone that was then polished with other technologies—we shifted to a new strategy that leverages the combined accuracy and length of HiFi reads to enable assembly of highly repetitive centromeric satellite arrays and closely related segmental duplications (*12, 22, 29*).

## Genome assembly

The basis of the T2T-CHM13 assembly is a high-resolution assembly string graph (*31*) built directly from HiFi reads. In a bidirected string graph, nodes represent unambiguously assembled sequences and edges correspond to the overlaps between them, due to either repeats or true adjacencies in the underlying genome. The HiFi-based string graph was constructed using a purpose-built method that combines components from the HiCanu (*22*) and Miniasm (*32*) assemblers along with specialized graph processing. Although HiFi reads are very accurate, their primary error mode is small insertions or deletions within homopolymer runs, so, like HiCanu, the first step of the T2T string graph construction process was to “compress” homopolymer runs in the reads to a single nucleotide (e.g., [A]_n_ becomes [A]_1_ for *n* > 1) (*33*). All compressed reads were then aligned to one another to identify and correct small errors, and differences within simple sequence repeats were masked to overcome this other known source of HiFi errors (*22*). After compression, correction, and masking, only exact overlaps were considered during graph construction, and new methods were developed for iterative graph simplification, as described in the supplementary methods (Fig. S3, Note S4). Edges in the resulting string graph correspond to exact overlaps of at least 8 kbp in homopolymer-compressed space.

In the resulting graph, most chromosomes are represented by one or more connected components, each having a mostly linear structure (**Fig. 2A**). This suggests very few perfect repeats greater than roughly 10 kbp exist between different chromosomes or distant loci, with the exception of the five acrocentric chromosomes, which form a single connected component in the graph. Another complex region is the HSat3 array on Chromosome 9, which includes a recent multi-megabase tandem HSat3 duplication consistent with the 9qh+ (*34*) karyotype of CHM13 (Fig. S4). Minor fragmentation of the chromosomes into multiple connected components resulted from HiFi sequencing dropout across some GA-rich simple sequence repeats, presumably due to a bias of the HiFi sequencing or base-calling process (*22*). These gaps were later filled using a prior ONT-based assembly (CHM13v0.7) (*11*).

**Fig. 2.**
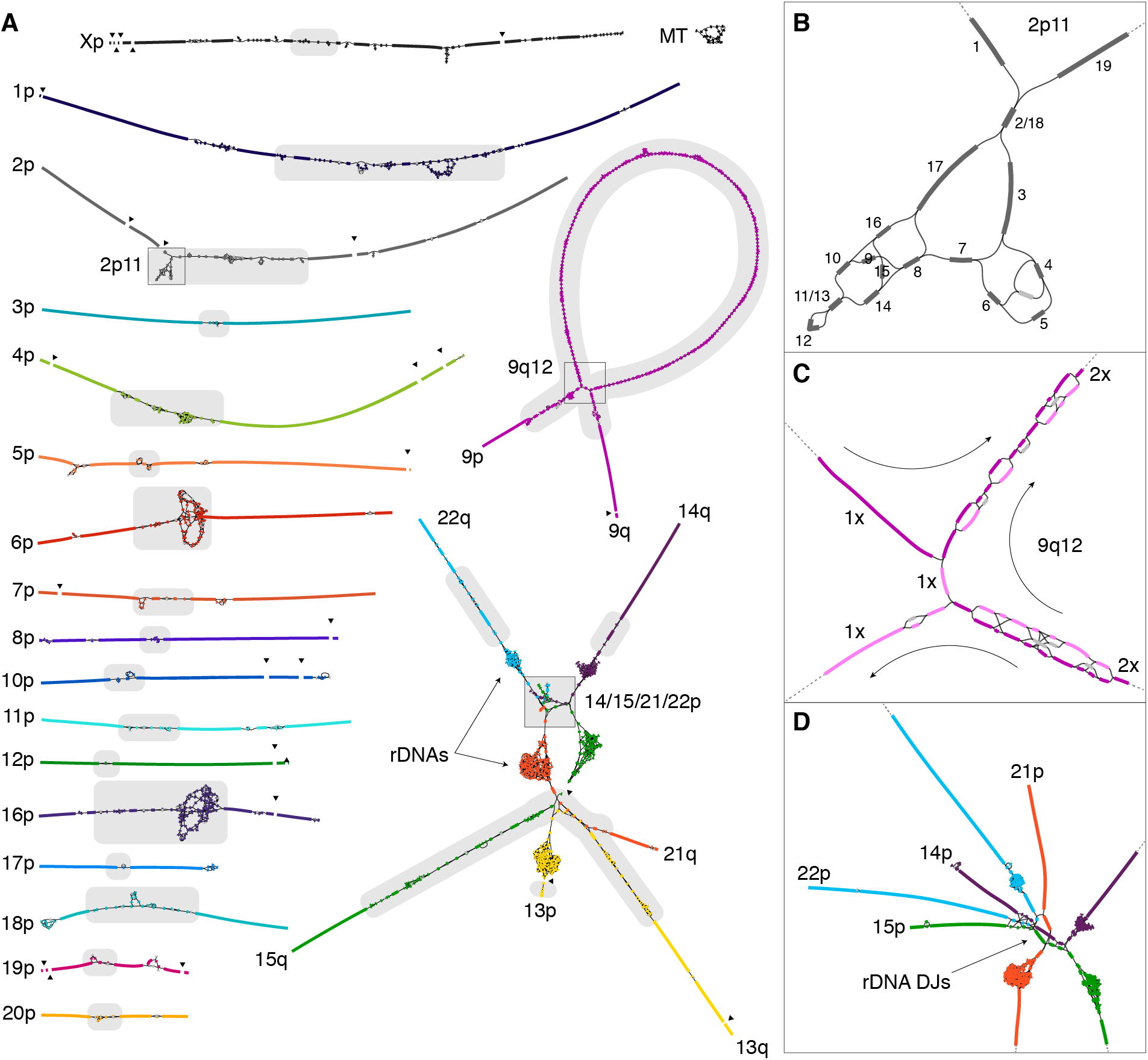
HiFi-based assembly string graph of the CHM13 genome. **(A)** Bandage (*36*) string graph visualization, where nodes represent unambiguously assembled sequences colored by source chromosome and scaled by length. Edges correspond to the overlaps between node sequences, due to either repeats or true adjacencies in the underlying genome. Centromeric satellite sequences are the source of most ambiguity in the graph (gray highlights). The graph is partially fragmented due to HiFi coverage dropout surrounding GA-rich sequence (black triangles). Local graph structures are enlarged with insets. The correct graph walks through these complex structures were resolved and confirmed with ultra-long ONT reads. **(B)** The identified graph traversal for the 2p11 locus is given by numerical order. Based on a depth-of-coverage analysis, the unlabeled light gray node represents an artifact or possible heterozygous variant and was not used. **(C)** The multi-megabase tandem HSat3 duplication (9qh+) at 9q12 requires two traversals of the large loop structure (note that the size of the loop is exaggerated because graph edges are of constant size). The first traversal of the loop is given in dark purple and the second in light purple. Nodes used by both traversals are also in dark purple and typically have twice the sequencing coverage. **(D)** The telomeric ends of four acrocentric p-arms form an ambiguous graph structure due to the highly similar sequence shared between all four chromosomes, specifically within the distal junction (DJ) sequence adjacent to the rDNA arrays, which themselves form dense, but separate, clusters of small nodes.

Ideally, the complete sequence for each chromosome should exist as a walk through the string graph where some nodes may be traversed multiple times (repeats) and some not at all (errors and heterozygous variants) (Fig. S5). To help identify the correct walks, we estimated coverage depth and multiplicity of the string graph nodes (Note S4), which allowed most tangles to be resolved as unique walks visiting each node the appropriate number of times (**Fig. 2B-D**, Fig. S5). Most of these walks were identified via manual curation of the graph. Low coverage nodes (e.g., <15% the average coverage) were presumed to stem from sequencing errors or somatic variants and were either pruned from the graph or omitted from the walks. Half-coverage nodes arranged as simple bubbles were most often heterozygous variants and the longer of the two was chosen with ties broken arbitrarily. In the remaining cases, the correct path was ambiguous and required integration of ONT reads. Where possible, raw ONT reads were aligned to the HiFi-based string graph using GraphAligner (*35*) to guide the correct walk, but more elaborate strategies were required for the large and highly similar satellite array duplications on chromosomes 6 and 9 (Fig. S6, Note S4). Only the five rDNA arrays, constituting approximately 10 Mbp of sequence, were not resolved as walks through the string graph and required a specialized assembly approach (described below). With this exception, an accurate consensus sequence for each chromosome was obtained from a multi-alignment of HiFi reads corresponding to the selected graph walks (Note S4), resulting in the CHM13v0.9 draft assembly.

## Assembly validation and polishing

The first step of genome assembly validation is to confirm that the constructed assembly is consistent with the data used to generate it (*37*). To evaluate concordance between the reads and the assembly we mapped all available primary data, including HiFi, ONT, ILMN, Strand-seq, and Hi-C, to the v0.9 draft assembly using Winnowmap2 (*38*) for long reads and BWA (*39*) for short reads (Note S5). Structural variants were identified with Sniffles (*40*), and small variants were called with DeepVariant (*41*) for ILMN and HiFi and PEPPER-DeepVariant for ONT (*42*). Small variants were further filtered using Merfin (*43*) to exclude any corrections that were not supported by the underlying ILMN and HiFi reads. After manual curation to differentiate true errors from heterozygous variants and mapping artifacts, a total of 4 large variants and 993 small variants were corrected, 52% of which were small indels within homopolymers. An additional 44 large and 3,901 small heterozygous variants were cataloged during curation (*44*), including the hemizygous insertion of an hTERT vector on Chromosome 21, consistent with the immortalization process used to create the CHM13 line (this insertion was excluded from the final assembly). The assembled sizes of major repeat arrays were consistent with ddPCR copy-number estimates for those tested (Tables S1-2, Fig. S7), and both Strand-seq (Figs. S8-9) and Hi-C (Fig. S10) data were concordant with the overall structure of the assembly, showing no signs of misorientations or other large-scale structural errors. In addition, the assembly correctly resolved 644 of 647 previously sequenced CHM13 BACs at >99.99% identity, with the three unresolved BACs appearing to be errors in the BACs themselves rather than the T2T assembly (Figs. S11-14).

The entire validation process was then repeated on the polished assembly, and investigation of the remaining variant calls revealed additional base-calling errors within some telomeric [TTAGGG]_n_ repeats. These putative errors were primarily a result of decreased coverage by both HiFi and ONT technologies towards the telomeres and not flagged by the initial variant-calling pipeline due to a telomere-associated strand bias in the ONT data. Telomeric ONT reads were only found oriented in the direction of the chromosome end and never away from it, which led to low-confidence variant calls and omission from polishing. After further curation and adjustment of the variant calling strategy (Note S4), an additional 454 corrections were made to the telomeres using PEPPER (*42*), followed by addition of the rDNA arrays as described below, resulting in a gapless CHM13v1.1 assembly—the first telomere-to-telomere representation of a human genome.

Mapped sequencing read depth across the final assembly shows uniform coverage across all chromosomes (**Fig. 3A**), with 99.86% of the assembly within three standard deviations of the mean coverage for both HiFi and ONT (HiFi coverage 34.70 ± 7.03, ONT coverage 116.16 ± 16.96, excluding the mitochondrial genome). Ignoring the 10 Mbp of rDNA sequence, where most of the coverage deviation resides, 99.99% of the assembly is within three standard deviations (Note S5). This is consistent with uniform coverage of the genome and confirms both the overall accuracy of the assembly and the absence of aneuploidy in the sequenced CHM13 cells. Copy-number concordance with raw ILMN and HiFi data also increased with successive versions of the assembly (Figs. S15-16). Local coverage anomalies were, however, observed across multiple satellite arrays (Table S3, Note S6). Given the uniformity of coverage increases and decreases across these arrays, association with specific satellite classes, and the sometimes opposite effect observed for HiFi and ONT, we hypothesize that these anomalies are related to systematic biases introduced during either sample preparation (e.g., shearing bias) or sequencing (e.g., polymerase kinetics), rather than assembly error (Note S6). For example, HiFi coverage is consistently elevated across HSat2 and HSat3 arrays, while ONT coverage remains normal but with an apparent strand bias and reduced read lengths for HSat2 (**Fig. 3B-C**, Figs. S17-S21). On the other hand, both HiFi and ONT coverage is depleted across the AT-rich HSat1 arrays, with ONT reads also showing shorter read lengths (**Fig. 3D**, Figs. S17-18, Table S3). While the specific mechanisms require further investigation, prior studies have noted similar biases within certain satellite arrays and sequence contexts for both ONT and HiFi (*45, 46*).

**Fig. 3.**
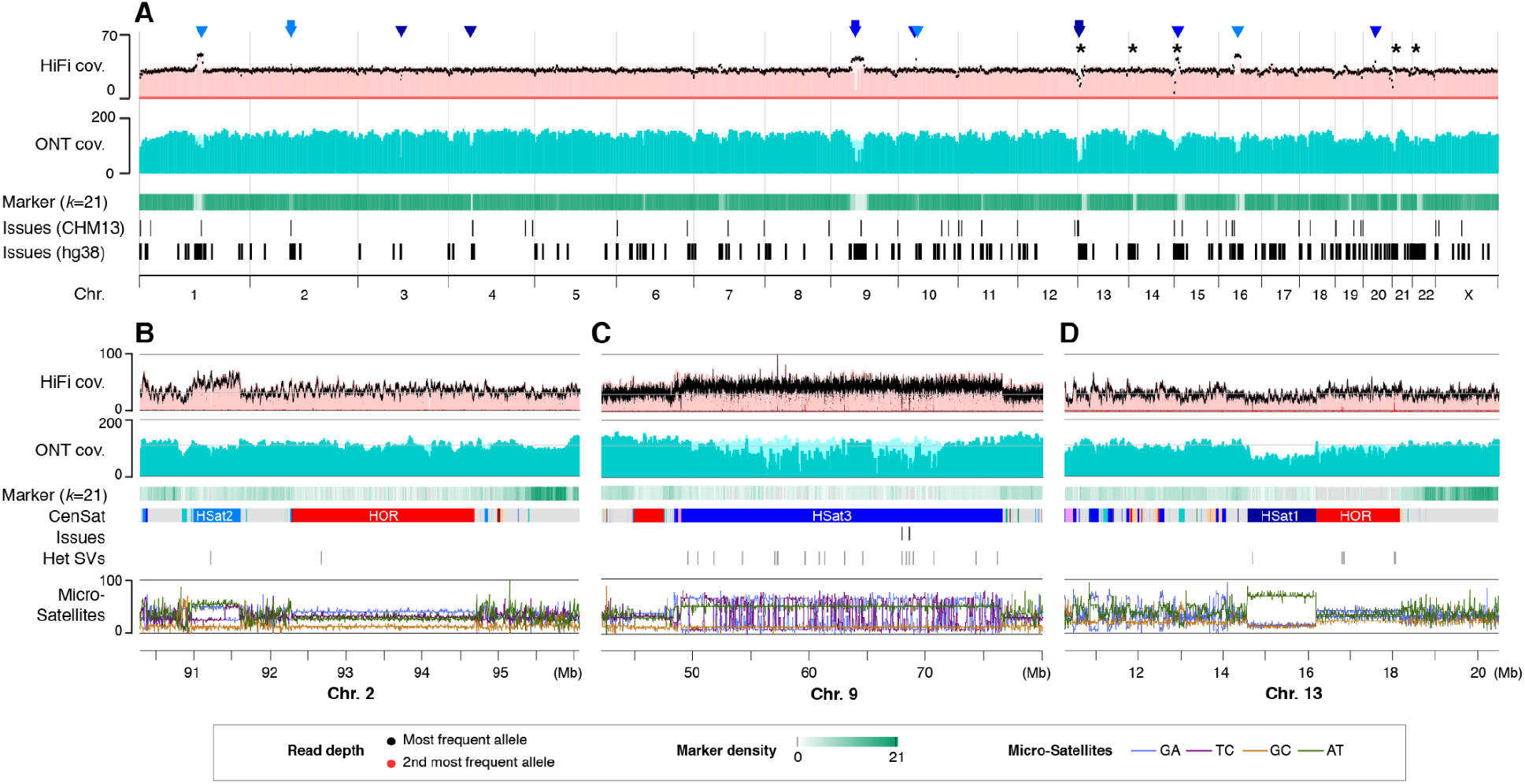
Sequencing coverage and assembly validation. Both HiFi and ONT sequencing reads mapped to the assembly show uniform coverage (cov.) across the whole genome as visualized by IGV (*51*), with the exception of certain human satellite repeat classes. Coverage deviations in these regions were found to be caused by sequencing biases associated with specific repeats rather than misassembly. **(A)** Whole-genome coverage of HiFi and ONT reads with primary alignments is shown in light shades and marker-assisted alignments overlaid in dark shades. Large HSat2 and HSat3 arrays are noted with light and dark blue triangles, respectively, and the location of the rDNA arrays is marked with asterisks (the inset regions are marked with arrowheads). Regions with low marker-assisted alignment coverage correspond with a lack of unique 21-mer markers (density shown in green), but are recovered by the primary alignments, albeit with low mapping quality. Suspected assembly issues in T2T-CHM13 are compared to known assembly gaps and issues in GRCh38/hg38 below, as reported by the GRC. **(B–D)** Enlargements corresponding to regions of the genome featured in Figure 2, along with annotations of the major satellite repeats contained (primarily HSat1, HSat2, HSat3, and alpha satellite HOR arrays). Elevated HiFi sequencing coverage is observed for HSat2 and HSat3, while reduced ONT coverage is observed for HSat1. Identified errors (Issues) and heterozygous variants (Het SVs) are shown below, which typically correspond with low HiFi coverage of the primary allele (black) and elevated coverage of a secondary allele (red). Microsatellite repeats (%) in every 128 bp window are shown at the bottom, labeled with dimer notation in homopolymer compressed space.

Due to the challenge of assembling them correctly, we performed targeted validation of all large satellite arrays and segmental duplications (Note S7). For centromeric alpha satellite arrays, we used the TandemTools package (*47*) to catalog additional variants that were missed by the standard approach. TandemTools was used throughout the process to guide development of the assembly method, and analysis of the final assembly shows high accuracy across all centromeric arrays (Fig. S22, Table S4). Independent ILMN-based copy number estimates of alpha satellite higher-order repeats (HOR) also correlate strongly with the assembly (Fig. S23). The beta satellite (BSat) and HSat arrays were separately validated by measuring the frequency of secondary variants identified by HiFi read mappings using a technique previously developed to identify collapsed segmental duplications (*48*). Because CHM13 is mostly homozygous, we expect to find very few heterozygous variants when mapping the raw reads back to the assembly and any variant clusters would indicate potential mis-assembly. This analysis shows consistent coverage across all satellite arrays, with only a handful of potential variants flagged (Fig. S24). A companion study (*26*) used this same approach to validate segmentally duplicated regions of the genome, along with an analysis of copy number variation compared to a collection of diverse human genomes, demonstrating that T2T-CHM13 represents these complex regions better than GRCh38.

In addition to high structural accuracy, we estimate the average consensus accuracy of the assembly to be between Phred Q67 and Q73 (Note S5), which is equivalent to 1 error per 10 Mbp and far exceeds the original Q40 definition of “finished” sequence (*49*). However, this represents an average across the entire genome and some regions are expected to be higher quality than others. In particular, regions of low HiFi coverage were found to be associated with an enrichment of potential consensus errors, as estimated from both HiFi and ILMN data (*44*). To guide future use of the assembly, we provide a curated list of all low-coverage and known heterozygous sites identified by the above validation procedures (Note S5). The total number of bases covered by potential issues in the T2T-CHM13 assembly is just 0.3% of the total assembly length compared to 8% for GRCh38 (**Fig. 3A**), making T2T-CHM13 a more complete, accurate, and representative reference sequence for both short- and long-read variant calling across human samples of all ancestries (*50*). Compared to GRCh38, T2T-CHM13 reduces false negative variant calls by adding 182 Mbp of novel sequence and removing 1.2 Mbp of falsely duplicated sequence, while simultaneously reducing false positive variant calls by fixing collapsed segmental duplications and other errors, affecting a total of at least 388 genes (68 protein coding) in GRCh38. Lastly, the T2T-CHM13 haplotype structure and SNP density is much more consistent than the mosaic GRCh38 when calling variants. A full comparison of GRCh38 versus CHM13 as a reference for variant calling is provided by Aganezov *et al*. (*50*), and a discussion of validation and polishing strategies for T2T genome assemblies by McCartney *et al*. (*44*).

## rDNA assembly

The most complex region of the HiFi string graph involves the human ribosomal DNA arrays and their surrounding sequence (**Fig. 2**). Human rDNAs are 45 kbp near-identical repeats that encode the 45S rRNA and are arranged in large, tandem repeat arrays embedded within the acrocentric p-arms. The length of these arrays varies widely between individuals (*52*) and even somatically, especially with aging and certain cancers (*53*). Based on ddPCR (Fig. S7) and whole-genome ILMN coverage (Figs. S15, S25), we estimate that the CHM13 genome contains approximately 200 rDNA copies per haplotype. Heterozygous structural variants were observed near the distal boundaries of the rDNA arrays on chromosomes 14, 15, and 21, including a megabase-scale heterozygous deletion within the Chromosome 15 array that was confirmed by FISH and a single 450 kbp spanning ONT read (Figs. S1-2, Note S1). In all cases we chose to include the variant that was most similar to the canonical rDNA unit, or in the case of Chromosome 15, the larger variant since it contained additional rDNA units. No other ONT reads spanned entire rDNA arrays, which is unsurprising given their multi-megabase size, and so additional copy number variation within the interior of the arrays could not be ruled out. However, all constructed assembly graphs exclude the presence of additional, non-rDNA, sequences within the arrays.

To assemble these highly dynamic regions of the genome, and overcome potential limitations of the string graph construction (Notes S4,8), we used a sparse de Bruijn graph approach (*54*). After recruiting HiFi reads from the rDNAs, we built a homopolymer-compressed sparse de Bruijn graph using a large word (*k*-mer) size, which formed a separate connected component for each rDNA array. We used this graph to separate all HiFi and ONT reads by chromosome, and then rebuilt the de Bruijn graph for each chromosome using a smaller word size and no homopolymer compression to improve connectivity. The resulting chromosome-specific graphs revealed a prominent central loop structure representing the 45 kbp rDNA unit surrounded by unique distal and proximal junctions (Fig. S26). ONT reads were then aligned to this graph with GraphAligner (*35*) to identify a set of walks, which were converted to sequence, segmented into individual rDNA units, and clustered into “morphs” according to their sequence similarity. Each morph represents a set of one or more nearly identical rDNA units, and most morphs are distinguishable by variable-length repeats within the intergenic spacer (Fig. S26). A consensus sequence was then constructed for each morph and copy number roughly estimated from the number of supporting ONT reads. Adjacency information was inferred from ONT reads spanning two or more rDNA units, and this was used to build a graph of morphs representing the structure of each rDNA array (Fig. S26). The most common morphs are self-adjacent in this graph, meaning they are arranged in continuous, head-to-tail, repeating blocks. The arrays on chromosomes 14 and 22 were found to contain a single primary morph repeated approximately 20 times and flanked by low-copy, more diverged units at the boundaries. In these cases, ONT reads extended far enough into the arrays to reconstruct the boundary units from the original HiFi string graph, and an appropriate number of polished primary morph copies were inserted into the center of the array. Chromosomes 13, 15, and 21 however, exhibited a more mosaic structure involving multiple, interspersed morphs, and the ONT reads were not long enough to determine the correct traversal of the morph graph. In these cases, after reconstructing the boundary units as before, the remaining primary morphs were polished and arranged in consecutive blocks, based on their coverage and order in the graph. The resulting (haploid) assembly contains 219 complete rDNA copies, totalling 9.9 Mbp of sequence. These rDNA assemblies approximate the true copy number and capture the common morphs on each chromosome, but the internal portions of arrays 13, 15, and 21 have been artificially ordered and should be treated as model sequences.

## A truly complete genome

The T2T-CHM13v1.1 assembly includes gapless telomere-to-telomere assemblies for all 22 human autosomes and Chromosome X, comprising 3,054,815,472 bp of nuclear DNA, plus a 16,569 bp mitochondrial genome (CHM13 does not have a Y chromosome). This complete assembly adds or corrects 238 Mbp of sequence compared to GRCh38, defined as regions of the T2T-CHM13 assembly that do not linearly align to GRCh38 over a 1 Mbp interval (i.e., are non-syntenic). The bulk of this sequence comprises centromeric satellites (180 Mbp), segmental duplications (68 Mbp), and rDNAs (10 Mbp), noting that there is overlap between regions identified as centromeric and segmentally duplicated (**Fig. 1B-C**). Of these regions 182 Mbp of sequence is uncovered by any primary alignments from GRCh38 and, therefore, completely novel to the CHM13 assembly. As a result, T2T-CHM13v1.1 substantially increases the number of known genes and repeats in the human genome (**Table 1**).

**Table 1.**
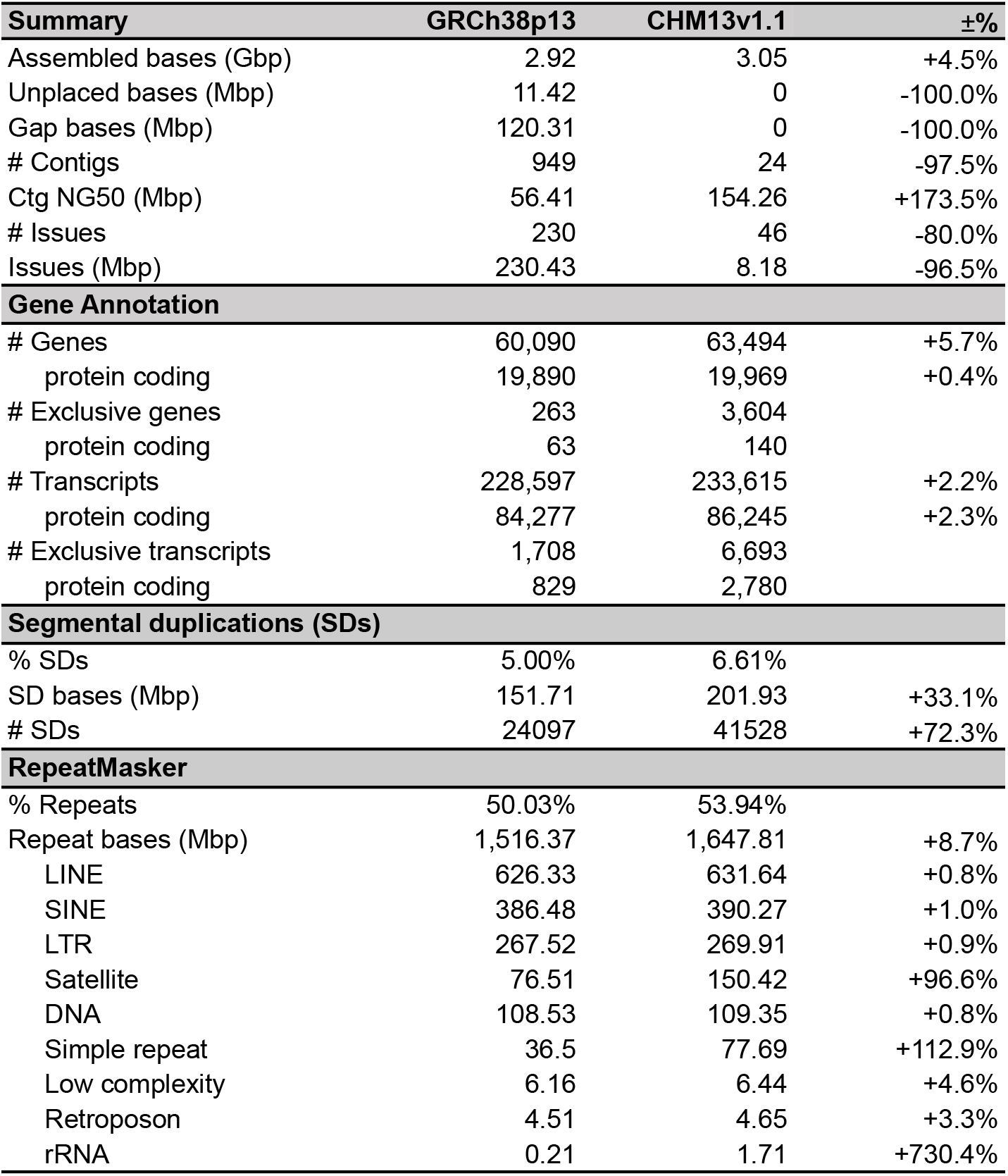
Comparison of GRCh38 and T2T-CHM13 human genome assemblies. GRCh38p13 summary statistics exclude “alts” (110 Mbp), patches (63 Mbp), and Chromosome Y (58 Mbp). Assembled bases: all non-N bases. Unplaced bases: not assigned or positioned within a chromosome. # Contigs: GRCh38 scaffolds were split at three consecutive Ns to obtain contigs. NG50: half of the 3.05 Gbp human genome size contained in contigs of this length or greater. # Exclusive genes/transcripts: for GRCh38, GENCODE genes/transcripts not found in CHM13; for CHM13, extra putative paralogs that are not in GENCODE. Segmental duplication analysis is from (*26*). RepeatMasker (*55*) analysis is from (*56*), with the “Unknown” category not shown.

| Summary | GRCh38p13 | CHM13v1.1 | ±% |
| --- | --- | --- | --- |
| Assembled bases (Gbp) | 2.92 | 3.05 | +4.5% |
| Unplaced bases (Mbp) | 11.42 | 0 | -100.0% |
| Gap bases (Mbp) | 120.31 | 0 | -100.0% |
| # Contigs | 949 | 24 | -97.5% |
| Ctg NG50 (Mbp) | 56.41 | 154.26 | +173.5% |
| # Issues | 230 | 46 | -80.0% |
| Issues (Mbp) | 230.43 | 8.18 | -96.5% |
| **Gene Annotation** |  |  |  |
| # Genes | 60,090 | 63,494 | +5.7% |
| protein coding | 19,890 | 19,969 | +0.4% |
| # Exclusive genes | 263 | 3,604 |  |
| protein coding | 63 | 140 |  |
| # Transcripts | 228,597 | 233,615 | +2.2% |
| protein coding | 84,277 | 86,245 | +2.3% |
| # Exclusive transcripts | 1,708 | 6,693 |  |
| protein coding | 829 | 2,780 |  |
| **Segmental duplications (SDs)** |  |  |  |
| % SDs | 5.00% | 6.61% |  |
| SD bases (Mbp) | 151.71 | 201.93 | +33.1% |
| # SDs | 24097 | 41528 | +72.3% |
| **RepeatMasker** |  |  |  |
| % Repeats | 50.03% | 53.94% |  |
| Repeat bases (Mbp) | 1,516.37 | 1,647.81 | +8.7% |
| LINE | 626.33 | 631.64 | +0.8% |
| SINE | 386.48 | 390.27 | +1.0% |
| LTR | 267.52 | 269.91 | +0.9% |
| Satellite | 76.51 | 150.42 | +96.6% |
| DNA | 108.53 | 109.35 | +0.8% |
| Simple repeat | 36.5 | 77.69 | +112.9% |
| Low complexity | 6.16 | 6.44 | +4.6% |
| Retroposon | 4.51 | 4.65 | +3.3% |
| rRNA | 0.21 | 1.71 | +730.4% |

To provide an initial annotation, we used both the Comparative Annotation Toolkit (CAT) (*57*) and Liftoff (*58*) to project the GENCODE v35 (*59*) reference annotation onto the new T2T-CHM13 assembly. Additionally, CHM13 Iso-Seq transcriptome data was processed using StringTie2 (*60*) to generate a set of *de novo* assembled transcripts, which were provided as complementary input to CAT. Genes identified by Liftoff, but missing from the CAT annotation, were then added to provide a comprehensive annotation of the newly assembled sequence (Note S9). This represents an initial draft annotation of the newly completed regions of the genome that will require further analysis and validation that is beyond the scope of this study.

The draft T2T-CHM13 annotation totals 63,494 genes and 233,615 transcripts, of which 19,969 and 86,245 are predicted to be protein coding, respectively, with 469 predicted frameshifts in 387 genes (**Table 1**, Fig. S27, Tables S5-7). Only 263 GENCODE genes (448 transcripts) are exclusive to GRCh38 and have no assigned ortholog in the CHM13 annotation (Fig. S28, Tables S8-9). At least 24 of these genes correspond to presumed false duplications and other errors in GRCh38 (*50*) (Fig. S29), while most others originate from repetitive regions that may be successfully annotated in the future. In comparison, a total of 3,604 genes (6,693 transcripts) are exclusive to CHM13 (Tables S10-11). Due to the segmentally duplicated nature of the newly assembled regions, these genes represent putative paralogs and localize to pericentromeric regions and the acrocentric p-arms, including 876 rRNA transcripts (**Fig. 1A**). Of these new genes, 140 are predicted to be protein coding based on their GENCODE paralogs and have a mean of 99.5% nucleotide and 98.7% amino acid identity to their most similar GRCh38 copy (Table S12). Due to CHM13’s more representative copy number in complex regions of the genome, many of these additions represent gene family expansions, which has the dual benefit of adding new paralogs to the reference and improving variant analyses of the previously known copies (*50*). While some of the new paralogs in the CHM13 annotation may also be present (but unannotated) in GRCh38, 2,226 of the genes exclusive to CHM13 (115 protein coding) fall within regions of the genome with no GRCh38 primary alignments (Note S9) and are verifiably novel.

## Acrocentric chromosomes

We focus here on describing the genomic structure of the newly completed p-arms of the five acrocentric chromosomes, which, despite their obvious importance for basic cellular function, have remained largely unsequenced to date. This omission has been due to the high enrichment for satellite repeats and segmental duplications throughout the acrocentric p-arms, which has prohibited *de novo* sequence assembly and limited their past characterization to cytogenetics (*61*), restriction mapping (*62*), and BAC sequencing (*63*–*65*). All five acrocentric p-arms follow a similar structure consisting of a variable-sized rDNA array embedded within distal and proximal satellite arrays, but the size of these arrays is highly polymorphic (*52, 53*).

Their importance to genome biology includes ribosome biogenesis, nucleolus formation (*66*), chromosomal instability (*67*), and known genetic conditions including ribosomopathies (*68*), Robertsonian translocations (*69*), and Down Syndrome (*70*). However, the lack of a reference sequence for these five chromosome arms has resulted in their exclusion from high-resolution genomic analysis and association studies.

The T2T-CHM13 assembly uncovers the complete sequence of all acrocentric p-arms for the first time, which vary in size from 10.1 Mbp to 16.7 Mbp each and amount to 66.1 Mbp of new sequence (**Fig. 4**). Each p-arm contains a single rDNA array varying in size from 0.7 Mbp (Chr14) to 3.6 Mbp (Chr13), surrounded by abundant satellite repeats and segmental duplications. Compared to other human chromosomes, the acrocentrics are also unusually similar to one another in terms of their p-arm sequence and composition and, as a result, were the only chromosomes not separated into individual components during string graph construction (**Fig. 2D**). Specifically, we find that 5 kbp windows on the acrocentric p-arms align with a median identity of 98.7% between chromosomes, creating many opportunities for interchromosomal exchange. This high degree of similarity is presumably due to recent non-allelic or ectopic recombination stemming from their colocalization in the nucleolus (*64*). Additionally, no 5 kbp window is unique considering alignments >80% identity, and 96% of the non-rDNA sequence can be found elsewhere in the genome using a similar criteria, suggesting that the acrocentric p-arms are frequent and dynamic sources of segmental duplication.

**Fig. 4.**
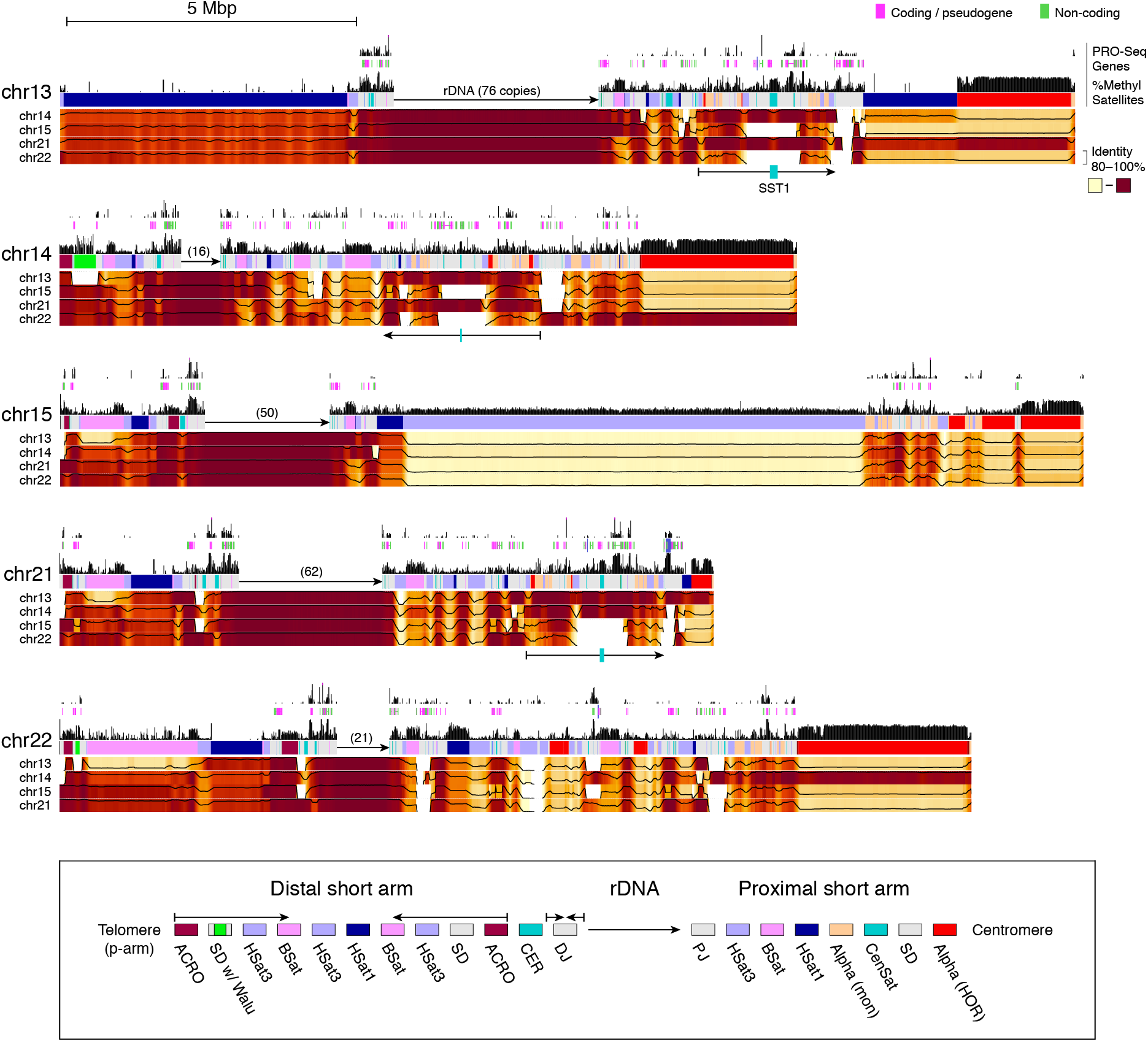
Short arms of the human acrocentric chromosomes in CHM13. Each p-arm is shown along with RNA polymerase activity (PRO-Seq), annotated genes, percent of methylated CpGs, and color-coded satellite repeat annotation. ILMN PRO-Seq alignments were filtered by unique 51-mers to avoid cross mapping in these repetitive regions, and so may appear sparser than reality. Methylation data is derived from ultra-long ONT reads and is more comprehensive. The rDNA arrays are represented only by a directional arrow and copy number due to their high self-similarity, which limits mapping. Below the annotations are percent identity heatmaps versus the other four p-arms, where sequence identity was computed in 10 kbp windows and smoothed over 100 kbp intervals. Each position shows the maximum identity of that window to any window in the other chromosome. The legend shows the names and organization of satellite repeats, which is generally conserved on the distal short arms, including a large symmetric duplication (long arrows) and a highly conserved inverted repeat in the DJ (short arrows). The proximal short arms show a greater diversity of structures, and so the legend simply lists some common elements. The proximal short arms of Chromosomes 13, 14, and 21 share a large segmentally duplicated core, including small alpha satellite HOR arrays and a central, highly methylated, SST1 array (long arrows with a central teal block). Chromosomes 13/21 and 14/22 also share a striking degree of similarity across the centromere and pericentromere.

Situated between the distal and proximal short arms, CHM13’s five rDNA arrays are in the expected arrangement, organized as head-to-tail tandem arrays with all 45S transcriptional units pointing towards the centromere. No inversions were noted within the arrays and the vast majority of rDNA units are full length, in contrast to some prior studies that reported embedded inversions and other non-canonical structures (*65, 71*). Each array appears highly homogenized, and there is more variation between rDNA units on different chromosomes than within chromosomes (Fig. S30), which suggests that intra-chromosomal exchange of rDNA units via non-allelic homologous recombination is more common than inter-chromosomal exchange. For example, most 45S gene copies on the same chromosome are 100.0% identical to one another (with some alternative morphs in the longer arrays), while the identity of the most frequent 45S morphs between chromosomes ranges from 99.4–99.7%. A minor chromosome 15 morph shows the most similarity to the KY962518.1 reference sequence, which was originally derived from a human Chromosome 21 BAC clone (*65*). These two sequences align across the entirety of the 45 kbp rDNA at 98.9% identity. The primary Chromosome 21 morph is also highly similar to KY962518.1, but includes a 96 bp deletion in the intergenic spacer (IGS). As expected, the 13 kbp 45S is more highly conserved than the IGS, with all major CHM13 45S morphs aligning between 99.4–99.6% identity to KY962518.1. Certain rDNA variants appear chromosome-specific in CHM13, including single-nucleotide variants within the 45S and its upstream promoter region (Fig. S31). The most evident variants are repeat expansions and contractions within the tandem repeat that immediately follows the 45S and the large, CT-rich simple sequence repeat located in the middle of the IGS. Most CHM13 rDNA morphs can be distinguished based solely on structural differences within these two polymorphic sites, and each array contains a major morph that is unique to that chromosome (Fig. S32).

Beginning with the p-arm telomere and extending to the rDNA array, the structure of all five distal short arms follows a similar pattern involving a symmetric arrangement of inverted segmental duplications and ACRO, HSat3, BSat, and HSat1 repeats (**Fig. 4**); however, the sizes of these repeat arrays varies widely among chromosomes. CHM13 Chromosome 13 is most unique in its distal structure, having lost half of the inverted satellite duplication and gained a massively expanded HSat1 array relative to the other four chromosomes. Despite their variability in size, all satellite arrays share a high degree of similarity (typically >90% identity) both within and between acrocentric chromosomes. Chromosomes 14 and 22 also feature the expansion of a newly discovered 64-bp Alu-associated satellite repeat (“Walu”) within the distal inverted duplication (*56*), the location of which was confirmed via FISH (Fig. S33). Several of the non-satellite sequences show evidence of transcription, including a number of gene predictions. The distal junction (DJ) immediately prior to the rDNA array includes centromeric repeats (CER) and a highly conserved and actively transcribed 200 kbp palindromic repeat, which agrees with the few hundred kilobases of DJ sequence previously characterized via BACs (*64, 72*).

Extending from the rDNA array to the centromere, the proximal short arms are larger in size and show a higher diversity of structures compared to the distal arrays, but they remain highly similar at the sequence level with each arm being entirely composed of shuffled segmental duplications, transposable element composite arrays (e.g. ACRO_COMP), and satellite repeats including HSat3, BSat, HSat1, HSat5, and various centromeric satellites including both monomeric and higher-order repeat (HOR) alpha satellite arrays. With the exception of Chromosome 13—which includes a large, proximal HSat1 array—HSat3 and BSat are most prevalent in terms of total bases. Chromosome 15 includes a distinctive 8 Mbp HSat3 array that comprises the majority of its proximal short arm. Many of the proximal BSat arrays show a previously unreported mosaic inversion structure that we also discovered in some HSat arrays elsewhere in the genome (*27*) (Fig. S34). All regions of segmental duplication on the acrocentric p-arms appear highly similar between chromosomes, often exceeding 99% identity, and these regions also show evidence of active transcription (*56*) (**Fig. 4**, PRO-Seq). Notably, the proximal short arms of chromosomes 13, 14, and 21 appear to share the highest degree of structural and sequence similarity, including a large region of segmental duplication with a central and highly methylated SST1 array (**Fig. 4**). This similarity coincides with these three chromosomes being most frequently involved in Robertsonian translocations (*73*), providing possible mechanistic insight into their formation. As previously described (*74, 75*), alpha satellite HORs in the arrays of active centromeres of chromosomes 13/21 and chromosomes 14/22 are almost identical within each pair, but not between them. However, small alpha satellite HOR arrays within segmental duplications of the proximal short arms show a different pattern (**Fig. 4**). Here, chromosomes 13, 14 and 21 share similar HOR subsets, while Chromosome 22 is distinguished by two copies of a segmental duplication in which a Chromosome 22-specific alpha satellite HOR has amplified (*27*). Using this new reference sequence as a basis, further study of additional genomes is needed to understand which features of the acrocentric p-arms observed in CHM13 are conserved across the human population.

## Analyses and resources

A number of companion studies were carried out to fully characterize the new sequences and structural variants revealed by the T2T-CHM13 assembly. These include comprehensive analyses of centromeres, segmental duplications, transcriptional and epigenetic profiles, mobile elements, and variant calls. Altemose *et al. (27)* carried out the first high-resolution study of satellite sequence organization at sites of centromere formation and discovered new sequences coincident with centromere-specific chromatin. Vollger *et al*. (*26*) identified nearly double the number of previously known near-identical segmental duplication alignments in the human genome, thereby identifying many new genes and regions susceptible to rearrangement via unequal crossing over. Additionally, comparison of T2T-CHM13, GRCh38, and high-quality primate assemblies revealed unprecedented patterns of structural heterozygosity and massive evolutionary differences in duplicated gene content between humans and the other apes *(26)*. Gershman *et al. (76)* developed methods for epigenetic profiling of complete human genomes and uncovered novel methylation patterns across all families of human repeats, including a hypomethylated and highly dense chromatin region within the centromere which may guide kinetochore formation. Hoyt *et al. (56)* carried out a comprehensive update of human repeat models and annotations, revealing previously unknown repeat arrays, transposable element (TE) derived composite elements, and transcriptionally active TEs embedded within large satellite arrays. Coupled with PRO-Seq (*77*) and the ONT-derived methylation data, sites of engaged RNA polymerase were annotated genome-wide, revealing unique transcriptional and epigenetic profiles across highly repetitive regions of the human genome. Lastly, Aganezov *et al*. (*50*) compared variant calling results for GRCh38 and T2T-CHM13, and observed improvements in both short- and long-read variant calling with the use of CHM13, including within medically relevant genes that were found to be mis-assembled or falsely duplicated within GRCh38.

These studies have generated a rich variety of omics datasets for CHM13 that are all browsable via a UCSC Assembly Hub (**Data and materials**). In addition to the sequencing data, assembly, and gene annotation described here, the browser provides a central resource for all results of the above noted companion studies, including detailed annotations of centromeres and satellite DNAs *(27)*; genome-wide repeat and transposable element annotations *(56)*; segmental duplications and population copy number estimates *(26)*; variant calls and mappability scores *(50)*; and whole-genome alignments to GRCh38. Importantly, the newly finished regions of the genome are readily accessible to long-read mapping and variant calling, and many others can be sparsely mapped with short reads using unique marker sequences (Fig. S35, Table S13, Note S11). We have surveyed these new regions using a variety of modern omics approaches, all of which are available in the browser, including RNA-Seq *(27)*, Iso-Seq *(26)*, PRO-Seq *(56)*, CUT&RUN *(27)*, and ONT methylation *(76)* experiments. An additional interactive dotplot visualization *(78)* is also provided to explore the newly uncovered genomic repeats. Finally, the CHM13hTERT cell line itself is in the process of being banked at Coriell (Camden, NJ, USA) and will be made available for research use, providing a physical reference material alongside the digital reference sequence, something that is not possible for GRCh38.

To highlight the utility of these genetic and epigenetic resources mapped to a complete human genome, we provide the specific example of Facioscapulohumeral muscular dystrophy (FSHD) region gene 1 *(FRG1*), which is a poorly understood candidate gene for FSHD located in the subtelomeric region of human Chromosome 4q35. The precise etiology of FSHD remains unclear due to complex epigenetic factors surrounding *FRG1* (*79*), *including the neighboring double homeobox 4 (DUX4*) transcription factor (*80*) and polymorphic D4Z4 macrosatellite array (*81*). A near-identical duplication of the D4Z4 array on Chromosome 10q26 was previously characterized, but the T2T-CHM13 assembly reveals 23 paralogs of *FRG1* duplicated throughout previously unfinished regions of chromosomes 9, 13, 14, 15, 20, 21, and 22 (**Fig. 5**). These duplicated genes were previously identified by fluorescence *in situ* hybridization (*82*) and underwent recent amplification in the great apes (*83*), but only 9 paralogs are found in GRCh38, hampering sequence-based analysis. The new paralogs’ association with all five acrocentrics is particularly notable, given that *FRG1* also localizes to nucleoli and is thought to play a role in RNA biogenesis (*84*). One of the few paralogs included in GRCh38, *FRG1DP*, is located in the centromeric region of Chromosome 20 and shares high identity (97%) with some of the newly assembled paralogs (*FRG1BP4*–*10*) situated in or near the acrocentrics (Fig. S36, Table S14-15, Note S12). When aligning HiFi reads to GRCh38, absence of these paralogs from the reference caused their reads to incorrectly align to *FRG1DP*, demonstrating the difficulty of investigating these genes with GRCh38 (**Fig. 5B**). Most of the new paralogs are found in other human genomes (**Fig. 5C**), and all paralogs except *FRG1KP2* and *FRG1KP3* have CpG islands overlapping the transcription start site with varying degrees of methylation and expression evidence in CHM13 (**Fig. 5D**, Table S16). This is just one example of the thousands of putative new paralogs uncovered by the T2T-CHM13 assembly, which we expect to drive future discovery in human genomic health and disease.

**Fig. 5.**
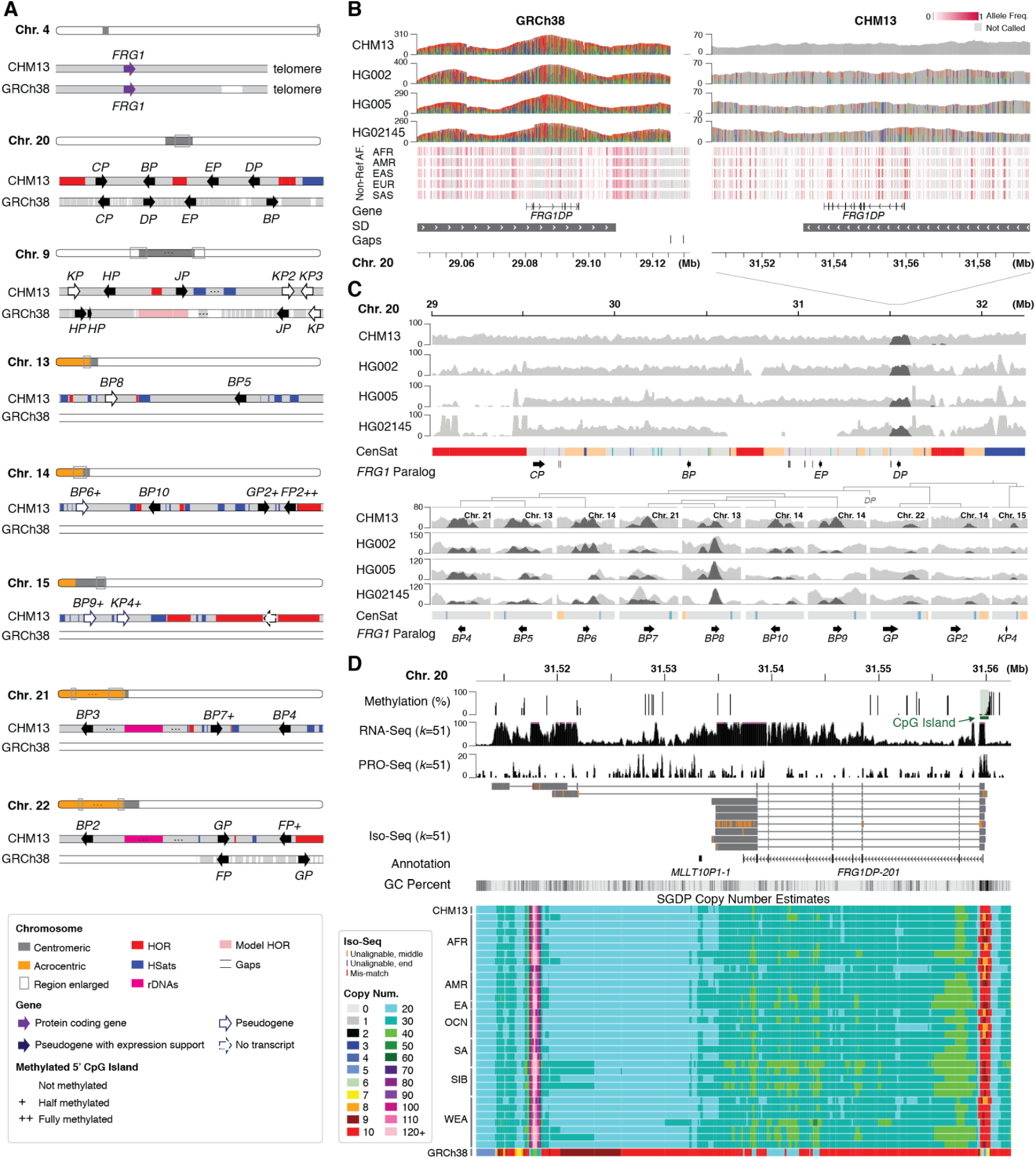
Newly resolved *FRG1* paralogs in the T2T-CHM13 assembly. **(A)** Diagram of the segmentally duplicated protein-coding gene *FRG1* and its 23 paralogs located on multiple chromosomes of CHM13. Paralog names are presented without the “*FRG1*” prefix for brevity and genes are drawn larger than their actual size. All paralogs were found near centromeric satellite arrays, either adjacent to centromeres or on the acrocentric p-arms, with most copies having expression support and CpG islands present at the 5’ start site with varying degrees of methylation (see legend). **(B)** HiFi read coverage and variants are shown for CHM13 and three other human samples mapped to the paralog *FRG1DP* in both GRCh38 and CHM13 references. When using GRCh38 as the reference, most of the region has excessive HiFi read coverage and non-reference variants, indicating that reads from the missing paralogs are mis-mapped to this *FRG1DP* (only variants with >80% coverage shown). When CHM13 is used as the reference, HiFi read coverage follows the expected 34–40x coverage, with no variants for CHM13 (as expected) and a typical heterozygous variation pattern for the other three samples (all variants >20% coverage shown). These non-reference alleles were also found in other populations from 1000 Genomes Project ILMN data, confirming them as true polymorphic variants. **(C)** Mapped HiFi read coverage (light gray) from the same four samples is also shown for other *FRG1* paralogs in CHM13, with an extended context shown for Chromosome 20. Coverage of all HiFi reads that mapped to *FRG1DP* in GRCh38 are highlighted (dark gray), showing the various paralogous copies they actually originate from (*FRG1BP4*–*10, FRG1GP, FRG1GP2*, and *FRG1KP4*). Background read coverage shows copy number variation within the surrounding regions, but with relatively even coverage across FRG1 paralogs, with the exception of *FRG1EP*, which appears to be absent from HG02145. Most other parlogs have diploid or haploid coverage levels, except *FRG1BP7* and *FRG1BP9*, which show higher coverage in some genomes, suggesting copy number polymorphism. **(D)** Methylation and expression results suggest transcription of *FRG1DP* in CHM13 (annotated as *FRG1DP-201*), with methylation depressed in the 5’ CpG island, a number of well-aligned ISO-Seq reads, and both RNA-Seq and PRO-Seq evidence (again filtered with 51-mer unique markers to prevent mis-mapping). In the copy number display, each *k*-mer of the CHM13 assembly is painted with a color representing the copy number of that *k*-mer sequence in a sample from the Simons Genome Diversity Panel (SGDP) (*85*). The CHM13 and GRCh38 tracks show the copy number of these same *k*-mers in the respective assemblies. The region surrounding *FRG1DP* shows an average of 20–40 (diploid) *k*-mer copies in CHM13, with a diploid copy number of 10 in the first exon (red) and a highly repetitive sequence towards the tail (purple/pink). This matches all samples from the SGDP, whereas GRCh38 looks entirely unlike any sample and systematically underrepresents the true copy number. All displayed browser tracks are available from the T2T-CHM13 UCSC Assembly Hub.

## Future of the human reference genome

The complete, telomere-to-telomere assembly of a human genome marks a new era of genomics where no region of the genome is beyond reach. Prior updates to the human reference genome have been incremental and the high cost of switching to a new assembly has outweighed the marginal gains for many researchers. In contrast, the T2T-CHM13 assembly presented here includes five entirely new chromosome arms and is the single largest addition of new content to the human genome in the past 20 years (**Fig. 1D**). This 8% of the genome has not been overlooked due to its lack of importance, but rather due to technological limitations. High accuracy long-read sequencing has finally removed this technological barrier, enabling comprehensive studies of genomic variation across the entire human genome. Such studies will necessarily require a complete and accurate human reference genome, ultimately driving adoption of the T2T-CHM13 assembly presented here.

One limitation of CHM13 is its lack of a Y chromosome. In order to finish a T2T reference sequence for all human chromosomes, we are in the process of sequencing and assembling the Y chromosome from the benchmark HG002 cell line, which has a 46,XY karyotype. The effectively haploid X and Y chromosomes can be assembled using the same methods developed here for the homozygous CHM13 genome. We have already sequenced, assembled, and validated the X chromosome from this line for use in the companion studies (*27, 76*) (Note S13, Figs. S37-39), and work on the HG002 Y chromosome is underway. Although much of the human Y chromosome is known to be heterochromatic and highly repetitive (*86*), we are confident that the strategy of combining HiFi and ONT reads used here will enable its completion (*12*).

Extending beyond the human reference genome, projects such as HapMap (*87*) and the 1000 Genomes Project (*30*) have been key to revealing genomic variation across human populations. However, technology limitations forced these early projects to focus mostly on a single reference genome and single-nucleotide variants. In contrast, long-read sequencing can recover the full spectrum of genomic variation including single-nucleotide variants, variable number short tandem repeats, and larger structural variants (*88*). Although short-read data cannot be confidently mapped to all regions of the human genome, reanalysis of 1000 Genomes short-read data (*50*) and copy number variation across a diverse set of human samples (*26*) has already shown that T2T-CHM13 constitutes a better reference even for short-read analyses. Improved computational methods and long-read omics assays are now needed to comprehensively survey polymorphic variation within the new regions of the genome assembled here, which we expect to reveal new phenotypic associations.

Highly accurate, long-read sequencing, combined with tailored algorithms, promises the *de novo* assembly of individual haplotypes and sequence-level resolution of complex structural variation. This will require the routine and complete *de novo* assembly of diploid human genomes, as planned by the Human Pangenome Reference Consortium (*89*). A collection of high-quality, complete reference haplotypes will transition the field away from a single linear reference and towards a reference pangenome that captures the full diversity of human genetic variation. Complete assembly of the CHM13 genome and our companion analyses have given only a small glimpse of the extensive structural variation that lies within the complete genome. Ideally, every genome could be assembled at the quality achieved here, since the small variants recovered by short-read resequencing approaches represent only a fraction of total genomic variation. However, taking the next step towards the telomere-to-telomere assembly of heterozygous diploid genomes, and further automating this process, presents a difficult challenge that will require the continued improvement of sequencing and assembly technologies. Until this ultimate goal is realized, and every genome can be completely sequenced without error, the T2T-CHM13 assembly represents a more complete, representative, and accurate reference than GRCh38, and we suggest it succeed GRCh38 in all studies requiring a linear reference sequence.

## Data and materials availability

*Sequencing data and assemblies (NCBI BioProject PRJNA559484):*

https://www.ncbi.nlm.nih.gov/bioproject/559484

*Sequencing data, assemblies, and other supporting data on AWS:*

https://github.com/marbl/CHM13

*Assembly issues and known heterozygous sites:*

https://github.com/marbl/CHM13-issues

*UCSC assembly hub browser:*

http://genome.ucsc.edu/cgi-bin/hgTracks?genome=t2t-chm13-v1.0&hubUrl=http://t2t.gi.ucsc.edu/chm13/hub/hub.txt

*Dotplot visualization and browser:*

https://resgen.io/paper-data/T2T-Nurk-et-al-2021/views/t2t-identity

*T2T Consortium homepage:*

https://sites.google.com/ucsc.edu/t2tworkinggroup

## Supplementary materials

### SupplementaryMaterials.pdf

- Notes S1–S13x
- Figs. S1–S39
- Tables S3-4, S13–S16

### SupplementaryTables.xlsx

- Tables S1, S2, S5–S12

## Acknowledgements

We would like to thank Mark Akeson, Andrew Carroll, Pi-Chuan Chang, Arthur Delcher, Maria Nattestad, and Mihai Pop for discussions on sequencing, assembly, and analysis; AnVIL, Amazon Web Services, DNAnexus, UW Genome Sciences IT Group, and the UConn Computational Biology Core for computational support; the NIH Intramural Sequencing Center, the UConn Center for Genome Innovation, and the Stowers Imaging Facility for experimental support. This work utilized the computational resources of the NIH HPC Biowulf cluster (https://hpc.nih.gov). Certain commercial equipment, instruments, or materials are identified to specify adequately experimental conditions or reported results. Such identification does not imply recommendation or endorsement by the National Institute of Standards and Technology, nor does it imply that the equipment, instruments, or materials identified are necessarily the best available for the purpose.

## Author contributions

*Analysis teams (leads*): Assembly:* SN*, SK*, MR*, MA, HC, CSC, RD, EG, MKi, MKo, HL, TM, EWM, IS, BPW, AW, AMP. *Acrocentrics:* AMP*, JLG*, MR, SEA, MB, RD, LGL, TP. *Validation:* AR*, AVB*, AM*, MA*, AMM*, WC, LGL, TD, GF, AF, KH, CJ, EDJ, DP, VAS, KS, YS, BAS, FTN, JT, JMDW, AMP. *Segmental duplications:* MRV*, EEE*, SN, SK, MD, PCD, AG, GAL, DP, CJS, DCS, MYD, WT, KHM, AMP. *Satellite annotation:* NA*, IAA*, KHM*, AVB, LU, TD, LGL, PAP, EIR, ASt, BAS, AMP. *Epigenetics:* AG*, WT*, SK, AR, MRV, NA, SJH, GAL, GVC, MCS, RJO, EEE, KHM, AMP. *Variants:* SA*, DCS*, SMY*, SZ*, RCM*, MYD*, JMZ*, MCS*, NFH, MKi, JM, DEM, NDO, JAR, FJS, KS, ASh, JW, CX, AMP. *Repeat annotation:* SJH*, RJO*, AG, PGSG, GAH, LGL, AFAS, JMS. *Gene annotation:* MD*, MH*, ASh*, SN, SK, PCD, ITF, SLS, FTN, AMP. *Browsers:* MD*, PK. *Data generation:* SJH, GGB, SYB, GVC, RSF, TAGL, IMH, MWH, MJ, JK, VVM, JCM, BP, PP, ACY, US, MYD, JLG, RJO, WT, EEE, KHM, AMP. *Computational resources:* CSC, AF, RJO, MCS, KHM, AMP. *Manuscript draft:* AMP. *Figures:* SK, SN, AMP, AR. *Editing:* AMP, SN, SK, AR, EEE, KHM, with the assistance of all authors. *Supplement:* SN, SK, with the assistance of the working groups. *Supervision:* RCM, MYD, IAA, JLG, RJO, WT, JMZ, MCS, EEE, KHM, AMP. *Conceptualization:* EEE, KHM, AMP.

