## Supplementary Materials for "The complete sequence of a human genome"

### “The complete sequence of a human genome” Supplementary Materials


**1. Molecular Methods** 3

Chromosome spreads and Fluorescent In-Situ Hybridization (FISH) 3

Estimating rDNA copy number from FISH images 3

**2. Ancestry analysis** 6

**3. Data Availability** 7

**4. Whole genome assembly** 8

Limitations of the current approach 8

HiFi graph construction 9

Read processing 9

Initial construction 9

Iterative graph simplification 10

Final graph overview 11

Manual graph resolution and gap filling 12

HiFi-only resolution 12

ONT-based tangle resolution 14

Resolution of Chr6 and Chr9 15

Consensus and polishing 15

Consensus 15

Gap filling 16

Polishing 16

**5. Assembly validation** 18

ddPCR copy number validation 18

Alignment-based validation 19

Strand-seq validation 19
Hi-C validation 21
BAC-based validation 22
K-mer based validation 24
Assembly completeness 25
**6. Technology-specific sequencing biases** 26
**7. Alpha-satellite and human satellite 1,2,3 validation** 34
**8. Resolution of the rDNA arrays** 40
**9. Genome analysis** 43
Assembly tracks 43
Annotation 43
**10. Acrocentrics and rDNA analysis** 48
**11. Mappability** 53
**12. FRG1 Analysis** 55
**13. HG002 ChrX Assembly and Validation** 59
Whole genome assembly 59
Molecular Methods 59
Pulsed field gel electrophoresis 59
Southern blotting 60
ddPCR 60
Alignment-based validation 61
**Bibliography** 63
**Additional files: SupplementaryTables.xlsx** Tables S1, S2, S5–S12

### 1. Molecular Methods

#### Chromosome spreads and Fluorescent In-Situ Hybridization (FISH)

For the preparation of chromosome spreads, cells were blocked in mitosis by the addition of Karyomax colcemid solution (0.1 µg/ml, Life Technologies) for 6-7h and collected by trypsinization. Collected cells were incubated in hypotonic 0.4% KCl solution for 12 min and pre-fixed by addition of Methanol:Acetic acid (3:1) fixative solution (1% total volume). Pre-fixed cells were collected by centrifugation and then fixed in Methanol:Acetic acid (3:1). Spreads were dropped on a glass slide and incubated at 65°C overnight. Before hybridization, slides were treated with 0.1mg/ml RNAse A (Qiagen) in 2xSSC for 45 minutes at 37°C and dehydrated in a 70%, 80%, and 100% ethanol series for 2 minutes each. Slides were denatured in 70% formamide/2X SSC solution pre-heated to 72°C for 1.5 min. Denaturation was stopped by immersing slides in 70%, 80%, and 100% ethanol series chilled to -20°C. Labeled DNA probes were denatured separately in a hybridization buffer by heating to 80°C for 10 minutes before applying to denatured slides. A fluorescently labeled probe for human rDNA (BAC clone RP11-450E20) was obtained from Empire Genomics (Williamsville, NY). Acrocentric chromosome paints were obtained from Oxford Gene Technology (Cambridge, UK). WaluSat DNA probe was PCR-amplified from CHM13 genomic DNA, labeled with biotin-dUTP by nick-translation reaction, and detected with fluorescently conjugated streptavidin post hybridization. Specimens were hybridized to the probe under a glass coverslip or HybriSlip hybridization cover (GRACE Biolabs) sealed with the rubber cement or Cytobond (SciGene) in a humidified chamber at C37°C for 48-72hours. After hybridization, slides were washed in 50% formamide/2X SSC 3 times for 5 minutes per wash at 45°C, then in 1x SSC solution at 45°C for 5 minutes twice and at room temperature once. Slides were mounted in Vectashield containing DAPI (Vector Laboratories).

#### Estimating rDNA copy number from FISH images

Wide-field images of chromosomal spreads were acquired on Nikon ECLIPSE Ti2 microscope equipped with Photometrics Prime 95B sCMOS camera and a 100x Plan Apo Lambda 1.45 NA objective. Acrocentric chromosome identities were assigned based on chromosome paints and morphological features. Individual rDNA loci were segmented via a selected threshold, and lines (at least 12 pixels wide) were created across each segmented rDNA locus to generate fluorescent intensity profiles. The averages of three ending values on both ends of each intensity profile were used to subtract the local background. The sum of background-subtracted intensity profile values represented the integrated intensity of each locus. The sum of all integrated intensities of all rDNA loci represented the total amount of rDNA per cell. The fraction of this total fluorescent intensity was calculated for each rDNA locus. The total rDNA copy number was estimated from Illumina sequencing data to be 405 copies per diploid genome. This number was multiplied by two as mitotic cells have a doubled genome (810 copies). The fraction of the total rDNA fluorescence intensity was used as a fraction of the total rDNA copy number to determine the number of copies on each identifiable chromosome and divided by two to get the final copy number per sister chromatid (Fig. S1). As CHM13 is homozygous, the rDNA copy numbers are similar on both haplotypes. Chromosomes 14 and 15 are exceptions, with the two haplotypes having a large difference in copy number. The shorter Chromosome 15 haplotype was confirmed by a single spanning ONT UL read (Fig. S2).

**A**

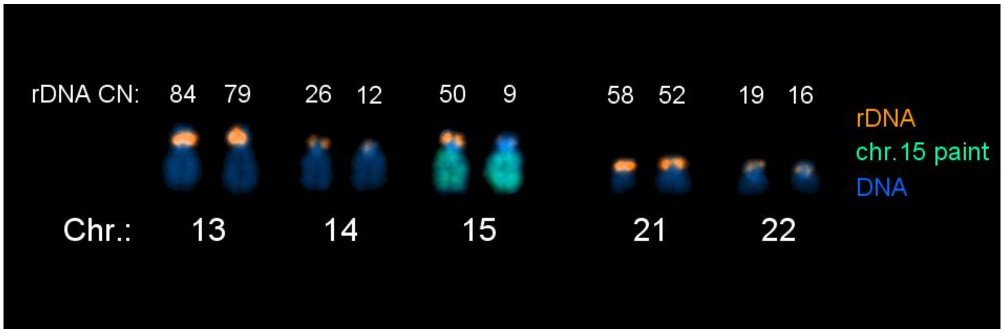


**B**

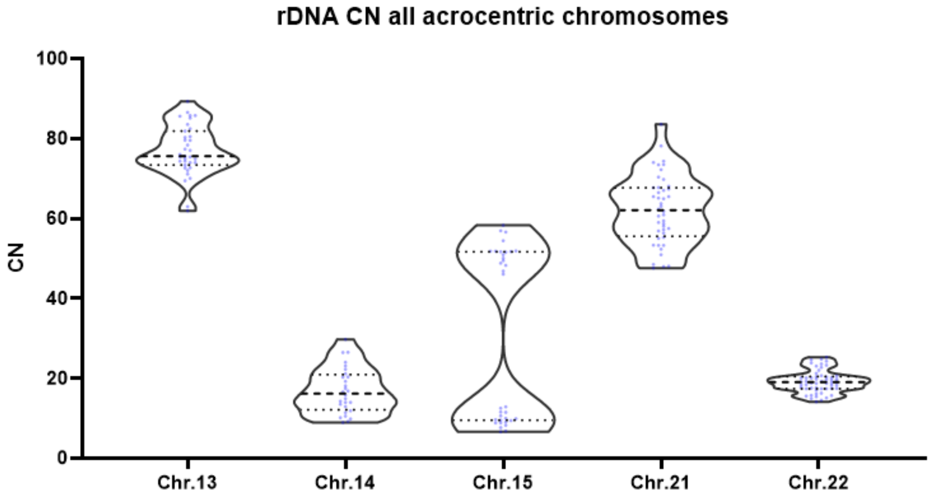


**Fig. S1: Estimation of rDNA copy numbers.** (A) Karyogram of acrocentric chromosomes from a CHM13 chromosome spread labeled by Fluorescent In-Situ Hybridization (FISH) with rDNA probe (orange) and Chromosome 15 paint (green). Chromosomes were counter-stained by DAPI. Chromosome 15 was identified by paint, other chromosomes were identified based on morphology. Estimated rDNA copy numbers per chromatids (rDNA CN) are shown on top of the acrocentric chromosomes with rDNA labeled in orange. Chromosomes 14 and 15 have larger differences in copy numbers, with the rest being within 5 copies. (B) Quantification of the rDNA copy number based on FISH. Chromosomal spreads from CHM13 cells were labeled with rDNA probe and acrocentric chromosome-specific paints. The rDNA copy numbers per specific chromatids were calculated from the fraction of the total fluorescent intensity of the rDNA signal on all chromosomes in a given spread and the Illumina sequencing estimate of the total copy number of rDNA repeats in the CHM13 genome. In addition to rDNA loci on painted chromosomes, all rDNA loci on other identifiable chromosomes were included.

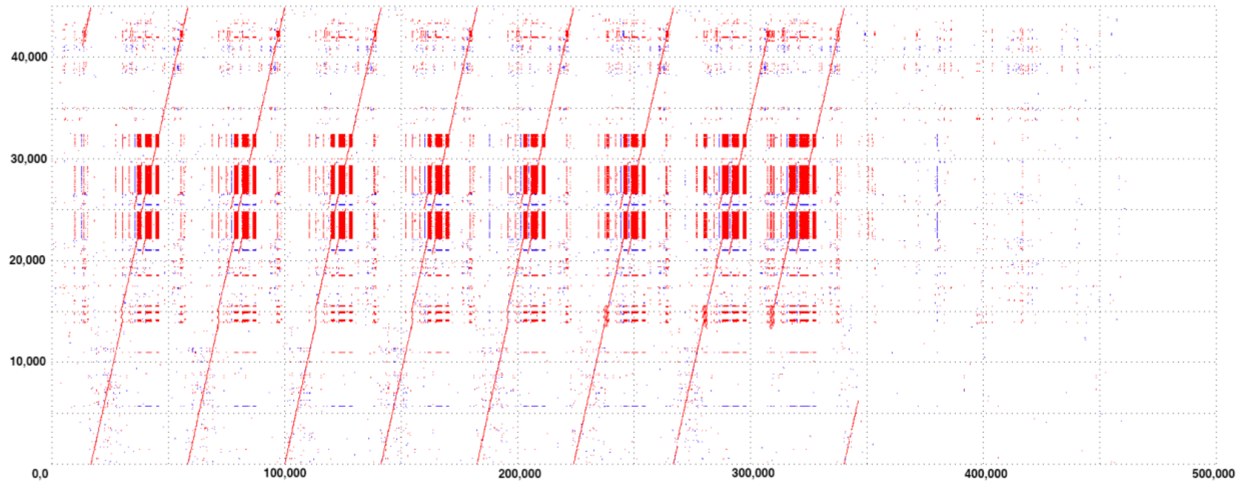


**Fig. S2: MUMmer** (*1*) **alignment of a canonical rDNA copy to a single ONT read.** The y-axis corresponds to the canonical rDNA unit (*2*) and the x-axis to the ONT read. Forward matches are in red and reverse complement matches in blue. The read has seven copies of the 45S gene cluster (0-15 kbp on y-axis) and shows a consistent structure within its repeat units. It is anchored in a non-rDNA sequence at the end (x-axis 350 kbp-450 kbp).

### 2. Ancestry analysis

We used RFMix v2.03-r0 (https://github.com/slowkoni/rfmix) to infer the local ancestry of the CHM13 genome (*3*). As a set of reference samples for ancestry, we used the 2,504 individuals sequenced in the Phase 3 release of the 1000 Genomes Project (*4*). Phased SNP genotypes for this reference set were generated by the 1000 Genomes Consortium through alignment and calling variants against the GRCh38 reference (*5*), and included only biallelic SNVs (http://ftp.1000genomes.ebi.ac.uk/vol1/ftp/data_collections/1000_genomes_project/release/20181203_biallelic_SNV/). We obtained a genetic map for GRCh38, originally generated on build 35 of the human reference genome by the HapMap Consortium (*6*), from Beagle (http://bochet.gcc.biostat.washington.edu/beagle/genetic_maps/).

CHM13 variants were called on GRCh38 with dipcall (https://github.com/lh3/dipcall) (*7*), through direct alignment of the CHM13 v1.0 assembly to the GRCh38 reference (*8*). For variable sites in the 1000 Genomes dataset that were not called as variants in CHM13, we assumed that CHM13 carried the reference allele. All CHM13 genotypes were represented as phased homozygous diploid genotypes (either reference/reference or alt/alt), which resulted in two sets of identical inferred local ancestry segments for each haplotype. We ran RFMix with default parameters (conditional random field spacing = 50 SNPs; random forest window size = 5 SNPs), grouping the 1000 Genomes reference panel into superpopulations (African, Ad Mixed American, East Asian, European, Southeast Asian). Since males are haploid for variants on the X chromosome, we included only female individuals in the reference panel for local ancestry inference for the X chromosome. Regions that overlapped with centromeres or were marked as inaccessible for the 1000 Genomes Project (http://ftp.1000genomes.ebi.ac.uk/vol1/ftp/data_collections/1000_genomes_project/working/20160622_genome_mask_GRCh38/) were excluded from results (UCSC Genome Browser; https://genome.ucsc.edu/cgi-bin/hgTables). The non-pseudoautosomal regions (PARs) of the X chromosome were inaccessible for this analysis due to the absence of variant calls outside the PARs. For identification of Neanderthal-introgressed haplotypes in CHM13, we used IBDmix (https://github.com/PrincetonUniversity/IBDmix), which detects introgressed archaic sequences in a set of individuals by identifying haplotypes that are identical-by-descent to a reference archaic sample (*9*). Of the three available Neanderthal individuals with high-coverage sequences, Vindija Neanderthal was chosen as the reference archaic individual due to its inferred closest relation to the Neanderthal population that admixed with modern humans (*10*). Local ancestry analysis indicated that the majority of the CHM13 genome is composed of European ancestries, we used European samples from the 1000 Genomes Project as the background set of modern individuals in addition to CHM13. These samples were sequenced to 30x coverage by the New York Genome Center, and variants were called on the GRCh38 reference (*11*). We required Neanderthal-introgressed haplotypes identified in CHM13 to have a LOD score of at least 4 and a length of at least 50 kb. We removed introgressed segments using bedtools within regions deemed inaccessible by the 1000 Genomes Project.

### 3. Data Availability

All data is available from NCBI GenBank (https://www.ncbi.nlm.nih.gov/assembly/) or NCBI Short Read Archive (https://www.ncbi.nlm.nih.gov/sra/) under the following accessions:

| **Data** | **NCBI Accession** |
| --- | --- |
| CHM13 sequencing data | SAMN03255769 |
| CHM13 T2T v1.0 assembly | GCA_009914755.2 |
| CHM13 T2T v1.1 assembly | GCA_009914755.3 |
| Hi-C | SRR13849427-SRR13849430 |
| 10XG Illumina | SRR10035390-SRR10035393 |
| PCRFree Illumina | SRX1009644 |
| ONT | SRR10035476-SRR10035573<br>SRR12564436-SRR12564459<br>SRR12618224-SRR12618325<br>SRR14000621-SRR14000622 |
| HiFi 10 kbp | SRR9087597-SRR9087600 |
| HiFi 20 kbp | SRR11292120-SRR11292123 |
| HG002 sequencing data | SAMN03283347 |
| HiFi 15+20 kbp | SRX7083054-SRX7083059 |
| HiFi 19kbp | https://s3-us-west-2.amazonaws.com/human-pangenomics/index.html?prefix=NHGRI_UCSC_panel/HG002/hpp_HG002_NA24385_son_v1/PacBio_HiFi/19kb/ |
| ONT | https://s3-us-west-2.amazonaws.com/miten-hg002/index.html?prefix=guppy_3.6.0/hg002/ |
| HG002 ChrX assembly | CP074113 |

### 4. Whole genome assembly

#### Limitations of the current approach

Our assembly string graph construction and tangle resolution procedures were specifically targeted to CHM13, which is largely homozygous. Thus, some of our design and heuristic decisions (e.g. arbitrary choice of haplotypes, bubble collapse) may not be applicable to assembling diploid genomes. Manual resolution would also be complicated by more heterozygous genomes due to shorter, more ambiguous nodes resulting from long homozygous regions with no single path traversing all nodes. Lastly, the basic string graph approach as described in Myers 2005 (*12*) has some fundamental deficiencies, arising from the initial removal of contained reads, making it a poor model for assembling datasets with varying read lengths, especially if the underlying genome is diploid/polyploid. See Fig. S3 for an example. These issues were largely avoided in this study due to the more or less uniform size of HiFi reads and the homozygous nature of CHM13.

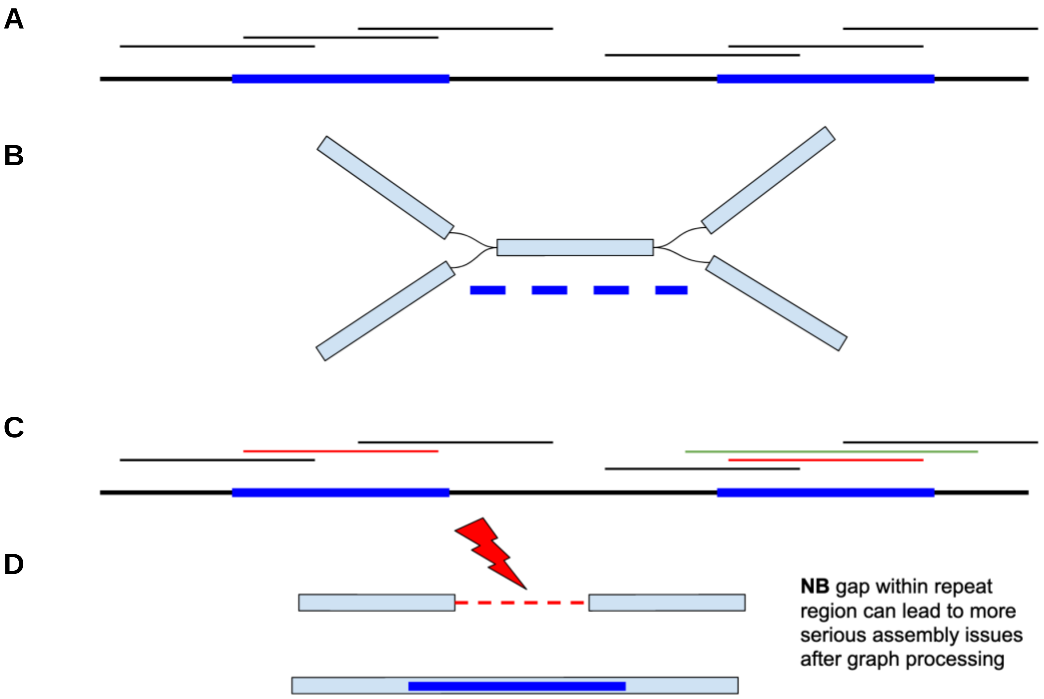


**Fig. S3: Fragmentation of a string graph constructed from reads of varying lengths.** (A) A genome with a two-copy repeat (shown in blue) sequenced by equal-length reads denoted by thin black lines. (B) Schematic bidirectional condensed string graph constructed from the input reads showing an unspanned repeat with branching. (C) Same genome sequenced by reads of varying lengths, the longer green read spanning one of the blue repeat copies contains reads originating from both repeat instances (red), leading to their exclusion from string graph construction. (D) The string graph constructed from this read set has a unitig spanning one of the repeat copies, while the second repeat copy is not represented by any continuous path. Note that more serious assembly issues might follow if the arising “dead-end” unitigs at the repeat boundary are further discarded by graph simplification procedures.

### HiFi graph construction

#### Read processing

HiFi 20 kbp libraries providing 32.4X average coverage of CHM13 (Data Availability) were used to construct a string graph. The graph construction started by applying slightly reshuffled modules of the HiCanu assembler to find and analyze overlaps between HiFi reads. For more details on individual steps see Nurk *et al.* (*13*). First, the reads were homopolymer-compressed. Then all >99% identity overlaps (dovetail and contained) longer than 500 bp were identified. Each read was subjected to a conservative correction based on the overlap pile-ups (see “Overlap Error Adjustment" section in (*13*)). Unlike in HiCanu, here we introduced the corrections to the homopolymer-compressed read sequences rather than only adjusting identity scores. We then recomputed overlaps (>500 bp and >99% identity) on the corrected reads and the overlap alignment identity was adjusted by masking differences in microsatellite array regions to account for previously observed increases in errors in these regions in HiFi reads (*13*). The need to apply and recompute overlaps stems from a limitation of HiCanu but is also common in other overlapping tools which limit the number of overlaps between a pair of reads. Currently, HiCanu stores at most two overlaps for every pair of reads (matching and inverted orientation) with higher identity overlaps preferred. In rare cases this can lead to the true overlap being dropped in favor of a (usually) shorter but higher identity overlap. Re-computing overlaps on corrected reads partially mitigated this issue. Additionally, HiCanu’s overlap search settings were adjusted (OvlMerThreshold=20000) to correctly identify overlaps within large repetitive arrays (e.g. centromere, rDNA). All overlaps with adjusted identity below 100% were then discarded. Finally, the BOGART module of (Hi)Canu was used to identify and discard reads with structural errors based on the analysis of the overlap pile-ups (*13*, *14*).

#### Initial construction

The resulting homopolymer-compressed reads and overlaps were then provided to a modified version of miniasm (*15*) to construct a Myers string graph (with a 1 kbp overlap threshold).

The modification fixed a problem in the pseudocode in Myers 2005 (*12*) (which has made its way into the miniasm implementation). The pseudocode was missing a final verification that the identified overlap is actually redundant with respect to the other two overlaps based on their sizes. Without this check, an overlap could be deemed transitive even if it implied an incompatible arrangement of the reads.

Notably, miniasm’s GFA format includes information about the read composition of the individual graph nodes corresponding to the unbranching paths (unitigs) in Myers’ string graph. Using this information, each node in the graph had a coverage estimate assigned to it to facilitate downstream analysis. Previous methods to assign coverage to string graphs (e.g. the A-stat (*16*)) are not applicable to shorter unitigs consisting of just a few reads. Thus, we implemented a novel approach. Each read is assigned a coverage value by considering the read’s overlaps and taking a minimum depth across all positions. Each graph node is assigned the median value of all the reads comprising it.

All further string graph processing within miniasm (e.g. graph simplification, tip removal, bubble removal and ‘relatively weak’ link pruning) was disabled.

#### Iterative graph simplification

We used custom graph simplification procedures to replace miniasm’s graph simplifications. This was implemented as a set of modular tools on top of public GFA libraries (*17–19*) available at https://github.com/snurk/sg_sandbox.

Graph processing steps:

- *Tip removal.* Dead-end nodes shorter than a specified threshold (set to 25 kbp excluding overlaps to an adjacent node) were removed. Initial rounds had a limit on the number of reads forming a tip that is iteratively increased. This minimizes the amount of correct sequence clipped at both the telomeres and “coverage gap” regions.
- *Superbubble collapsing.* Thresholds were set at 2 kbp on both the longest path and the maximal length difference between alternative paths. Each superbubble was replaced with the path having the maximum minimal overlap between its nodes. Note that the aggressive 2 kbp length difference threshold is a legacy of earlier stages of pipeline development. However, it did not affect the representation of the repetitive regions. Firstly, the limitations of the procedure prevent it from simplifying tangles of high complexity (e.g. most HSats). Secondly, early-processing-stage subgraphs were spot-checked around the nodes that were used multiple times within the traversals in an attempt to verify perfect repeat representation. A notable exception was the HSat3 region on Chromosome 9 where variation between genomic repeat copies was lost. This region spans a recent multi-megabase duplication, leading to chains of distinctive bubbles, representing variation between the two copies. During reconstruction of this region an alternate version of the HiFi string graph was built with the difference threshold set to 5 bp (see “Resolution of Chr6 and Chr9” section).
- *Low coverage node removal.* Removal of short nodes with assigned coverage value below an iteratively increased threshold (2-3-4-5X).
- *‘Weak’ edge pruning.* Edges corresponding to overlaps smaller than an iteratively increasing (2-4-6-8 kbp) threshold were removed. Before each threshold increase, other procedures (e.g. superbubble collapsing, tip removal etc.) were launched. Notably a handpicked absolute threshold was preferred over the miniasm strategy of pruning the overlaps based on their ratio against the size of the longest overlap for the same (side of the) node (read).
- *‘Simple’ bulge removal.* Removal of short nodes that have an alternative path of similar length (subject to additional conditions). This step is complementary to superbubble removal in tangled regions, where no clear superbubble subgraph can be identified.

- *Removal of some unusable edges.* An edge was considered unusable if it could not be a part of a traversal which uses all of the ‘unique’ and reliable (classified based on length and coverage) nodes (of a particular graph region) exactly once. At present, we only covered several cases forming distinct commonly occurring subgraphs, leaving the development of a more general procedure for future work.

The graph is compacted after every procedure by merging the non-branching paths in the graph into longer nodes. The progressive increase of the overlap threshold allows the avoidance of higher levels of repeat entanglement potentially arising from using a small overlap threshold, while leading to better preservation of continuity compared to using a single larger threshold. This strategy is similar in spirit to the progressive increase of the k-mer size, commonly used by de Bruijn graph based assemblers (*20*).

Based on subsequent chromosome reconstruction attempts, the resulting graph was manually adjusted in a few regions in order to restore some edges. In one case, Canu chose to store an overlap inconsistent with the graph, and this overlap was manually replaced with an alternate overlap (Canu can store only one overlap per read pair orientation). With the exception of the fragmentation caused by absent HiFi read coverage, and bubbles representing larger heterozygous / polymorphic indels, the resulting assembly graph had relatively few artifacts (e.g. redundant or non-genomic nodes, or missing edges). These artifacts reflected minor issues in our processing strategies and parameters as well as inherent deficiencies of string graphs built from reads of varying lengths (see ”Limitations of current approach”).

#### Final graph overview

To facilitate manual inspection of the graph, all graph nodes were aligned to GRCh38 (*21*) using mashmap (*22*) and assigned tentative chromosome labels (these alignments were not used to inform chromosome reconstruction). This showed that graph regions corresponding to different chromosomes are almost never connected by graph edges. Moreover, the connected components have a mostly linear structure, suggesting that the genome has few perfect repeats longer than 8 kbp (edges shorter than this are pruned during “Weak Edge Pruning”, above) on different chromosomes or at distant positions of the same chromosome. Edges connecting nodes of the acrocentric chromosomes were an exception, potentially pointing to recent recombination events across distal rDNA junction regions. Another exception is in the HSat3 region on Chromosome 9, which spans a recent multi-Mbp tandem duplication, consistent with the 9qh+ karyotype of CHM13 (Fig. S4), resulting in edges between the two copies.

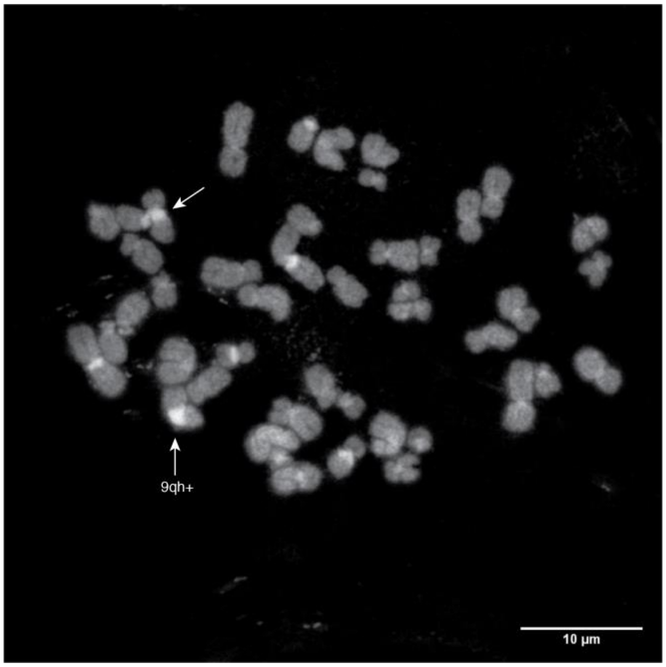


**Fig. S4: 9qh+ spread.** Micrograph of DAPI fluorescence from a mitotic chromosome spread from the CHM13 cell line at passage 10 (*23*). This spread was also labeled with multicolor chromosome paints for chromosome identification (not shown). Arrows denote the large pericentric Human Satellite 3 array on Chromosome 9. Bar, 10 µm.

### Manual graph resolution and gap filling

#### HiFi-only resolution

Reconstruction of some regions required the use of ONT UL reads (see”ONT-based tangle resolution”). In many cases, genomic paths through tangles could be unambiguously identified by analysis of the graph structure when considering estimated node multiplicities (based on coverage). This included cases where there was only a single traversal visiting every node (Fig. S5A) or the coverage information indicated the appropriate traversal (Fig. S5B). While handling bubbles representing heterozygous / polymorphic indels (see Fig. S5C,D for the example), our reconstruction preferred the longer alternative (unless its length was within 10 kbp of the shorter one and less than half the coverage). Somewhat surprisingly, this approach allowed us to obtain draft reconstructions for tangles consisting of dozens of nodes (Fig. S5C,D) without (or with very limited) usage of the ONT UL. We first considered resolutions of the most complicated tangles (e.g. those representing HSAT regions on Chromosomes 4 and 16) as low confidence, but subsequent ONT and HiFi based validation has mostly supported our initial resolutions.

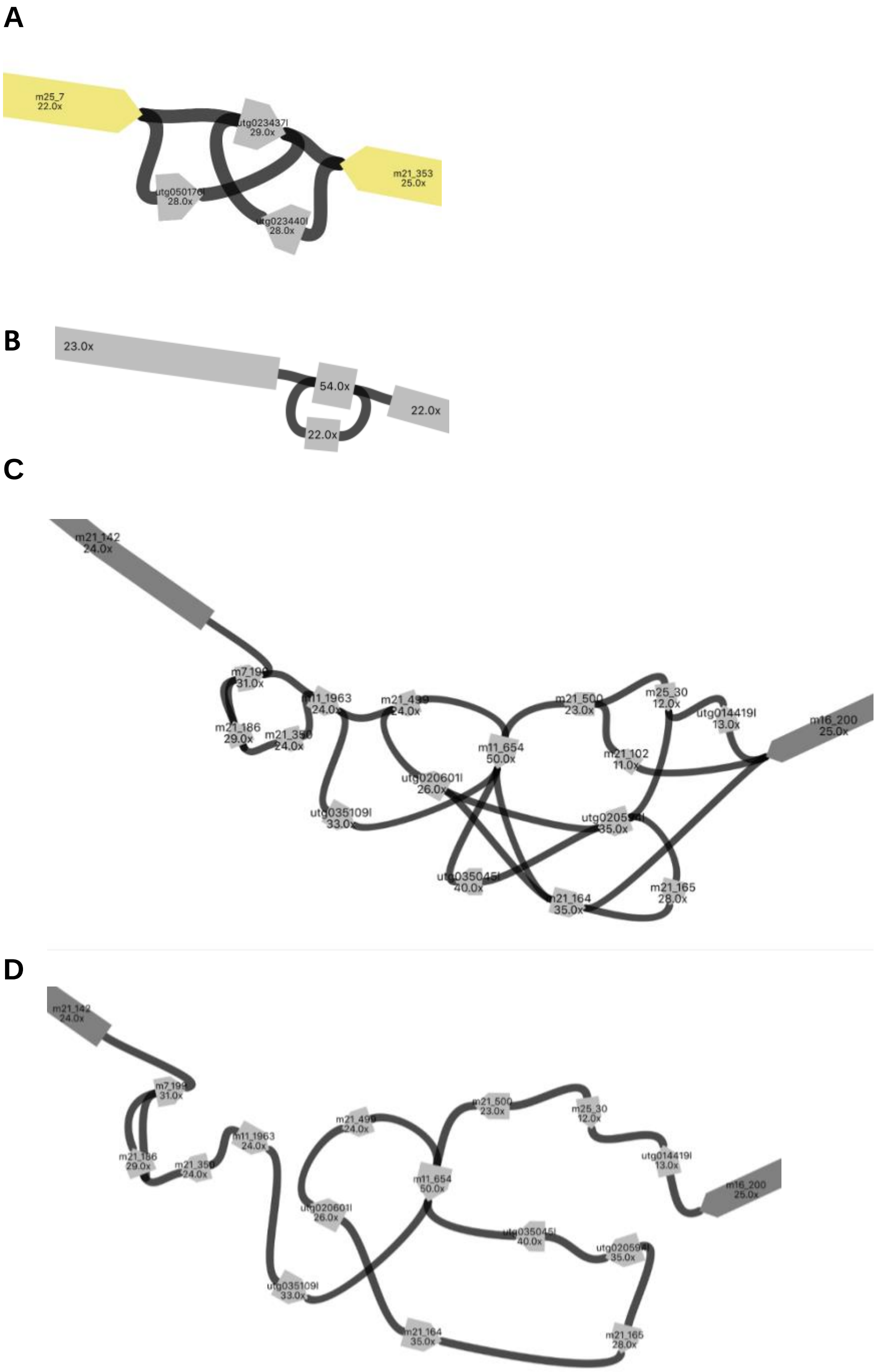


**Fig. S5: Examples of tangles resolved using Hamiltonian paths.** (A) There is only a single path that goes through all the nodes exactly once, as would be expected given their coverage relative to the large unique nodes. The path is m25_7+, utg050176l+, utg023437l-, utg023440l-, m21_353-. (B) Given the two-fold increase in the coverage of a single node, the genomic reconstruction should traverse the loop twice. (C) Subgraph corresponding to the AMY region on chr1. The original graph is shown without simplification from the automated graph construction pipeline. (D) We start by removing the heterozygous (4 bp insertion) variant represented by m21_102. We proceed with the iterative removal of edges that can not be a part of a genomic traversal assuming that it exists in the graph and visits all nodes with coverage < 40X only once. For example, note that the node utg035045l has a single incoming edge (leading from the node utg020594l) which we have to use for utg035045l to be a part of the traversal. Since utg020594l is expected to have multiplicity one, we can remove the other edge outgoing from it (leading to utg020601l). Repeated application of this strategy brings us to the final simplified subgraph, which has a single unambiguous traversal using node m11_654 (cov 50X) twice.

#### ONT-based tangle resolution

When graph information alone was insufficient, alignments of homopolymer-compressed Oxford Nanopore ultra-long (ONT UL) reads were used (*23*, *24*) to inform the reconstruction. Here, all available homopolymer-compressed ONT UL reads (longer 100 kbp) were aligned to the complete assembly graph using GraphAligner v1.0.12 (*25*). In some cases, these alignments were sufficient to provide reliable evidence for a particular path through the tangle. Unfortunately, GraphAligner's paths could not be used to reliably disambiguate traversal of more complex tangles. Moreover, due to GraphAligner limitations some of the highly tangled regions did not produce any alignments. When multiple alignment paths of similar overall score are present in the graph, GraphAligner has limited guarantees of picking the optimal one. In these cases, a more "brute-force" strategy was implemented that considered all candidate paths from the particular region of the graph. Here, ONT UL reads were aligned to every corresponding sequence using Winnomap v2.0 (*26*, *27*) and identity scores and break-points compared to identify reads exhibiting substantial differences in alignment quality between the candidate reconstructions. Alignment reports for these "informative" reads were then inspected to select the best-supported candidate path (Fig. S6).

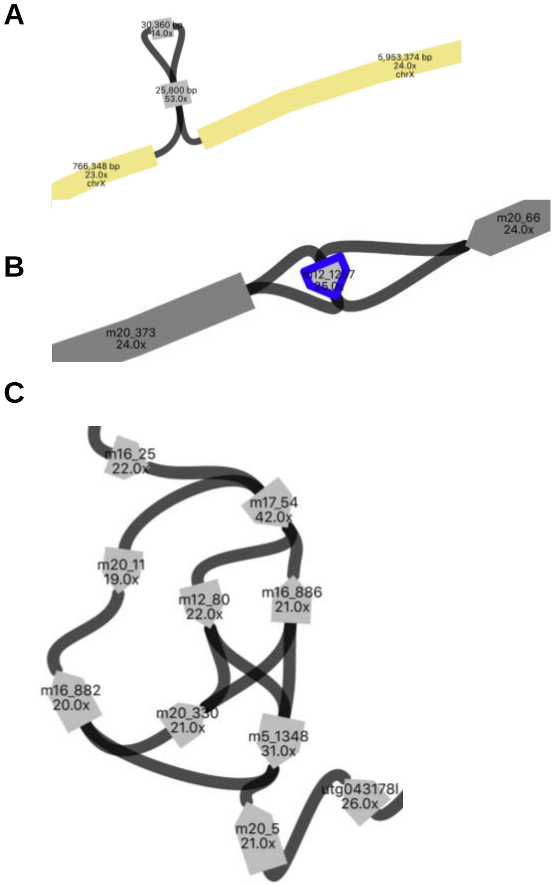


**Fig. S6: Examples of tangles resolved using ONT alignments.** (A) Example of a tangle that cannot be resolved based exclusively on the HiFi reads. The top node (14x) can be traversed in either the forward strand or the reverse strand. While such “hairpins” can potentially represent a perfect (reverse-complement) palindrome, in all the situations that we studied this was not the case, allowing us to identify the correct orientation of the node in the hairpin based on the sequence-to-graph alignments of ONT UL reads by GraphAligner (*25*). (B) Another example with an ambiguous orientation for the center node (m12_1277). (C) An example from Chromosome 4 where GraphAligner alignment paths were insufficient to identify the full traversal. The tangle was resolved by generating two candidate traversals (m16_25+, m17_54+, m12_80+, m20_330-, etc; m16_25+, m17_54+, m16_886-, m20_330-, etc) and analyzing Winnowmap (*26*, *27*) alignments of the ONT UL reads against each of the corresponding sequences.

##### Resolution of Chr6 and Chr9

Unfortunately, several regions in the graph could not be resolved using the methods outlined in the previous sections. For instance, the core of the rDNA arrays remained unresolved. Additionally, the Chromosome 6 centromere and Chromosome 9 Hsat3 array could not be easily resolved using the HiFi string graph and semi-manual analysis of ONT UL alignments. During Chromosome 6 centromere reconstruction, information about the higher-order repeat organization obtained by CentroFlye (*28*) from ONT UL data was used to inform the choice of graph traversal. For the reconstruction of the Chromosome 9 Hsat3 array (involving a large-scale recent duplication) MBG (*29*) was used to build a sparse de Bruijn graph from homopolymer-compressed HiFi reads. This graph was resolved by aligning the HiFi and ONT UL reads, identifying unique nodes based on their length and estimated coverage, and connecting them based on the read alignments. An alternate version of the HiFi string graph was built with the superbubble collapsing threshold set to 5 bp (see ‘Initial Construction’) to preserve minor differences between the duplicated sequences. This graph was then searched for a traversal corresponding to the draft sequence using GraphAligner.

#### Consensus and polishing

##### Consensus

After identifying the genomic graph traversals associated with a particular chromosome, their sequences were generated following the layout/consensus paradigm.

Throughout the analyses in previous sections, sequences were associated with graph nodes and paths (particularly for ONT UL read alignments). Prior to this point, they were obtained as simple concatenations of the homopolymer-compressed substrings of individual reads (joined according to the overlaps sizes). For the generation of the final consensus sequence, the graph traversal was translated into a backbone read layout—an orientation-aware concatenation of the chains of reads comprising each of the nodes along a path. Then, using procedures from the HiCanu assembler, the contained reads were incorporated into the layout, requiring 100% adjusted overlap identity, with the layout of the raw (uncompressed and uncorrected) read sequences being inferred as in (*13*). The consensus sequence was then computed via HiCanu’s POA-graph of the resulting layout (*13*, *14*). Upon validation of intermediate results using TandemTools (*30*), the logic and the parameters were adjusted to produce accurate layouts and consensus, even in highly repetitive genomic regions (in particular centromeres). A limitation of the Canu codebase, allowing a read to be used only once per layout, forced special handling of the nodes used more than once in the same traversal. The default strategy was to randomly distribute the reads comprising such nodes between their occurences. In cases where there were not enough (non-contained) reads for this strategy to produce a valid layout, the traversal was split at the duplicated node, consensus sequences produced for each piece, and finally merged.

GA-gaps were filled as in the 'Gap Filling' section. The per-base consensus accuracy (QV) was estimated to be Q70.2 by Merqury (*31*) based on the analysis of 21-mer spectra from the combination of HiFi and PCR-free Illumina reads (after filtering low-copy count 21-mers, Fig. S15).

#### Gap filling

Several chromosomes were represented by multiple disconnected components in the HiFi assembly graph, primarily due to the previously described (*13*) deficiency in HiFi sequencing, leading to depleted (or absent) HiFi coverage in (GA)-rich microsatellite regions. To mitigate the problem, ONT UL read alignments (aligned with winnowmap v1.1 (*26*, *27*)) were used to identify the graph unitigs flanking each coverage gap. After consensus sequences were generated for all recovered graph traversals (see above), gap 'patching' was performed using the previously published v0.7 assembly (*23*) as the source of the missing sequence. While gaps could have been filled *de novo* using spanning ONT UL reads, patching with v0.7 assembly sequence (based on the same ONT UL data) had the benefit of being extensively polished by other sequencing technologies. Patch coordinates were identified by aligning consensus sequences to the v0.7 assembly using minimap2 (*32*) with the command `minimap2 -H -x asm20`. Alignments were subject to strict criteria on size (> 4 kbp), identity (>98%) and number of allowed unaligned flanking bases (200bp). All 25 gaps across the genome were successfully patched, with a maximal patch size of 6.5 kbp.

#### Polishing

Given the high quality of this draft, a conservative polishing approach was taken. Vertebrate telomeric sequences (TTAGGG) were identified as in the Vertebrate Genome Project (*33*), identifying 45 of the expected 46 telomeres (22 chromosomes + X,  CHM13 has no Y chromosome). This highlighted that the telomere on the p-arm of Chromosome 18 was missing from this initial assembly. Telomeric reads were manually identified and added, increasing the assembly by 4,862 bp (*34*). Subsequently, short-reads were aligned to the assembly using bwa-mem v0.7.15 (*35*) and long-reads using Winnowmap2 v1.11 (*26*, *27*). Single-nucleotide variants (SNVs) were identified using the "hybrid" model in DeepVariant v1.0 (*36*), combining HiFi and short-read data. PEPPER-DeepVariant v0.3 (*37*) was used to identify SNVs from ONT UL data. Variants were filtered to eliminate poorly supported variants (VAF<=0.5, GQ<=30 for Illumina/CCS, and GQ<=25 for ONT) and this list of SNVs was further filtered using Merfin (*38*) which ensured the SNV change does not introduce *k*-mers not present in the Illumina or HiFi data. Larger structural variants (SVs) were identified using Parliament2 (*39*) for short-reads and using Sniffles (*40*) v1.0.12 from long reads (CLR, HiFi, ONT) with the command `sniffles -s 3 -d 500 -n -1 <input.bam>`. SVs with low support (<30% of reads supporting ALT allele) were removed (*34*). The technology-specific call sets were merged using Jasmine v1.0.2 (*41*) with the parameters `max_dist=500 min_seq_id=0.3 spec_reads=3 --output_genotypes`, and all variants >=30bp supported by at least two technologies were manually inspected using IGV (*42*). In total, 993 SNP and 3 SV corrections were made (*34*). The Merqury-estimated QV increased from Q70.2 to Q72.6. The initial polishing filtered SNPs near chromosome ends due to low coverage and ONT sequencing strand bias, leading to lower quality calls in telomeric sequence. Therefore, we ran a second round of polishing. To do so, we re-trained the existing model of PEPPER on HG002 Chromosome 20 by removing the forward strand reads entirely as the model internally is dependent on having reads from both strands (*34*). Using that model we generated a set of candidate variants in the telomere regions with PEPPER-DeepVariant v0.3. Next, we calculated the depth in the telomere regions using samtools (*43*). Finally, we used a custom script that takes the candidate variants and calculates the Levenshtein distance between the canonical telomere *k*-mer and the sequence we derive after applying the candidate variant. If the candidate had minimum allele frequency of 0.5, minimum genotype quality of 2, and reduced the Levenshtein distance to the canonical telomere *k*-mer compared to the existing sequence, we accepted the edit. We also trimmed the sequences in the telomere to remove regions where we observed a read depth lower than five. Using this method we made 454 telomere edits in the v1.0 assembly (*34*). The Merqury-estimated QV increased further to Q73.9 (67.86 from PCR-Free Illumina-only; 69.80 from HiFi-only).

### 5. Assembly validation

Our assembly strategy was iteratively refined by validation of intermediate results, most notably by TandemTools (*30*). We identified deficiencies in the pipeline (e.g. missing high-coverage overlaps and low consensus quality, 'Read Processing' and 'Consensus' sections) which were corrected before building the final chromosome reconstructions. We validated the final assembly using a mixture of techniques and sequencing data, as detailed below. Remaining known issues have been catalogued and are available at https://github.com/marbl/CHM13-issues.

#### ddPCR copy number validation

ddPCR reactions were performed using unique primers designed for amplicons in targeted regions (Table S1: "ddPCR array assays", S2:"ddPCR gene assays").

Each reaction consists of 10 uL 2x ddPCR QX200 Evagreen Supermix, 0.2 uL of restriction enzyme for fragmentation, 1 uL 10 uM primer mix, 1 uL of 0.1-1 ng CHM13 DNA template and 7.8 uL with nuclease free water. Mastermixes were then emulsified with Evagreen droplet generator oil (Bio-Rad) using a QX200 droplet generator according to the manufacturer's instructions. After droplet generation, thermocycling was performed with the following parameters: 10 min at 95°C, 40 cycles consisting of a 30-s denaturation at 94°C and a 60-s extension at 59°C, followed by 10 min at 98°C and a hold at 4 °C. Control reactions without the DNA were performed to rule out non-specific amplification.

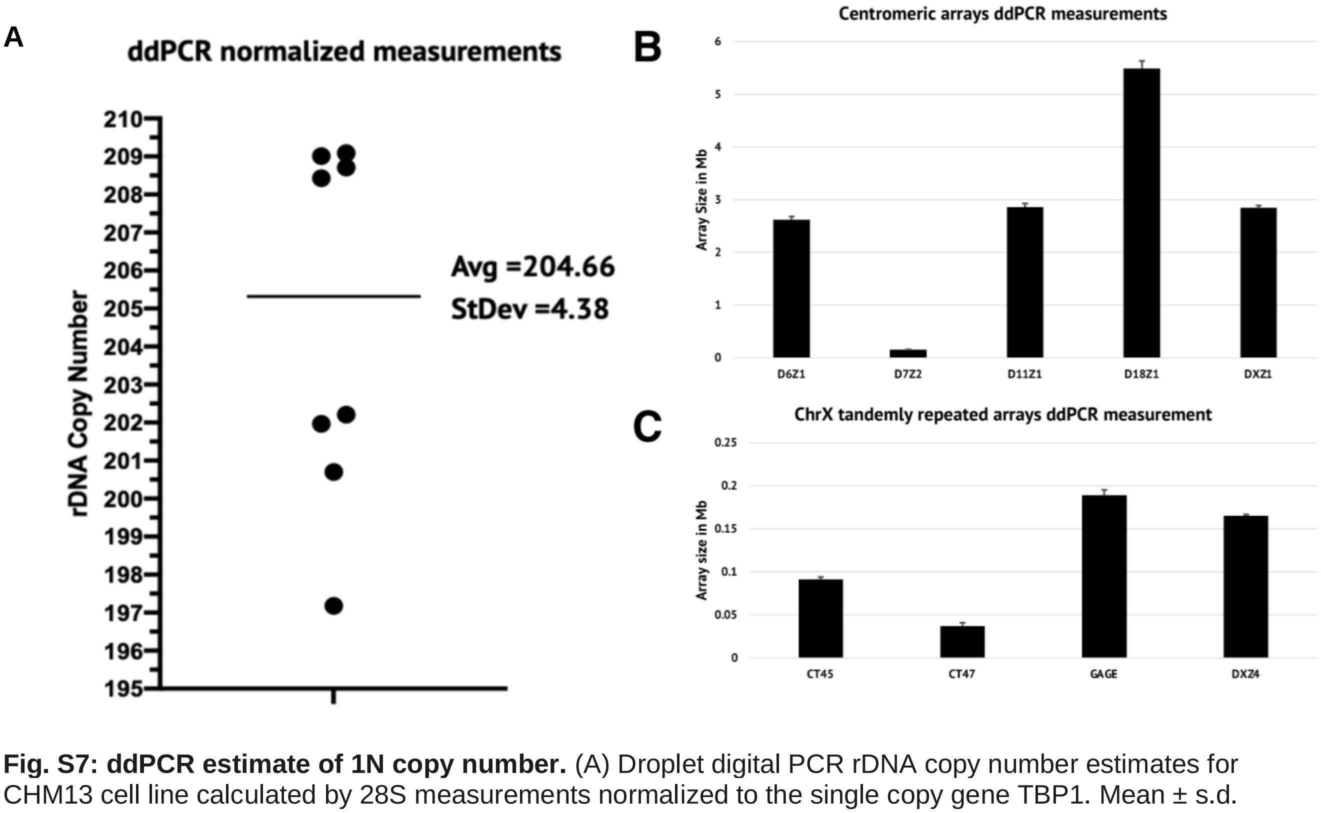


**Fig. S7: ddPCR estimate of 1N copy number.** (A) Droplet digital PCR rDNA copy number estimates for CHM13 cell line calculated by 28S measurements normalized to the single copy gene TBP1. Mean ± s.d. is shown. The estimate is consistent with the FISH estimates in Fig. S1. (B) Higher-order repeat (HOR) sizing from ddPCR. All estimates are consistent with annotated (*44*) sizes in the assembly (D6Z1 =

S1C6H1L = 2.77 Mbp; D7Z2 = S5C7H2 = 0.20 Mbp; D11Z1 = S3C11H1L = 3.38 Mbp; D18Z1 = S2C18H1L = 4.97 Mbp; DXZ1 = S3CXH1L = 3.11 Mbp). (C) Tandem repeat sizes on Chromosome X from ddPCR. All estimates are consistent with the previously published (*23*) and current reconstruction (CT45 = 0.15 Mbp; CT47 = 0.04 Mbp; GAGE = 0.19 Mbp; DXZ4 = 0.17 Mbp).

Following PCR amplification, the 96-well plate was transferred to a QX200 droplet reader (Bio-Rad). Positive droplets were automatically determined by the QuantaSoft software. Concentrations reported where copies/µL of the final ddPCR reaction were adjusted according to the TBP1 single copy gene present at Chromosome 6. The normalized copy number values for 28S repeated array were calculated as follows: [(28S target copies/µL)/(TBP1 copies/µL)] x 10 (Fig. S7A). The normalized copies for other arrays were calculated as for 28S, using the single-copy gene listed in Table S1.

#### Alignment-based validation

We mapped all available data (HiFi, ONT) to the assembly using Winnowmap v2.01 (*26*, *27*) and PCR-free Illumina data using bwa v0.7.15 (*35*). PCR duplicate-like redundancies were removed using biobambam2 `bamsormadup` (v2.0.87) (*45*) with default parameters. For long-read analysis, non-primary alignments were filtered using `samtools view -F 256` (*43*). The resulting read coverage was evenly distributed across all technologies, with the notable exception of some human satellite classes (see 'Technology-specific sequencing biases' for details). We used these alignments to generate summary statistics (min, max, mean, median) for 10 kbp non-overlapping windows of the assembly for each of: alignment length, read length, identity, and MAPQ.

#### Strand-seq validation

We validated the assembly using Strand-seq (*46*, *47*) data from CHM13 (*24*, *48*) and compared it to GRCh38. We first aligned paired-end Strand-seq data to both CHM13 and GRCh38 assemblies using BWA-MEM (version 0.7.15). Next we used breakpointR (*49*) to detect regions with recurrent changes in strand directionality across multiple Strand-seq libraries. Regions with the majority of reads mapped in minus orientation are suggestive of unresolved inversion or contig misorientation, while regions where we see a mixture of plus and minus reads are suggestive of possibly collapsed or low mappability (*50*). Overall we observe no large inversion or misorientation (Fig. S8A) as well as no chromosome mis-assignment (Fig. S8B) events in CHM13 assembly.

A

B

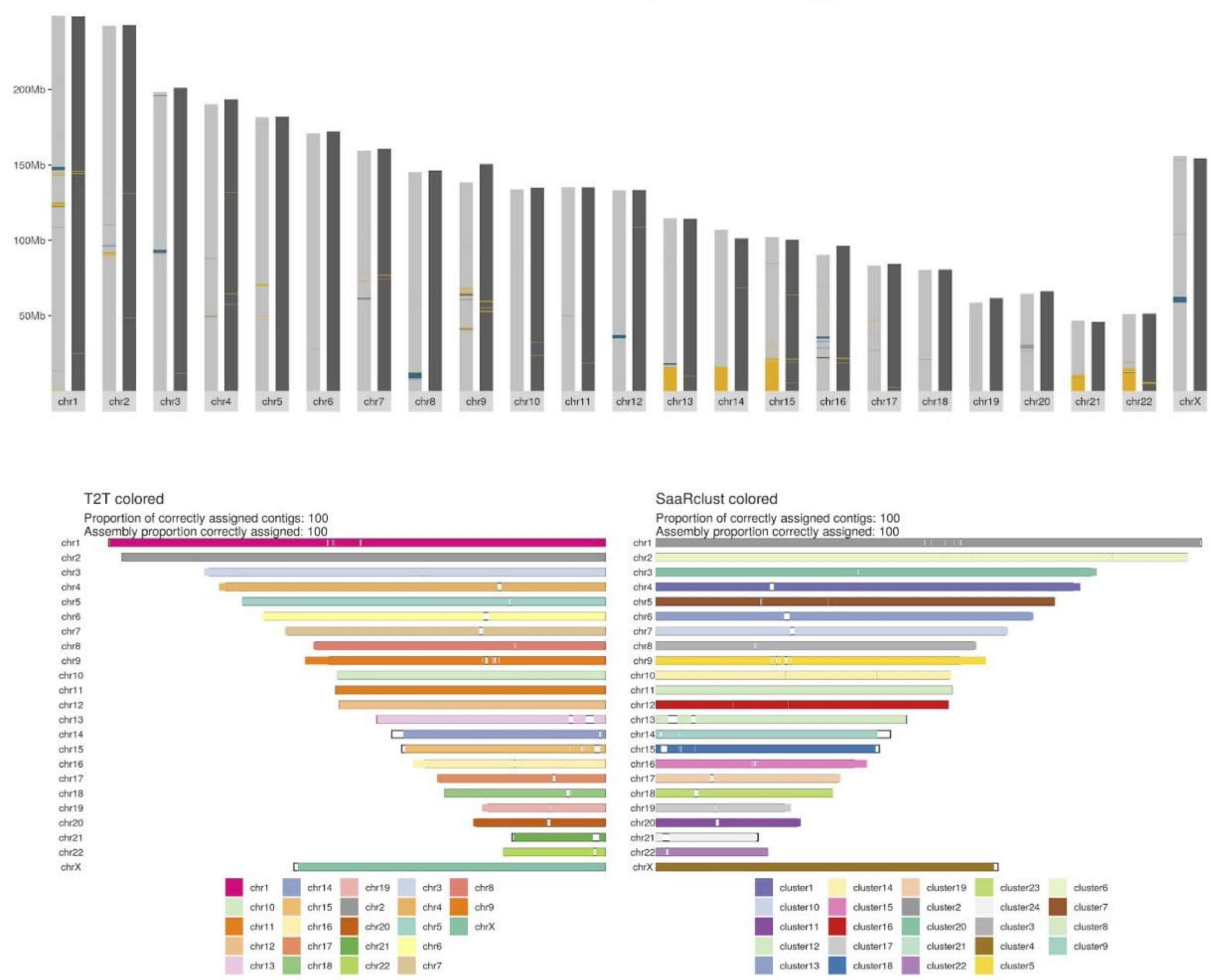


**Fig. S8: Strand-seq supports correct orientation and chromosome assignments.** (A) Comparison of GRCh38 and a preliminary version of the CHM13 assembly. GRCh38 is plotted as light gray bars and CHM13 as dark gray bars. Changes in assembly directionality (shown in blue) might point to an unresolved inversion or misoriented contigs. Other highlighted assembly features are low mappability (or possibly collapsed) regions (shown in yellow). There are few inverted regions in CHM13 in contrast with GRCh38 (0.0002 Mbp vs 28.41 Mbp). GRCh38 has a high count of low mappability regions, including the short arms of all acrocentric chromosomes which correspond to large gaps. In contrast, in CHM13, the low mappability regions are reduced to 8.99 Mbp from 101.10 Mbp in GRCh38. (B) Ideogram shows assembled contigs aligned to CHM13 colored based on cluster identity determined by SaaRclust (*50*, *51*). In an ideal scenario there is a single color per chromosome. The left ideogram is colored by contig names in the assembly. As expected, there is a single color per chromosome. Right ideogram is colored based on the cluster ID assigned to each 200 kbp window in a contig. In case of large scale chromosomal mis-joins, we expect to see two or more colors assigned to a contig. None of the CHM13 contigs show this, all being consistent with a single chromosome bin.

The Strand-seq alignments allowed us to verify chromosome assignments even in repetitive regions, such as the acrocentric chromosomes. Fig. S9 shows the mapping of strand-seq reads across Chromosome 15. In contrast to GRCh38, which has no reads mapped to the short arm,

CHM13 shows a consistent read orientation across the entire chromosome, confirming the coloring in Fig. S8B and the correctness of the acrocentric reconstructions.

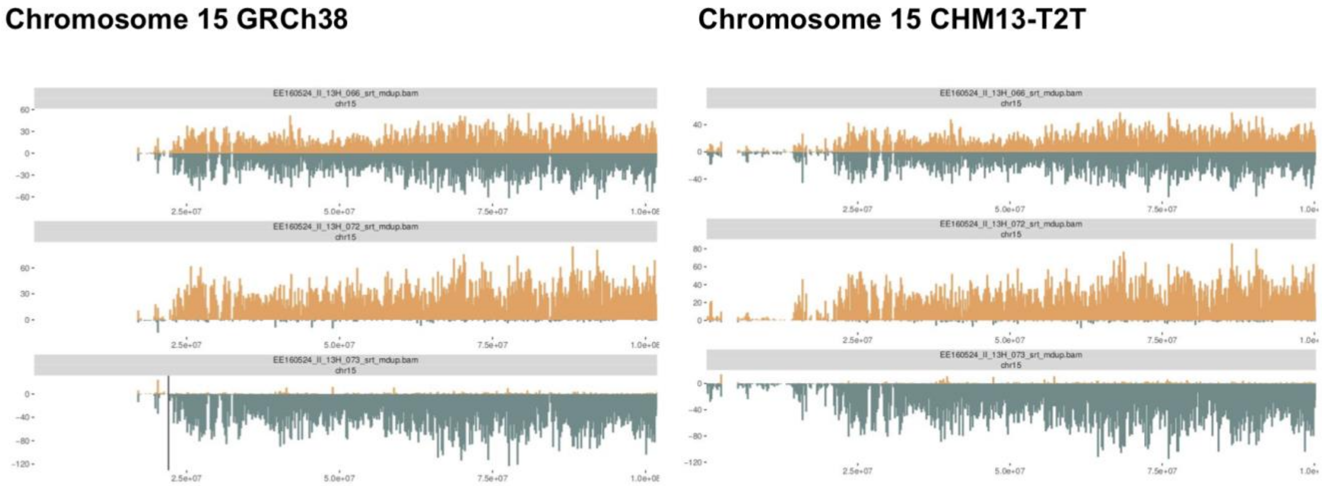


**Fig. S9: Mapping of short Strand-seq reads in rDNA loci of Chromosome 15.** We visualize the mapping of Strand-seq reads to Chromosome 15 of GRCh38 (left) and CHM13 (right). For the same three Strand-seq libraries we count Strand-seq reads mapping to either plus strand (teal) or minus strand (orange) of the reference genome (GRCh38 or CHM13) in 200 kbp bins reported as vertical bars (plus - below zero; minus - above zero). Only reads with MAPQ >10 are plotted. As expected, there are no reads mapping to GRCh38 from 0-15 Mbp. In contrast, there are enough reads mapping to CHM13 to confirm the orientation and assignment of the short arm of this chromosome.

#### Hi-C validation

We used Hi-C data to validate the long-range structure of the assembly. Hi-C reads were aligned to the CHM13v1.0 assembly using the VGP Hi-C mapping and visualization pipeline (*33*). In brief, both ends of a read pair were mapped independently using BWA-MEM (*35*) with the parameter `-B8`, and filtered when mapping quality was <10. Chimeric reads were trimmed from the restriction site onward, leaving only the 5′ end. The filtered single-read alignments were then rejoined as paired read alignments. Alignments were converted with PretextMap (https://github.com/wtsi-hpag/PretextMap) and visualized with PretextView (https://github.com/wtsi-hpag/PretextView) The resulting Hi-C interaction matrix showed a clean assembly, with strong interactions within chromosomes and few strong interactions between chromosomes (Fig. S10), supporting the overall structure of the assembly.

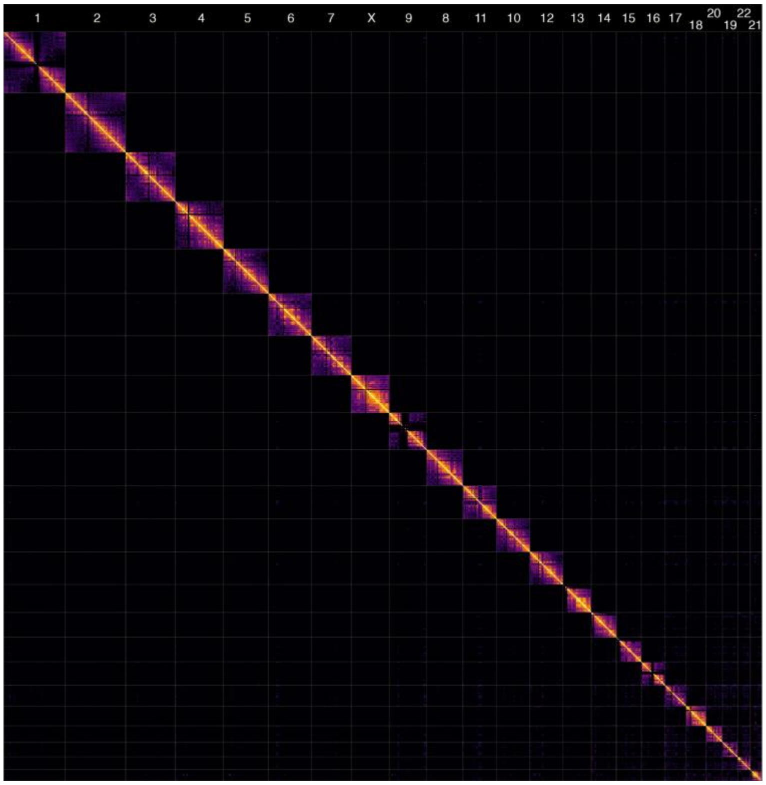


**Fig. S10: Hi-C interaction matrix visualized with PretextView (https://github.com/wtsi-hpag/PretextView).** There are strong signals within chromosomes. There is no off-diagonal or no chross-chromosomal signal indicating the chromosomes are complete and not chimeric. Black regions within chromosomes (e.g. Chromosomes 1 or 9) indicate repetitive regions where no short-reads could be confidently anchored. The short arms of the acrocentric chromosomes (13, 14, 15, 21, 22) show sufficient signal to confirm the correct assignment of each short arm to the chromosome.

#### BAC-based validation

We validated the assembly using previously sequenced and newly released BACs for CHM13 (*24*, *48*), including corrections made by HiCanu (*13*, *52*). 644 out of 647 BAC sequences had a continuous mapping to the assembly at Q41.94 (averaged across all aligned bases). To confirm the correctness of the assembly in the regions corresponding to the three remaining BACs, we selected all ONT UL reads with primary alignments to Chromosome 17 and Chromosome X of CHM v1.0 assembly at the locations of the partial BAC alignments. We also included a "resolved" BAC (AC279712.1) from Chromosome X as a control. Selected ONT reads were then aligned to the BACs in question with Winnowmap v2.0.1 (the command `winnowmap -z150 -t 16 -ax map-ont -H -k 19`, retaining only primary alignments (`samtools view -F 256`). Alignments were inspected in IGV. The three "unresolved" BACs (AC279506.1, AC279581.1, AC279712.1) show uneven coverage and an increase in variant positions, indicating mismatches versus the ONT data (Fig. S11-13). In contrast, the control BAC exhibits even coverage and no variant sites (Fig. S14). As the assembly did not use ONT data for consensus, this provides independent confirmation of its structure. It is likely that the relatively low QV (compared to the Merqury estimated values) is due to limitations of BAC quality as opposed to the assembly.

A

B

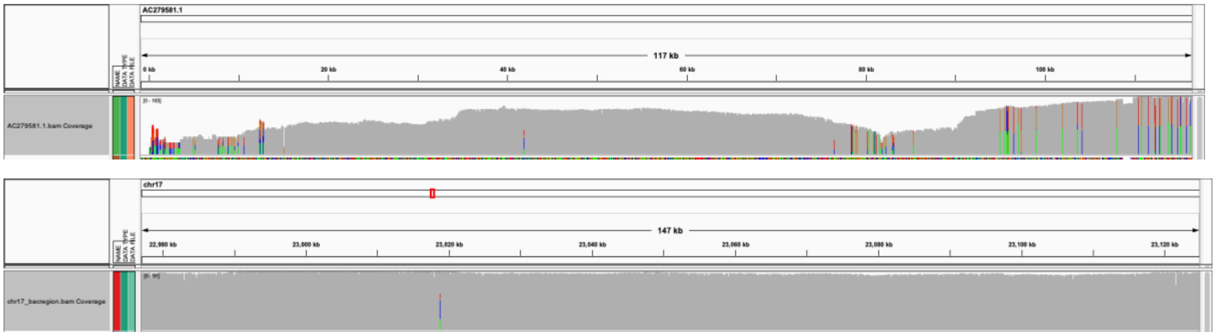

**Fig. S11: Concordance of ONT reads with BAC AC279506.1 and corresponding assembly region.** This 180,858 bp long BAC aligned incompletely, breaking at 20 kbp (BAC:19,686-180,858 aligned to chr17:22,937,576-23,096,790 in rev strand). (A) Alignments for the first 100 kbp of the BAC show low coverage at the start (before 20 kbp) with an increase in variant positions. There is also low coverage in the BAC at 80 kbp. (B) In contrast, the corresponding position in the assembly (chr17:22,900,000-23,080,000) shows no such coverage dropouts and no increase in variant calls.

A

B

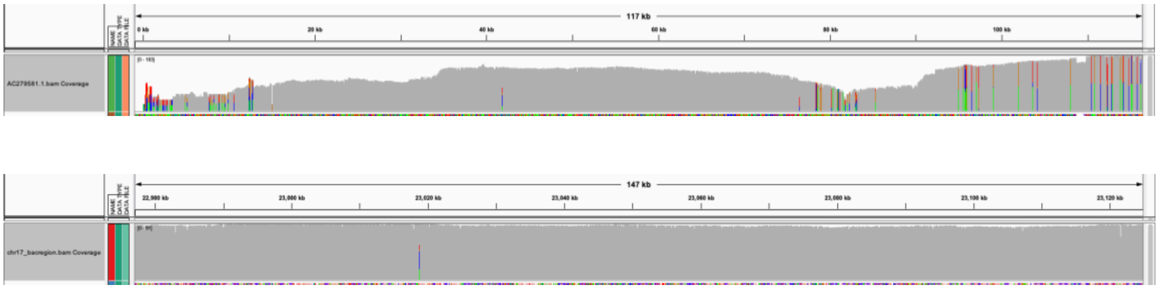

**Fig. S12: Concordance of ONT reads with BAC AC279581 and corresponding assembly region.** This 117,950 bp long BAC aligned incompletely, breaking at 94 kbp (BAC:25-93,747 to chr17:22,9770,000-23,070,901 in fwd strand). (A) Alignments for the BAC show low coverage at the start with another drop in at 80-90 kbp and an increase in variant positions after 90 kbp. (B) In contrast, the corresponding position in the assembly (chr17:22,980,000-23,120,000) shows no such coverage dropouts and no increase in variant calls.

A

B

C

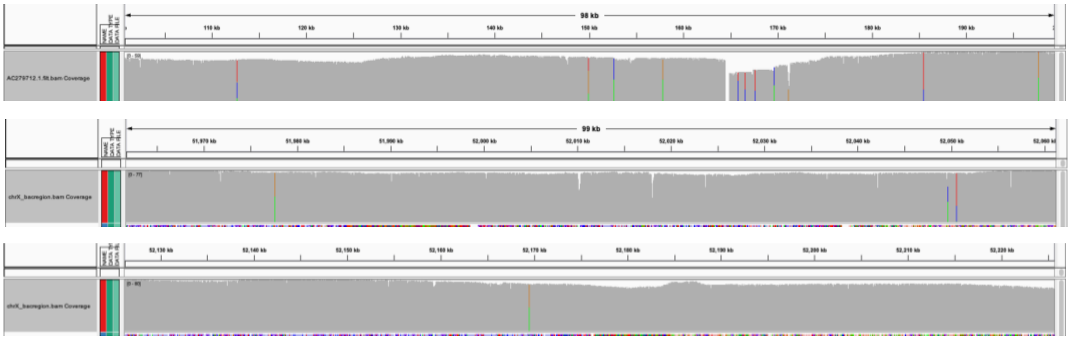

**Fig. S13: Concordance of ONT reads with BAC AC279712 and corresponding assembly regions.** This 329,285 bp long BAC aligned in two pieces to the same location in the assembly (BAC:4-164,475 and BAC:164,810-329,291 aligned to chrX:52,011,667-52,176,139 in fwd strand). This indicates a potential repeat collapse in the assembly. (A) Alignments for the 110-190 kbp region of the BAC show a dropout in coverage at 165 kbp, corresponding to the break between the two alignments. (B) The start position of the alignment in the assembly (chrX:51,970,000-52,060,000) shows no such coverage dropouts and no increase in variant calls. (C) The end position of the alignment in the assembly (chrX:52,130,000-52,220,000) shows even coverage and no variant artifacts to indicate a repeat collapse.

**A**

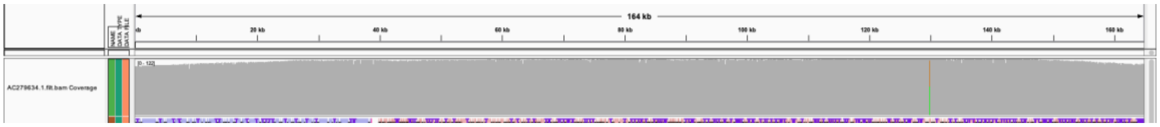


**B**

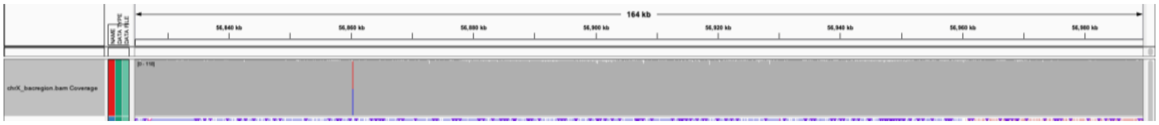


**Fig. S14: Concordance of ONT reads with BAC AC279634 and corresponding assembly region.** This 165,682 bp long BAC aligned in one piece (BAC:0-165,682 to chrX:56,824,611-56,990,290 in rev strand). (A) Alignments for the BAC show even coverage and no variant positions. (B) The corresponding position in the assembly (chrX:56,830,000-56,980,000) shows even coverage and no increase in variant calls.

### K-mer based validation

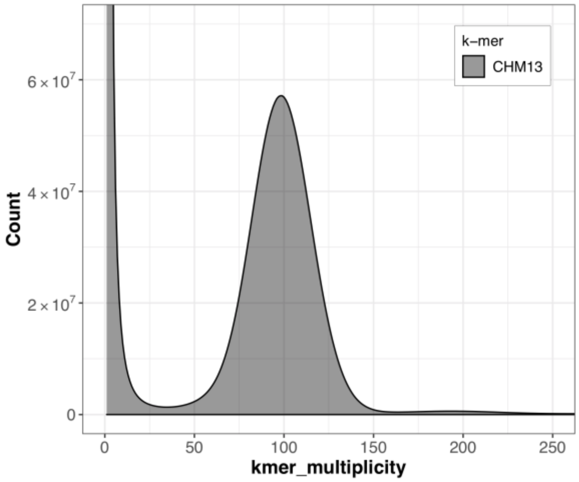


**Fig. S15: Distribution of k-mer counts from PCR-free Illumina data.** *k*-mer multiplicities were computed with meryl (*31*) and plotted with merqury (*31*).

Merfin (*38*) analysis confirmed that assembly k-mer multiplicity is consistent with both the HiFi and PCR-free (Fig. S15) sequencing data (Fig. S16), with a notable improvement from the v1.0 to the v1.1 assembly due to the filled in rDNA arrays.

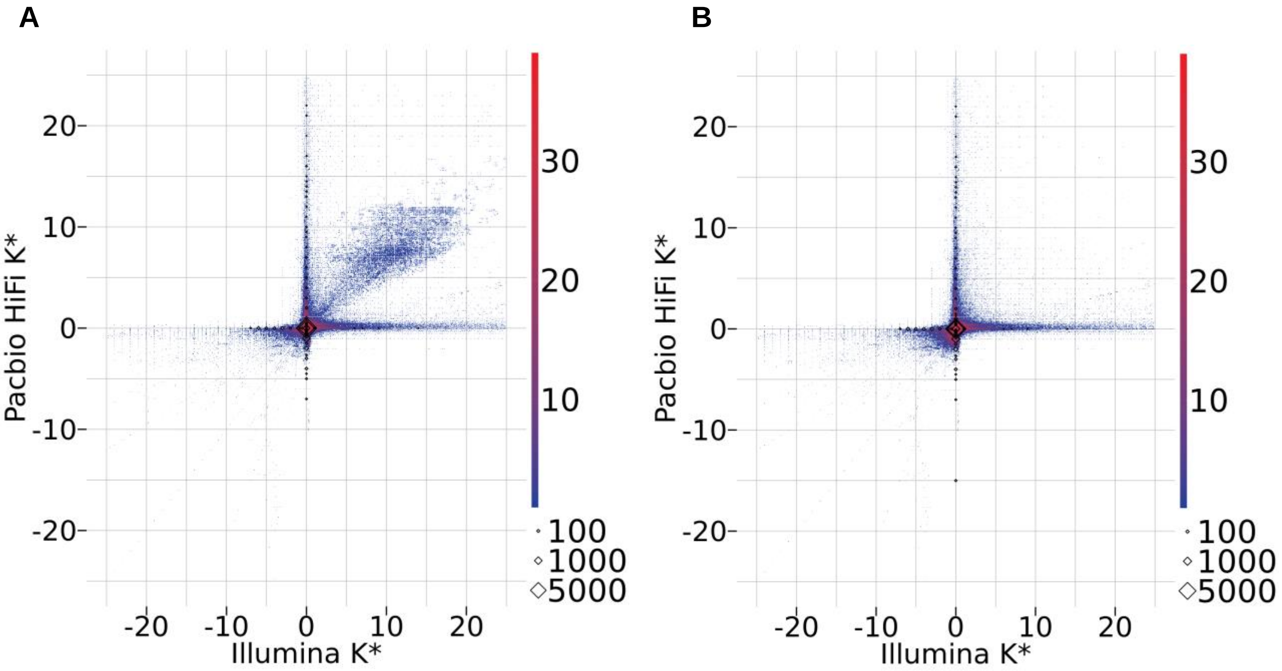


**Fig. S16: Merfin k* statistics showing assembly concordance with HiFi and PCR-free sequencing data.** Positive values indicate the k-mers have a higher copy in the sequencing data than the assembly (collapse in the assembly) while negative values indicate a lower copy in the sequencing data (expansion in the assembly). (A) *k*-mer multiplicities within v1.0 are mostly consistent with HiFi and PCR-free data (k* close to 0), with a cloud of *k*-mers collapsed in the assembly. (B) corresponding plot for the v1.1 assembly indicates that the collapsed regions in the assembly have been resolved.

#### Assembly completeness

Assembly completeness was evaluated using both using *k*-mer statistics and mapping-based statistics. Using PCR-free Illumina and HiFi data, Merqury (*31*) completeness for the assembly was 99.1% and 99.8%, respectively. Based on primary HiFi alignments, 97.5% of the reads (97.5% of bases) have an alignment with a MAPQ=60, >= 99% identity, and covering >= 90% of the read length. In contrast, for CHM13 v0.7 (*23*), only 92.7% of reads (92.6% of bases) meet this criteria. Some reads were expected not to map full length or at high identity due to heterozygous variants present in the CHM13 cell line but not in the assembly. Specific regions such as simple sequence repeats in HiFi data also show an elevated error rate (*13*). Thus, the criteria was relaxed to include reads with an alignment with MAPQ >= 20, > 90% identity, and covering > 50% of the read length. Additionally, reads assigned to the rDNA arrays, corresponding to the 5 known gaps in CHM13 v1.0, were ignored. Only 29,320 reads (524 Mbp) remained unaligned to the assembly. Assuming 32.4X coverage, this is only 15 Mbp of potentially missing bases (0.5% of the genome) and is likely an overestimate due to chimeric or erroneous reads. Comparatively, the v0.7 assembly has 154,951 reads (2,823 Mbp), or 87 Mbp of missing bases (2.85% of the genome). Both alignment-based and k-mer based analyses support that CHM v1.0 is highly complete.

### 6. Technology-specific sequencing biases

Despite overall uniform coverage by HiFi read mappings (with the known exception of GA-rich sequence), we observed that some multi-megabase long regions within the CHM13v1.0 assembly showed consistent excess or depletion of coverage. These corresponded to human satellite (HSat) regions (Fig. S24). While some small errors likely remain in the complex HSat arrays, the observed coverage discrepancies are very unlikely to originate from potential assembly errors, and instead point to sequencing bias. First, in all cases, coverage excess and depletion events were consistent with the satellite's class (increased coverage corresponded to Hsat2/3, and depleted coverage to Hsat1A). Second, analysis of HiFi read alignments in the regions of increased coverage revealed relatively few sites with a high-frequency of second-most common bases (Fig. S17), and also did not reveal extensive coverage spikes (beyond the uniform increase in the entire region). These observations indicated the absence of large-scale repeat collapses (*23*, *53*). Third, coverage anomalies were inconsistent between HiFi and ONT sequencing libraries with ONT read alignments showing no increase of coverage for HSat2/3 arrays, and an even larger depletion (approximately 50%) of HSat1A arrays coverage (Fig. S17). To further investigate this phenomenon we evaluated various statistics (identity, length by strand, etc) for ONT reads mapped across instances of HSat1, HSat2 and HSat3 satellite arrays and visualized them with IGV (*42*) in Fig. S18-20.

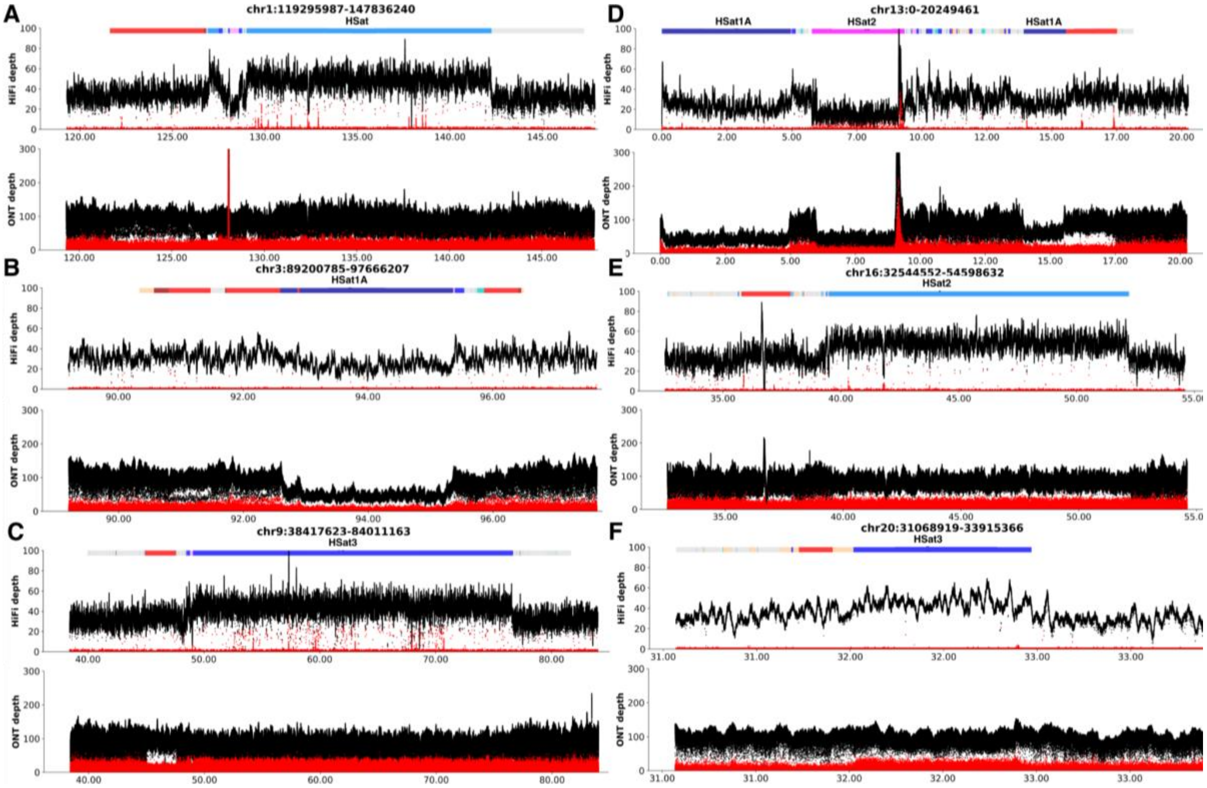


**Fig. S17: HiFi alignments for six genomic regions (with repeat annotation track showing HSat1A in light blue, HSat2/3 in dark blue, and AlphaSat in red).** Read coverage for HiFi and ONT mapped data is shown on the Y axis in black and second most frequent allele (not in the assembly) in red. Assembly position is shown on the X axis in Mbp. (A) Chr1:119295987-147836241 with the HSat2 region in dark blue (B) Chr3:89,200,785-97,666,208 with HSat1A in light blue (C) Chr9:38,417,623-84,011,164 with HSat3 in dark blue (D) Chr13:0-20,249,462 with two HSat1A regions in light blue and rDNA in pink (E)

Chr16:32,544,552-54,598,633 with HSat2 region in dark blue (F) Chr20:31068919-33915366 with the HSat3 region in dark blue. Repeat collapse events are commonly manifested by spikes of alternative base frequency, arising when reads originating in non-identical genomic repeat copies get mapped to the same assembly region (*53*). The low number of alternative-base frequency spikes across most Hsat2/3 arrays (with the only exception of the Hsat3 array on chr9 and the model-based rDNA on chr13), comparable with their density in the surrounding regions, indicates that the elevated coverage is unlikely to arise from the collapse of genomic repeat copies within the assembly. In contrast with HiFi, ONT read alignments do not display elevated coverage across the Hsat2/3 arrays on Chr1, Chr9, Chr16, and Chr20 and reveal even more depletion in HSat1A array coverage on Chr3 and Chr13. The inconsistency in coverage between sequencing technologies is indicative of technology biases. While HSat3 on Chr9 shows an elevated rate of alternative-base frequency spikes we note that assembly of this region is difficult to evaluate as it is one of the most repetitive regions in the genome (see 'Mappability' section) and many artifacts may stem from mapping ambiguities rather than assembly issues.

**A**

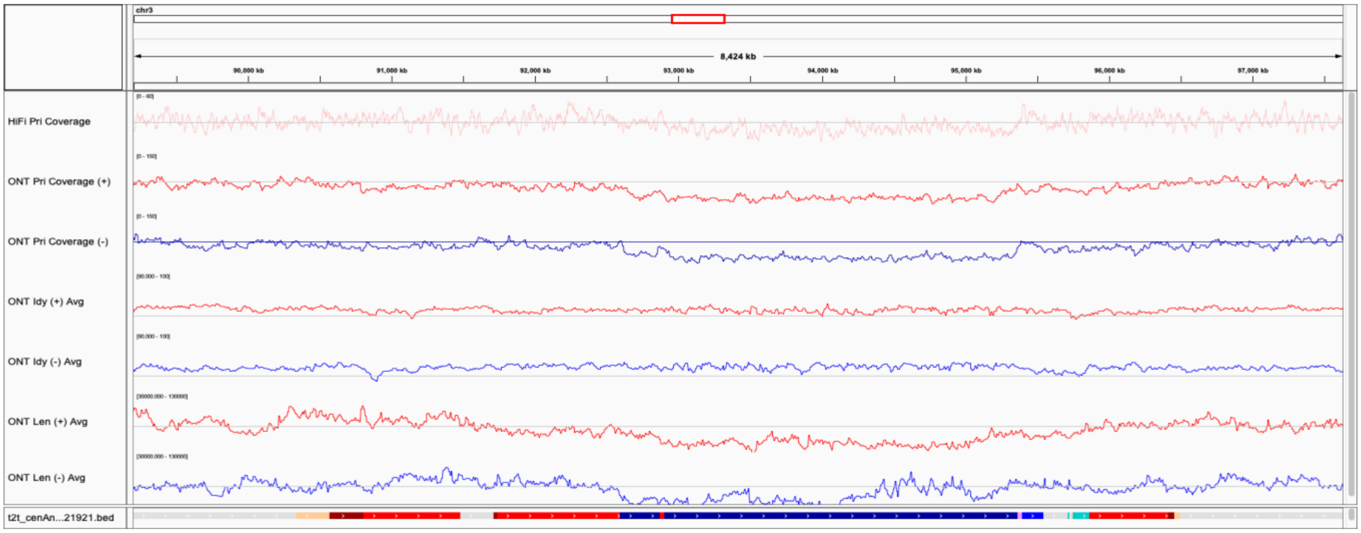


**B**

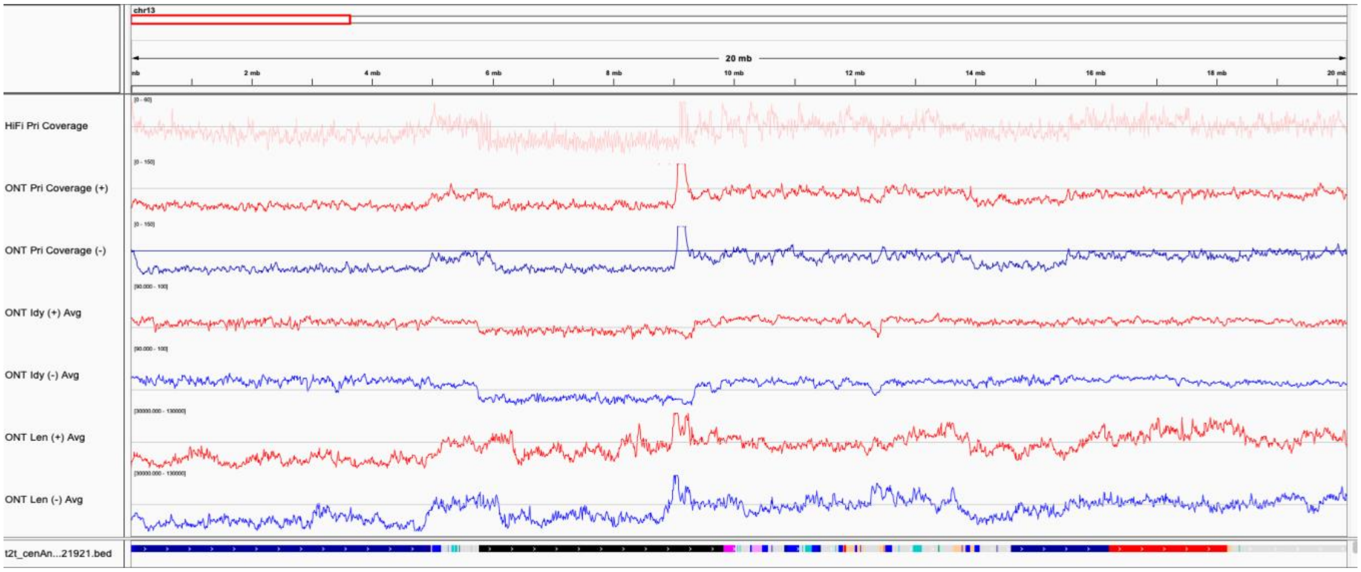


**Fig. S18: HiFi and ONT read statistics within HSat1 regions on Chr3 and Chr13.** (A) read statistics for the HSat1A region on chr3 (same region as on Fig. S17B, satellite array marked in dark blue in the annotation track). Top to bottom: coverage depth by 'primary' HiFi alignments, coverage depth by 'primary' ONT alignments by strand, identity of ONT read alignments by strand, and ONT read length by strand. Statistics were averaged in non-overlapping 10 kbp windows (see ‘Alignment-based validation’ section). The gray lines correspond to genome-wide means for each track. HiFi identity and read lengths are excluded because of their near-uniform identity and the tight library selection prior to sequencing. (B) Same statistics within two HSat1A regions on chr13 (same as in Fig. S17D, satellite array marked in dark blue in the annotation track). Coverage depletion can be seen across both ONT and HiFi reads for all shown satellite arrays. Note the decreased read length of the ONT reads mapping to the satellite arrays.

**A**

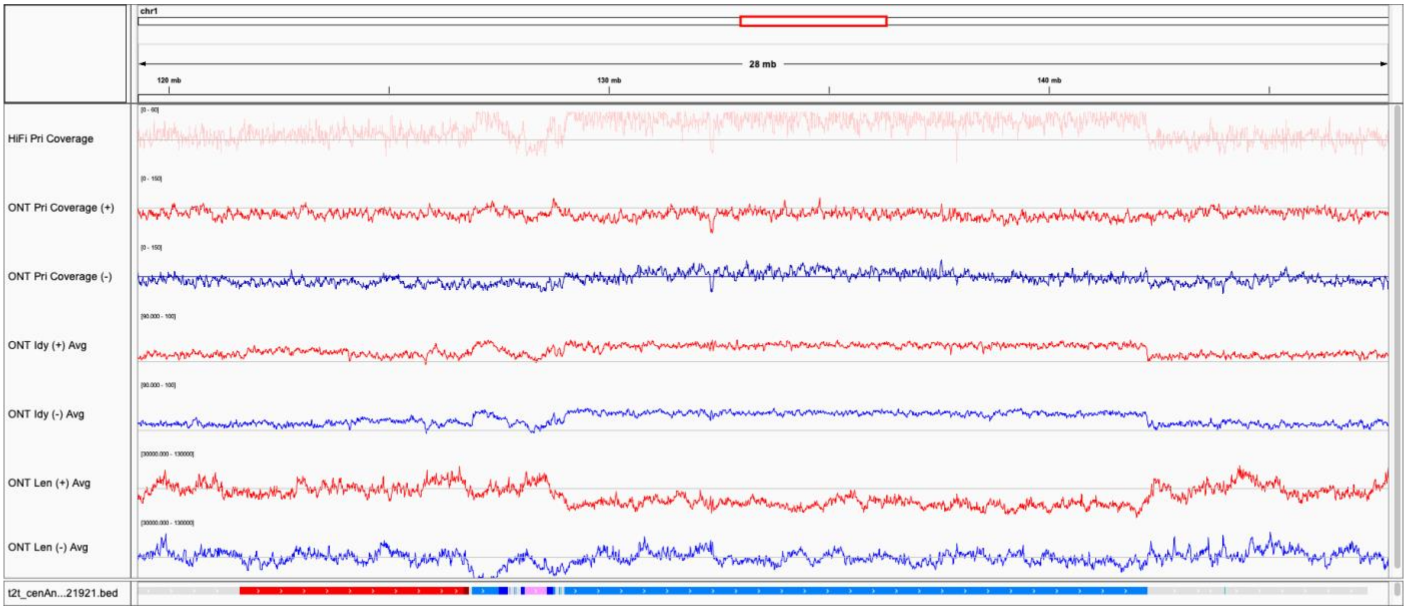


**B**

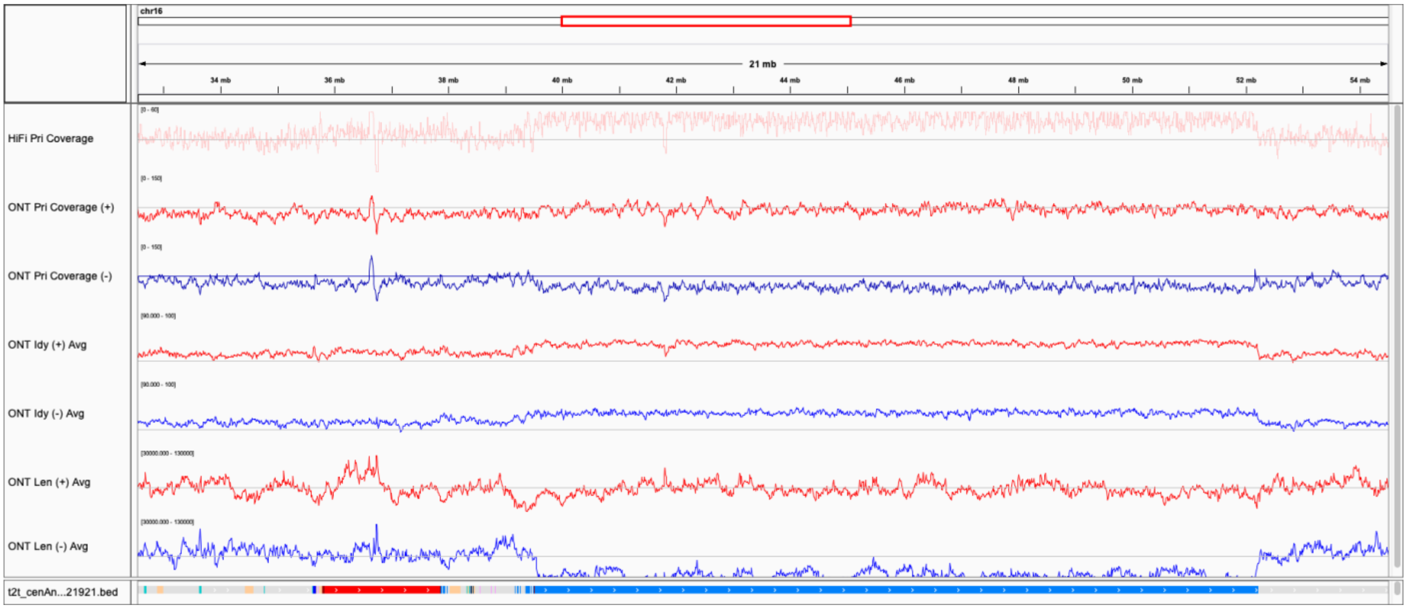


**Fig. S19: HiFi and ONT read statistics within HSat2 regions on Chr1 and Chr16.** (A) read statistics for the HSat2 region on Chr1 (same region as on Fig. S17A, satellite array marked in light blue on the annotation track). Note that the HSat2 array on Chr1 is in reverse orientation, with respect to the canonical unit, so its ‘forward’ strand corresponds to the ‘reverse’ strand of the array on Chr16. (B) Same statistics within the HSat2 region on Chr16 (same region as on Fig. S17E, satellite array marked in light blue on the annotation track). Note the drop of ONT read length on one strand (the negative strand of the repeat array) and an increase in identity on both strands.

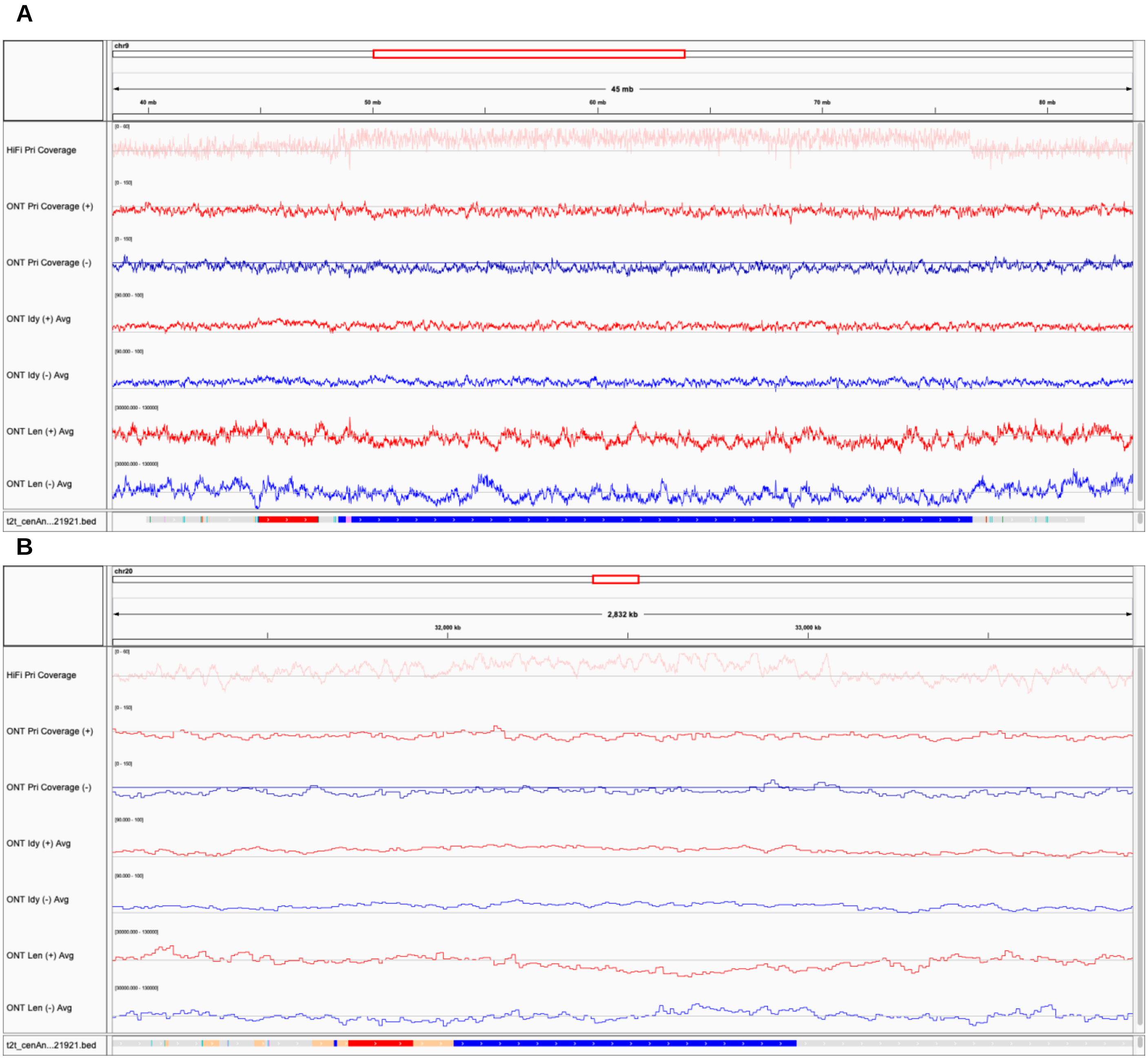


**Fig. S20: HiFi and ONT read statistics within an HSat3 region on Chr9 and Chr20.** (A) read statistics for the HSat3 region on Chr9 (same as in Fig. S17C, satellite array marked in dark blue in the annotation track). Overall, HSat3 tracks exhibit similar behavior to HSat2 in HiFi data but are closer to even coverage in ONT data and have less strand bias than HSat2. (B) Same statistics within HSat3 region on Chr20 (same as in Fig. S17F, satellite array marked in dark blue in the annotation track).

To summarize the properties of PacBio and ONT reads from different types of satellite regions, a mapping-independent strategy was implemented. In addition to the three HSat classes, alpha satellite (AlphaSat) was added as an example satellite with no HiFi coverage abnormalities. For each satellite class of interest, namely HSat1A, HSat2, HSat3, and AlphaSat relevant ONT reads (both Bonito and Guppy base-called), HiFi reads, and raw Pacbio subreads from which HiFi sequences were generated were recruited. Raw subreads were considered to assess the performance of the SMRT sequencing step separately from the subsequent subread consensus step (generating HiFi reads). Sequence assignment was performed using pre-identified sets of class and strand-specific marker *k*-mers (*k*=21). Namely, a 1 kbp long sequence was assigned to a particular satellite class if it contained 5 or more class-specific *k*-mers. A read/subread was assigned according to the majority rule across the assignments of its non-overlapping 1 kbp windows, with ties broken randomly. Relevant summary statistics, involving assigned reads and their class-assigned 1 kbp windows are presented in Table S3.

The enrichment of HSat2 in HiFi reads (>53%) is more pronounced than in the raw PacBio subreads (<45%, Table S3), a finding that is consistent with HSat2 assigned polymerase reads having the highest HiFi conversion rates across considered satellite classes (57.37% vs 44.08%, 50.52% and 52.59% for HSat1A, HSat3, and AlphaSat, respectively). Polymerase reads containing a subread assigned to a particular class counted as successfully converted if it resulted in a HiFi read assigned the same class. However, consistent values across HiFi and raw PacBio subreads in Table S3 show that the computational consensus step is unlikely to be the major factor of observed HSat enrichment/depletion in the PacBio HiFi dataset.

In ONT data, HSat1A and HSat2 were associated with shorter read lengths than AlphaSat and surrounding regions (Table S3, Fig. S18-19). We hypothesize that the AT-rich HSat1A may be unstable during library preparation and shears preferentially, with shorter molecules leading to lower coverage by both ONT data (due to shorter read lengths) and HiFi data (due to fewer molecules passing strict size selection). ONT signal processing methods have known biases in AT-rich repeats (*54*), which could lead to fewer reads successfully passing quality filters and even incorrect detection of molecule boundaries. ONT data associated with HSat2 exhibits a significant strand bias (Table S3 and Fig. S19), with a high depletion only in sequencing reads coming from the ‘reverse’ strand (“GAATG”-unit) vs the ‘forward’ strand of the repeat array (“CATTC”-unit). Note that the entire HSat2 array on Chr1 was inverted with respect to the canonical unit orientation. However, the higher accuracy of ONT Guppy reads and regular coverage of the forward strand masks these effects in ‘unstranded’ mapping-based coverage plots (Fig. S17). ONT reads were also depleted in AlphaSat while HiFi is consistent with expectation.

The biases exhibited by ONT read sets differed across Guppy and Bonito base-callers. While Bonito base-calling positively affected (reduced) AlphaSat and HSat2 depletion and reduced the strand bias, there was a negative impact on the identity of the reads from HSat1A.

**Table S3: Summary statistics for reads recruited across different satellite classes.**

| Data type | HSat | Total length (Mbp) | Expected length (Mbp) | Enrichment | Avg. read length (kbp) |
|---|---|---|---|---|---|
| ONT Guppy 3.6.0 | HSat1A+ | 0.37 | 0.83 | -55.18% | 29.43 |
| | Hsat1A- | 0.40 | 0.83 | -51.43% | 29.05 |
| | HSat2+ | 1.73 | 1.79 | -3.09% | 29.40 |
| | Hsat2- | 1.19 | 1.79 | -33.12% | 21.68 |
| | HSat3+ | 2.54 | 2.97 | -14.38% | 37.52 |
| | HSat3- | 2.19 | 2.97 | -26.12% | 36.51 |
| | AlphaSat+ | 4.27 | 5.31 | -19.65% | 34.27 |
| | AlphaSat- | 4.26 | 5.31 | -19.67% | 33.90 |
| ONT Bonito 0.3.1 | HSat1A+ | 0.36 | 0.80 | -54.27% | 32.06 |
| | Hsat1A- | 0.38 | 0.80 | -51.65% | 31.10 |
| | HSat2+ | 1.79 | 1.71 | 5.04% | 30.62 |
| | HSat2- | 1.25 | 1.71 | -26.77% | 21.57 |
| | HSat3+ | 2.22 | 2.83 | -21.79% | 39.27 |
| | HSat3- | 2.26 | 2.83 | -20.30% | 40.33 |
| | AlphaSat+ | 4.57 | 5.07 | -9.83% | 36.35 |
| | AlphaSat- | 4.67 | 5.07 | -8.05% | 36.40 |
| Pacbio subreads | HSat1A+ | 1.36 | 1.92 | -29.20% | 17.17 |
| | HSat1A- | 1.43 | 1.92 | -25.67% | 17.82 |
| | HSat2+ | 5.96 | 4.12 | 44.73% | 17.09 |
| | HSat2- | 5.92 | 4.12 | 43.88% | 17.10 |
| | HSat3+ | 9.54 | 6.83 | 39.64% | 17.31 |
| | HSat3- | 9.12 | 6.83 | 33.45% | 17.04 |
| | AlphaSat+ | 12.44 | 12.23 | 1.68% | 16.93 |
| | AlphaSat- | 12.61 | 12.23 | 3.07% | 17.08 |
| PacBio HiFi | HSat1A | 0.23 | 0.32 | -27.55% | 19.14 |
| | HSat2 | 1.05 | 0.68 | 53.13% | 19.14 |
| | Hsat3 | 1.58 | 1.14 | 39.13% | 19.11 |
| | AlphaSat | 2.16 | 2.03 | 6.32% | 18.99 |

For all types of data, except for HiFi reads, the statistics are additionally broken out by strand (orientation with respect to the canonical unit). The ‘Total length’ is the total size of all 1 kbp windows assigned to a particular satellite class. ‘Expected length’ was computed by multiplying the total length of reads within a particular dataset by the fraction of the assembly assigned to a particular satellite class (more precisely, fraction of the assembly bases within the 1 kbp windows that were assigned to a particular satellite class based on *k*-mer analysis). Assigned assembly fraction values were 0.44% for HSat1, 0.94% for HSat2, 1.56% for HSat3, and 2.79% for AlphaSat. Enrichment was computed based on the ‘total’ vs ‘expected’ length values, as (‘total’ - ‘expected’ / ‘expected’). ‘Avg. read length’ provides average length across assigned reads/subreads.

Finally, CLR kinetic information was inspected for PacBio polymerase reads originating from different types of satellite regions (Fig. S21). Kinetic information was collected for all ZMWs with at least 3 passes (but may not have produced a HiFi read) and had at least one subread assigned to the particular satellite class based on *k*-mer analysis. Unfortunately, this analysis did not lead to conclusive insights into the origin of HSat enrichment phenomena.

HSat2,3 arrays were associated with slightly longer sequencing times and polymerase lengths in PacBio data, compared to AlphaSat and whole-genome stats (Fig. S21). We hypothesize that these regions might be 'easier' for the PacBio polymerase to sequence, which would lead to more reads surviving the initial pre-extension step, and in turn contribute to the observed enrichment. HSat1A arrays are depleted in the longest (>250 kbp) polymerase reads, and are associated with slightly shorter sequencing time (vs AlphaSat and whole-genome).

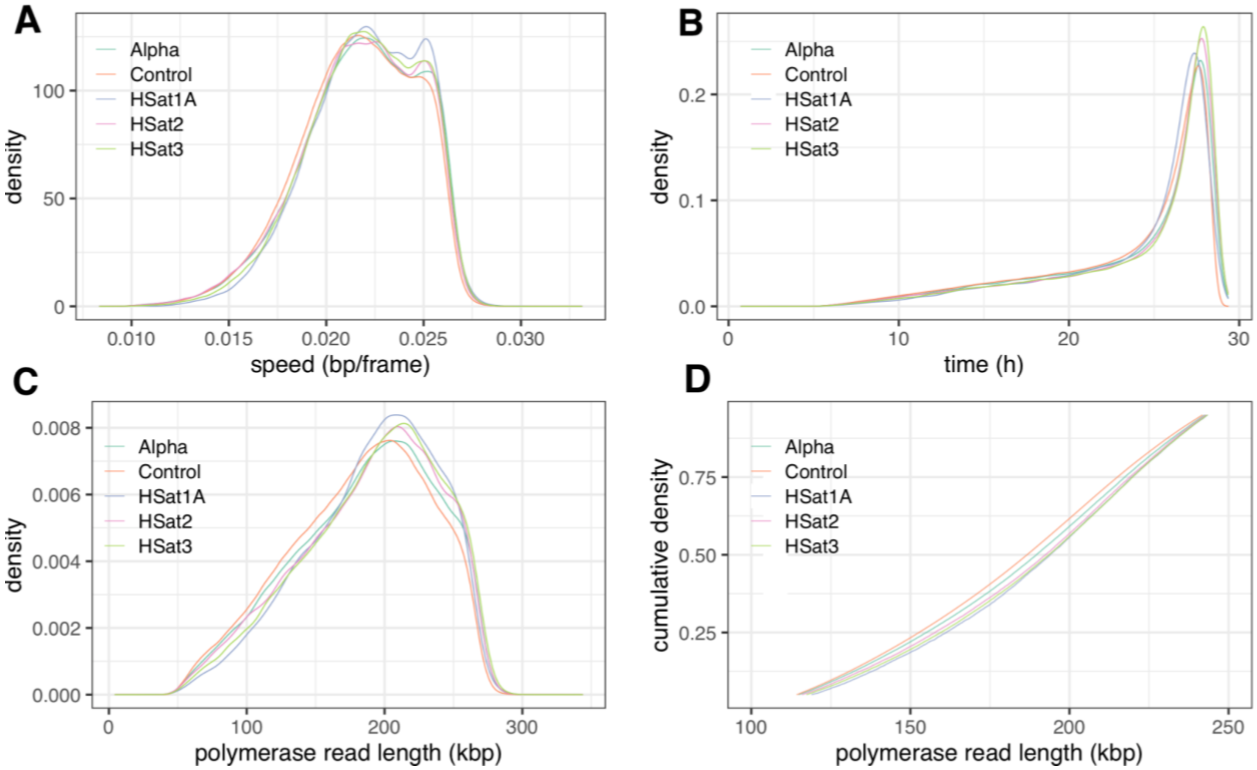


**Fig. S21: Kinetic information for reads from HSat and AlphaSat regions as well as all reads (Control).** The plots from top to bottom show (A) speed of sequencing, (B) total time sequencing a molecule, and polymerase read length ((C) regular and (D) cumulative). In all plots AlphaSat behaves similarly to the 'Control'. (A) No drastic difference in sequencing speeds was observed between the considered classes (AlphaSat in teal, HSat1A in blue, HSat2 in pink, and HSat3 in green), with all showing a bi-modal distribution. However, satellite reads are elevated in the second (faster) peak relative to control, with HSat1A having the largest increase. (B) The reading time for a polymerase read is longer for HSat2,3 and shorter for HSat1A, consistent with observations about polymerase read lengths. (C/D) The polymerase reads are of similar length, with a slight shift to longer reads in HSat2,3. HSat1A reads have higher density of reads in the 150-250 kbp range but then drop off and have lower fraction of polymerase reads longer than 250 kbp than AlphaSat or HSat2,3. Satellite reads are shifted to longer lengths when compared to the control.

Here, many complex human satellites have been assembled for the first time (*44*). The potential causes leading to the observed coverage biases were briefly explored, but more analysis is needed in future. It is unsurprising that machine-learning methods relying on learning from known sequence contexts, e.g. ONT base-calling and HiFi subread consensus, performed poorly in these regions. We hope that the availability of a finished human reference along with our observations will lead to improvements in the quality of both ONT and PacBio reads originating from challenging satellite genomic regions in the near future.

### 7. Alpha-satellite and human satellite 1,2,3 validation

TandemTools (*30*) provided orthogonal validation for the assembly of alpha and human satellite arrays, excluding rDNA (Table S4; Fig. S22). Quality assessment with both accurate PacBio HiFi and error-prone ONT reads revealed positions in the assembly with consistent discrepancies between mapped reads and the assembly in distances between consecutive rare *k*-mers (default $k = 19$), a *k*-mer was considered rare if its frequency in the assembly did not exceed 10 (*30*). A list of regions where the discrepancy was supported by at least 30% of covering reads and may indicate heterozygous sites (*34*) was compiled. All putative events within the centromeric satellite DNA were further manually inspected in IGV for validity and catalogued at https://github.com/marbl/CHM13-issues. Two sites with 100% deviated reads (on Chromosomes 18 and 19) corresponded to a low coverage region (Chr18) and a potential 2.4 kbp insertion in the assembly (Chr19). The insertion in Chromosome 19 was not corrected as the alternate sequence had ambiguous mapping support. Besides that possible error, arrays of centromeric satellite DNA do not contain large structural mis-assemblies (longer than 100 bp).

**Table S4. Coordinates of alpha and human satellite arrays in v1.0 assembly.**

| Chr. | Start | End |
|---|---|---|
| chr1 | 116796216 | 147241828 |
| chr2 | 85991672 | 99673016 |
| chr3 | 85805192 | 101415517 |
| chr4 | 44705247 | 59870604 |
| chr5 | 42077197 | 54596619 |
| chr6 | 53286920 | 66058622 |
| chr7 | 55414368 | 68714496 |
| chr8 | 39243541 | 51325076 |
| chr9 | 39952789 | 81694033 |
| chr10 | 34633784 | 46664580 |
| chr11 | 46061948 | 59413485 |
| chr12 | 29620490 | 42202482 |
| chr13 | 0 | 23171058 |
| chr14 | 0 | 17765925 |
| chr15 | 0 | 23279251 |
| chr16 | 30848291 | 57219476 |
| chr17 | 18892710 | 32487230 |
| chr18 | 10965698 | 25933550 |
| chr19 | 19655572 | 34768168 |
| chr20 | 21383653 | 37969531 |
| chr21 | 0 | 17078862 |
| chr22 | 0 | 20739833 |
| chrX | 52820107 | 65927026 |

Coordinates of alpha and human satellite arrays in v1.0 assembly of CHM13 that were validated by TandemTools (*30*). The rDNA arrays on chromosomes 13, 14, 15, 21, and 22 were excluded from this validation. Corresponding TandemTools plots are shown in Figure S22.

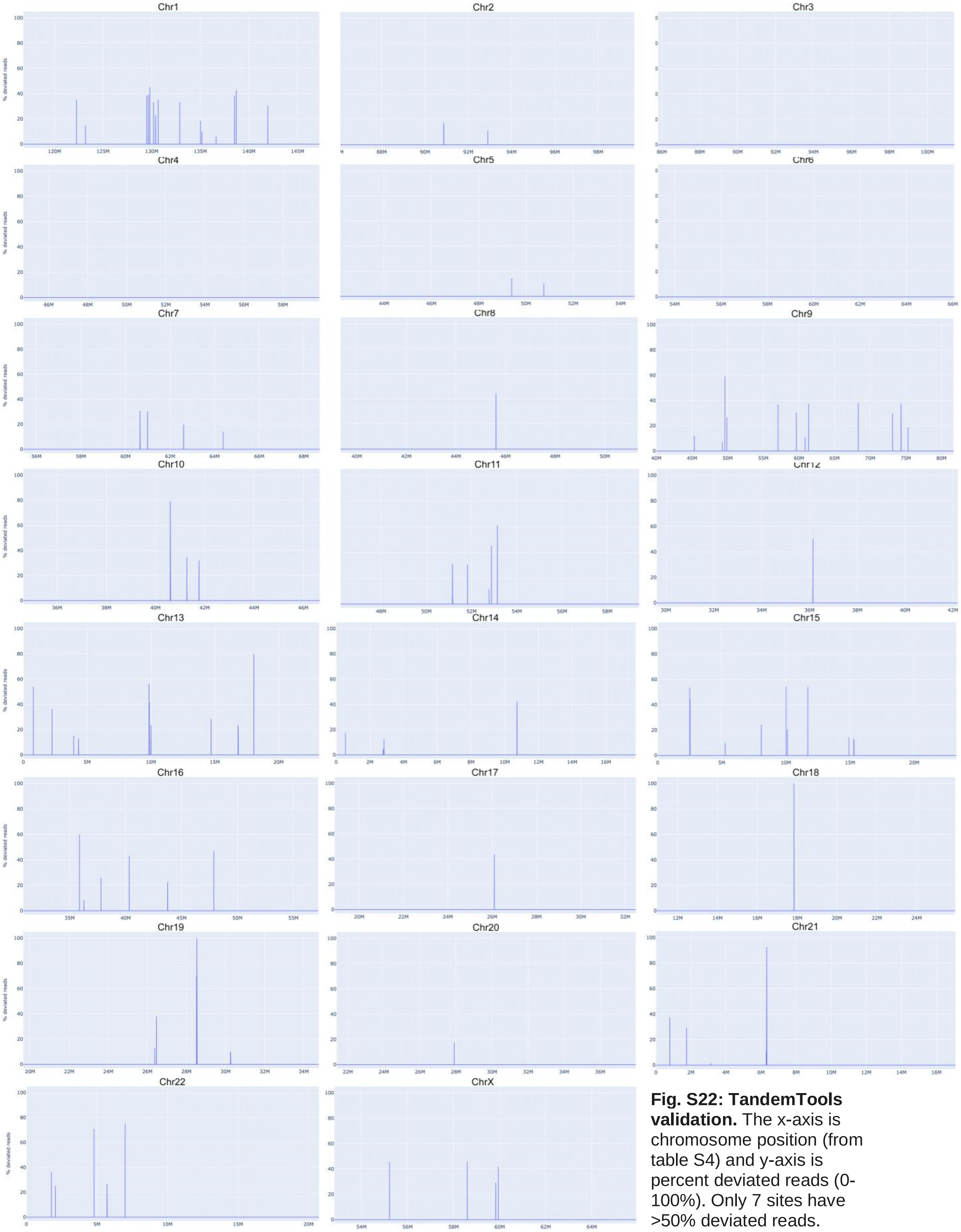


**Fig. S22: TandemTools validation.** The x-axis is chromosome position (from table S4) and y-axis is percent deviated reads (0-100%). Only 7 sites have >50% deviated reads.

The proportion of monomers of each live higher-order repeat (HOR) in sequencing reads versus the same proportion calculated from the HOR annotation (*44*) in our v1.0 assembly were also compared. The expected length of AS monomers was ≈ 170 bp. The number of the live HORs was 19, including S2C13/21 and S2C14/22 each located on 2 chromosomes and S1C1/5/19 located on 3 chromosomes. In these cases, the HOR counts were divided by 2 for doubles and by 3 for the triple. The sister HORs of the live HORs were not included (e.g. S3C17H1-B and C). Only the near full-length monomers (length >= 160 bp) were counted to reduce the risk of erroneous monomer classification. Monomer classification/identification was performed by HumAS-HMMER-HOR (https://github.com/enigene/HumAS-HMMER) described in (*55*) (for SF1 HORs only) and in (*44*) (all known HORs covered). Here, a HMMER platform (*56*) was used to identify (as the best match in a set of standards) each monomer of about 80 known HORs and about 30 classes of monomers in AS monomeric layers. Note that for monomers of homogeneous HORs only one representative was selected so the best match was equivalent to the best alignment. For divergent HORs, the standard was formed using a multi-alignment of representative monomers and therefore constitutes a true HMM. Conversion of the HMMER output file into a bed file was performed using the hmmertblout2bed script (https://github.com/enigene/hmmertblout2bed) (*55*).

**A** **B**

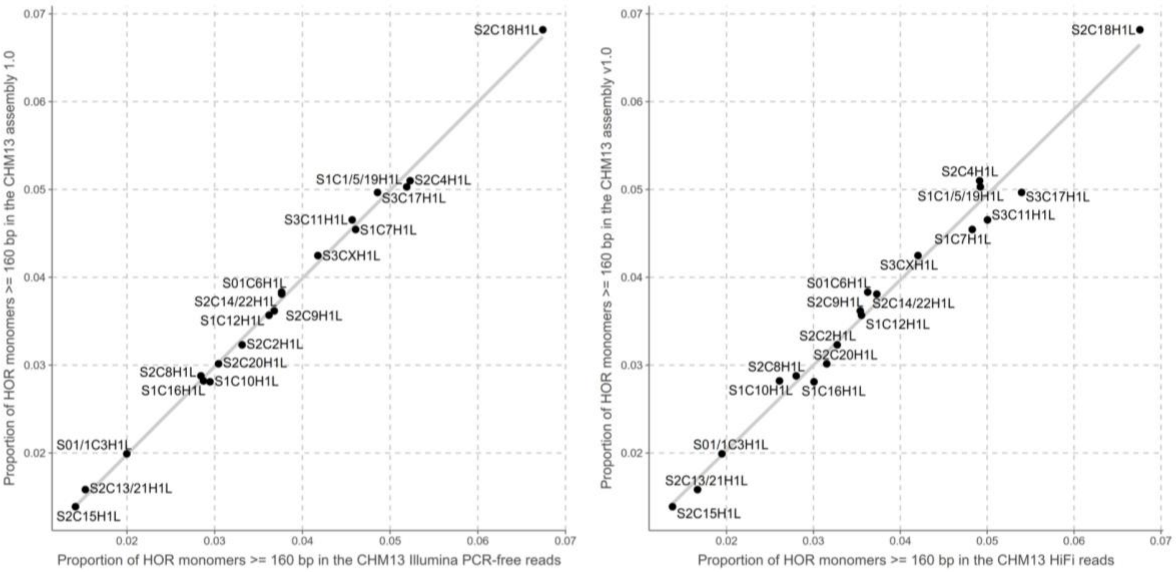


**Fig. S23: Proportions of HOR monomers in the CHM13v1.0 assembly vs PCR-free Illumina data (A) and HiFi data (B).** The line shown in the panels was calculated using a linear regression model in R (lm function) and geom_smooth function from the ggplot package. Note that the wider distribution of dots around the line in the HiFi sample may be due to more frequent sequencing errors in the HiFi reads, as compared to the Illumina reads.

Monomers were counted in both the PCR-free Illumina 250 bp data and the HiFi 20 kbp libraries. In the short read sample, a total of 8,314,086 HOR monomers were identified and 2,045,562 HOR monomers remained post length filtering. In the long read sample a total of 3,792,752 HOR monomers were identified and 3,720,342 HOR monomers remained post length filtering. In the assembly, a total of 426,773 HOR monomers were identified and 425,732 HOR monomers remained post length filtering (Fig. S23). There was a good agreement between the proportion of monomers in the sequencing data and the assembly for both data types.

HiFi data was aligned and the frequency of secondary alleles plotted as in (*23*). Reads were aligned with the following command: `pbmm2 align --log-level DEBUG --preset SUBREAD --min-length 5000` and filtered with `samtools -F 2308`. Plots were generated using NucFreq's `NucPlot.py` script (https://github.com/mrvollger/NucFreq) for selected regions (Fig. S24). Assembly loci with high second most frequent allele counts arise due to mis-assembly or heterozygous sites. Since CHM13 is homozygous (with few exceptions), we did not expect heterozygous sites. As expected, the assembly had a low level of secondary alleles and had even coverage, with the exception of sequencing artifacts in HSat regions (see 'Technology-specific biases' section). The rDNA arrays had depleted coverage and spikes at the borders due to being model sequences. Note that simpler rDNA arrays (see 'Resolution of rDNA arrays' section) on chromosomes 14, 22 showed even coverage through the array while model-based resolutions (chromosomes 13, 15, 21) showed a decrease in coverage and an increase in secondary allele frequency.

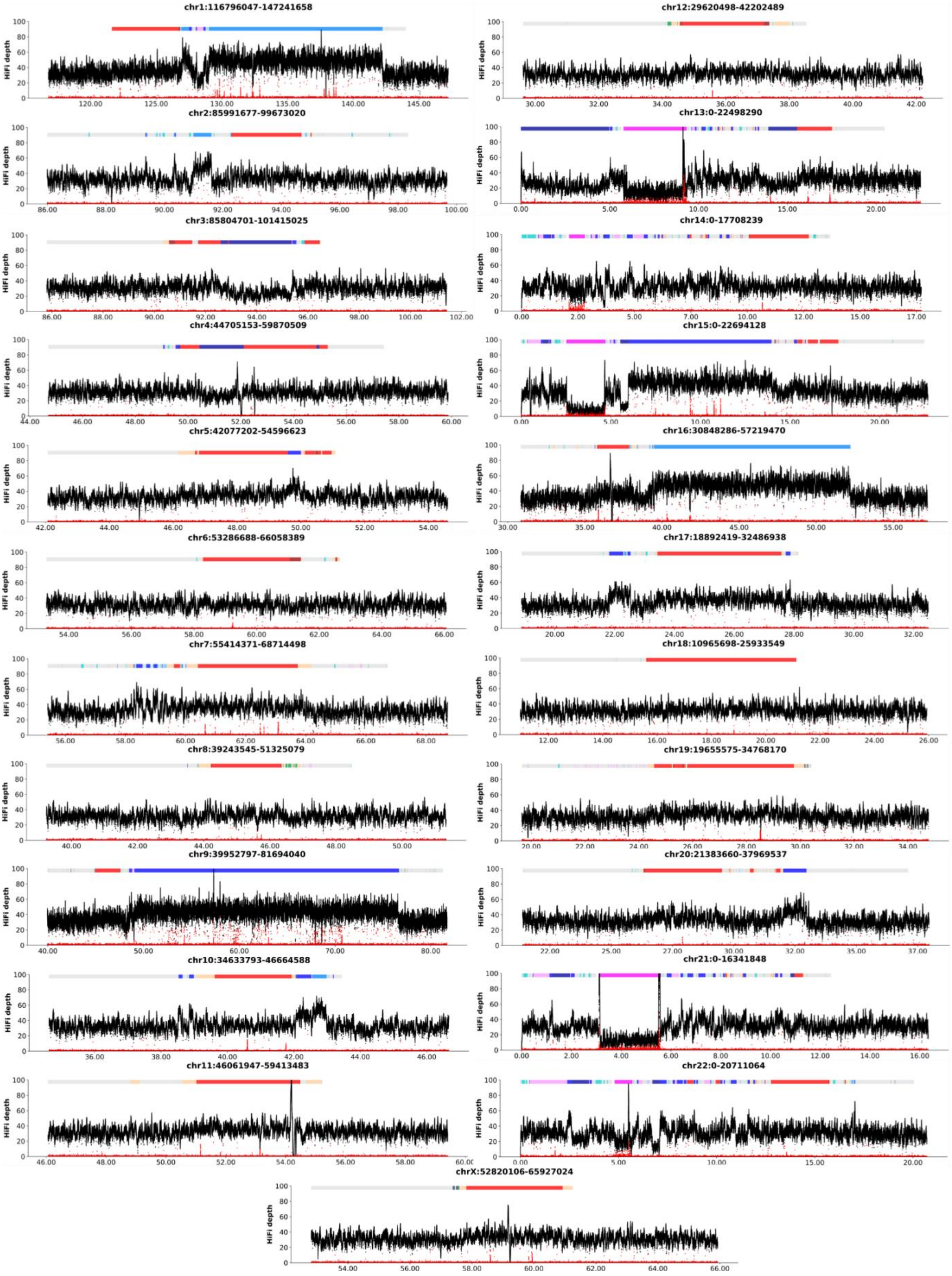
chr1:116796047-147241658
chr12:29620498-42202489
chr2:85991677-99673020
chr13:0-22498290
chr3:85804701-101415025
chr14:0-17708239
chr4:44705153-59870509
chr15:0-22694128
chr5:42077202-54596623
chr16:30848286-57219470
chr6:53286688-66058389
chr17:18892419-32486938
chr7:55414371-68714498
chr18:10965698-25933549
chr8:39243545-51325079
chr19:19655575-34768170
chr9:39952797-81694040
chr20:21383660-37969537
chr10:34633793-46664588
chr21:0-16341848
chr11:46061947-59413483
chr22:0-20711064
chrX:52820106-65927024
HiFi depth

**Fig. S24: NucFreq plot of centromeric and satellite regions in CHM13 v1.1.** HiFi coverage depth (black) along with secondary allele frequency (red) for all centromere and surrounding regions. Top bar shows annotations (*44*) with red corresponding centromeric HORs, light blue HSat1A, dark blue HSat2/3, and hot pink rDNA.

### 8. Resolution of the rDNA arrays

The five rDNA arrays on the acrocentric chromosomes 13, 14, 15, 21 and 22 are highly repetitive regions consisting of long tandem repeats with a repeat motif size of approximately 45 kbp, which further consists of other tandem repeat units. It was observed that most rDNA arrays consisted of a repeating major morph with a few distinct repeat units close to the PJ and DJ regions. Based on this observation, model array sequences were created by filling in the gaps between confidently assembled boundary repeat units of the arrays with copies of the major morph. The number of copies was estimated using both molecular and bioinformatic techniques, which resulted in consistent estimates (Figs. S1,S3,S25).

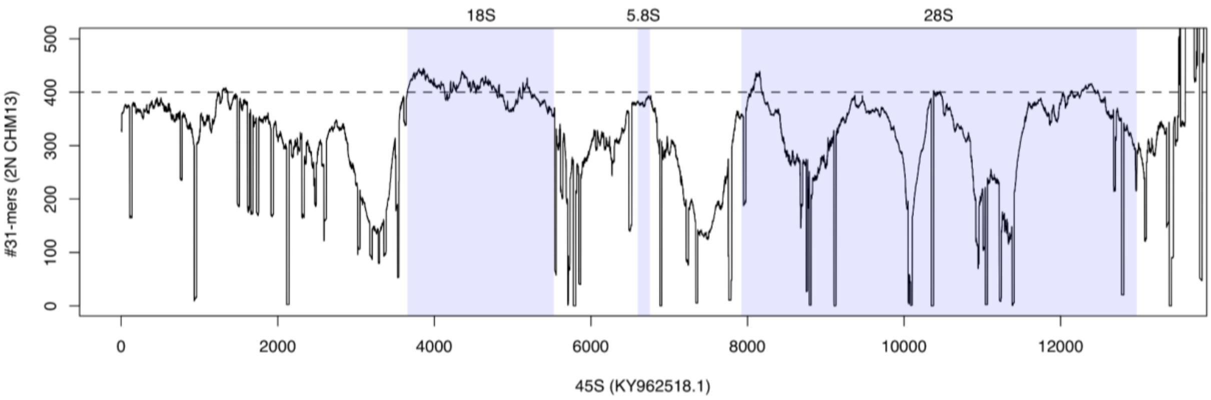


**Fig. S25: 45S *k*-mer count distribution in CHM13.** Diploid copy number was estimated from ILMN *k*-mers. The x-axis shows the canonical rDNA unit, and the y-axis the estimated (diploid) count of those *k*-mers in the CHM13 genome. Total ILMN sequencing depth was estimated to be 98X based on the modal *k*-mer copy number (Fig. S15), or 49X per haplotype, so the diploid copy number was estimated as the raw *k*-mer count divided by 49. Sharp drops in copy number correspond to small variants, while the smoother dips are due to larger indels and %GC bias in the ILMN data. The 18S gene shows relatively uniform coverage due to its high degree of conservation in the genome and more typical %GC (56%). This plot confirms a diploid copy number of ~400 rDNA units, or ~200 per haplotype, in the CHM13 genome, consistent with the FISH and ddPCR estimates.

A custom pipeline was used to analyze the rDNA arrays and recover consensus sequences of distinct repeat ‘morphs’. HiFi reads were first recruited based on a previous rDNA consensus sequence (*2*). The reads were then separated by chromosome via building a sparse de Bruijn graph constructed from recruited reads with MBG (*29*) (k=3501), in which five rDNA arrays corresponded to separate connected components (Fig. S26A). An array-specific sparse DBGs with k=201 from the HiFi reads (Fig. S26B) was then built and ONT reads were assigned to a particular array and corrected by aligning them to the chromosome-specific HiFi graphs with GraphAligner. Aligned read sequences were replaced by the alignment path in the graph. All read fragments were extracted, representing array-specific rDNA repeats from the corrected ONT reads, and grouped based on the pairwise edit distances through single-linkage clustering with a distance threshold of 200 (corresponding to a 99.5% identity). A consensus sequence for individual clusters, which was considered to represent distinct repeat morphs, was built with SPOA (*57*). Note, this approach was not sensitive enough to discern small SNP-level variations below the clustering threshold. For each array, the major morphs were identified, i.e. the morph corresponding to the highest number of ONT read fragments, and polished with Racon (*57*) using array-specific HiFi reads. Information of ONT reads spanning between different morphs was used to build a graph of morphs to represent the structure of the rDNA arrays (Fig. S26C) showing the simple structure of chromosomes 14 and 22 and the complex mosaic structure of chromosomes 13, 15 and 21.

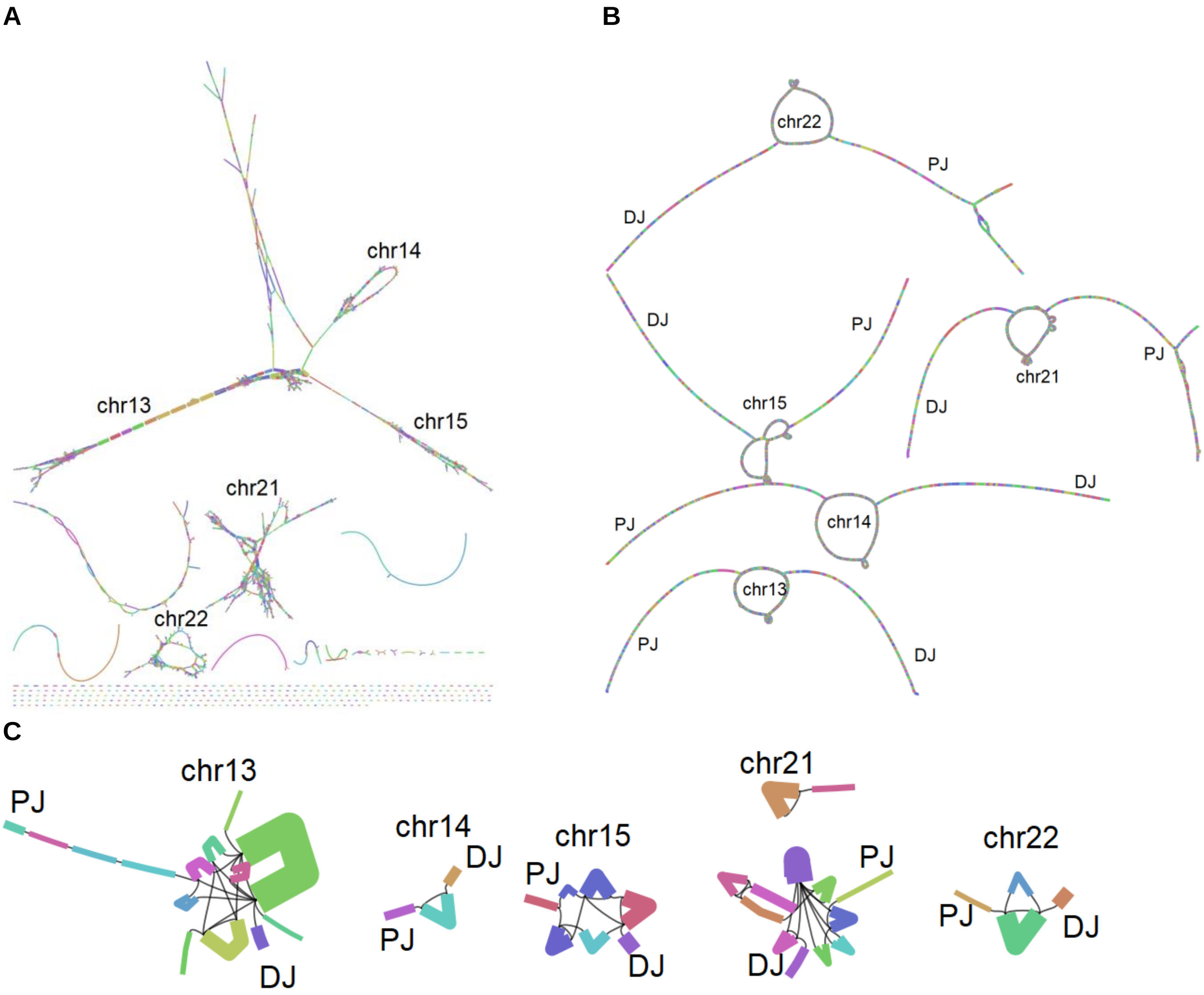


**Fig. S26: Overview of rDNA assembly process.** (A) A minimizer graph of rDNA HiFi reads with k=3501. Due to chromosome specific variation, the graph has five large clusters, one per chromosome. The chromosome specific variation enables clustering the HiFi reads by chromosome based on shared k-mers between the reads and the sequences of the nodes in each cluster. The graph is fragmented due to systematic coverage bias and sequencing errors in the HiFi reads, especially within the GA-rich regions of the intergenic spacer. (B) Minimizer graphs built from chromosome specific HiFi reads with k=201. The loop structure represents the rDNA array, including both morph-specific variation and sequencing errors. (C) A graph built from clustered ONT reads. Each node represents one cluster, and the thickness of the nodes corresponds to ONT read coverage. Edges are added whenever at least two ONT reads support a connection between the clusters. The clusters recover structurally unique rDNA morphs, although morphs with only minor SNP variation may be collapsed to the same cluster. The graph shows that chromosomes

14 and 22 consist of one major morph repeating multiple times while chromosomes 13, 15, and 21 have a more complex mosaic structure.

To accurately reconstruct the boundary repeat units of the rDNA arrays, ONT reads were extracted that anchored to the unique sequence in the PJ/DJ regions surrounding the rDNA arrays and separated by chromosome, PJ/DJ arm, and haplotype (heterozygous structural variants have been identified within some arrays (Fig. S1)). Racon was used to build a consensus for each group of ONT reads. To further correct the ONT-based consensus sequences using HiFi reads they were aligned to an updated HiFi string graph and substituted by the alignment path sequence. This approach allowed us to accurately recover several copies of the rDNA repeat unit at the ends of every array.

The rDNA array on chromosome 15 had a more complex structure (Fig. S26C). First, its length differed between the two haplotypes, with the shorter array comprising only 7 repeat units spanned by a single 450 kbp ONT read (Fig. S2). Second, its longer haplotype appeared to consist of multiple copies of three distinct morphs interleaved with each other. Thus, we included the three major morphs as three distinct blocks each containing the estimated number of copies. However, this does not accurately reflect the true interleaved ordering of the morphs and should be considered only a model until longer sequencing reads enable a more accurate reconstruction of this array in the future.

### 9. Genome analysis

#### Assembly tracks

A UCSC Genome Browser (*58*) hub has been released for the CHM13 v1.0 assembly available at: [http://genome.ucsc.edu/cgi-bin/hgTracks?genome=t2t-chm13-v1.0&hubUrl=http://t2t.gi.ucsc.edu/chm13/hub/hub.txt](http://genome.ucsc.edu/cgi-bin/hgTracks?genome=t2t-chm13-v1.0&hubUrl=http://t2t.gi.ucsc.edu/chm13/hub/hub.txt). These tracks were used to generate Figures 1, 3, 4, and 5 of the main text. Specifically, segmental duplication and censat annotations were computed as in (*44*, *59*) and plotted as non-overlapping regions. SD density in 10 kbp windows was computed using the kpPlotDensity (*60*) function and colored by log of the window density. Novel sequence in CHM13 was based on whole-genome LastZ (*61*) alignment of GRCh38 and CHM13, filtering for any regions without 1 Mbp of synteny (*59*). A more conservative set of regions not covered by GRCh38 was computed using Winnowmap v2.0.1 to align GRCh38 (excluding ChrY, patches, and alts) to CHM13 v1.1 with the command `winnowmap -ax asm20 -H --MD chm13.v1.1.fasta GCA_000001405.28_GRCh38.p13_genomic_noalt_noY.fasta > out.sam`. Primary alignments were converted to paf and filtered for primary only via `k8 paftools.js sam2paf -p out.sam > out.paf` and converted to unmapped regions by inverting the list of mappings w/MAPQ>0: `cat out.paf |awk '{if ($12 > 0) print $6"\t"$8"\t"$9}'|bedtools sort -i - |bedtools merge -i - |bedtools complement -i - -g chm13.sizes`. GRCh38 (*21*) issue XML files were downloaded from the GRC (ftp://ftp.ncbi.nlm.nih.gov/pub/grc/human/GRC/Issue_Mapping/) and filtered to exclude issues resolved in GRCh38 or a patch or marked as ‘Variant’ or ‘GRCHousekeeping’, resulting in 191 issues. Regions were padded by 1 Mbp, lifted over with the command `liftOver -bedPlus=3 -minMatch=0.5 hg38_issues.bed hg38.t2t-chm13-v1.0.all.chain.gz chm13_lifted_issues.bed unmapped.bed`, and trimmed back by 1 Mbp. A total of 43 issues could not be lifted over. Novel genes were identified from the combined CAT and LiftOff annotations (see ‘Annotation’ below).

The computed ancestry regions (see “Ancestry Analysis”), in GRCh38 coordinates, were lifted over from GRCh38 to CHM13 with the command `liftOver -bedPlus=3 -minMatch=0.75 ancestry.bed hg38.t2t-chm13-v1.0.all.chain.gz chm13_v1.0_lifted_ancestry.bed unmapped.bed.CHM13`.

#### Annotation

The Cactus (*62*) alignment between the v1.0 assembly and the primary contigs of GRCh38 (*21*), with chimp as an outgroup, was created with the following command:

```
cactus aws:us-west-2:t2t-jobstore-chm13 cactus-config-chm13-
t2t.draft_v1.v2.txt t2tChm13.draft_v1.v2.hal --maxCores 80 --
binariesMode local
```

Using the following config file for cactus:

```
(Chimp:0.00655,(GRCh38:0.0005,CHM13:0.0005));
Chimp   GCF_002880755.1_Clint_PTRv2_fixed.fa
GRCh38  GRCh38.primary.fa
CHM13   t2t-chm13-v1.0.fa
```

Iso-Seq reads were aligned using minimap2 (*32*) using the following command:

```
minimap2 -ax splice -f 1000 --sam-hit-only --secondary=no --eqx -
K 100M -t 4 --cap-sw-mem=3g mmdb/0.mmi iso_fastas/0_0.fasta
```

Stringtie2 (*63*) assembled the transcriptome using available Iso-Seq reads:

```
stringtie -p 8 chm13_1.t2t.sorted.bam.filtered.bam -L >
chm13_1.t2t.TM.stringtie.gtf
```

CAT (*64*) v2.2.1 (commit `96e7550f22387a669f0b98dfc0c94be825192e24`) was run in TransMap (*65*) mode using the following command:

```
luigi --module cat RunCat --hal=t2tChm13.draft_v1.v2.hal --
target-genomes='("CHM13",)' --ref-genome=GRCh38 --workers=10 --
config=cat.t2t.draft_v1.full.isoseq.config --work-dir work-chm13-
t2t --out-dir out-chm13-t2t --local-scheduler --assembly-hub --
maxCores 5 --binary-mode local
```

Using GENCODEv35 (*66*) and following config file for CAT:

```
[ANNOTATION]
GRCh38 = gencode.v35.annotation.gff3.noPAR
CHM13 = CHM13.TM.stringtie.merged.gff3
[ISO_SEQ_BAM]
CHM13 = data/chm13_1.t2t.sorted.bam,data/chm13_2.t2t.sorted.bam
```

We removed genes from the CAT annotation that had overlapping annotations from multiple genes in the same family, leaving the gene that was correct based on synteny.

The Liftoff (*67*) annotation was created with the following command using version 1.6.0:

```
liftoff chm13.draft_v1.0.fasta GRCh38.fa -sc 0.95 -copies -g
gencode.v35.annotation.gff3 -polish -chroms chroms.txt
```

To create the final annotation, we complemented the CAT result with missed Gencode genes and putative additional paralogs (with minimum sequence identity of 95%) from the Liftoff annotation. Only predictions that did not overlap any CAT annotations were added.

The annotation set on the v1.0 assembly was lifted over to the v1.1 assembly using liftover with the command

```
liftOver -gff CHM13.combined.v4.gff3 v1_to_v1.0423.chain
CHM13.combined.v4.liftover.v1.1.gff3 unmapped.txt
```

The rDNAs were annotated by mapping an assembly of an rDNA unit isolated from chromosome 21 (*2*) onto v1.1 with Liftoff. Using GenBank entry KY962518.1, the rDNA sequence was obtained and a gff3 file created with the coordinates of the 45S, 18S, 5.8S, and 28S subunits. Liftoff was then run with the following command to annotate all rDNAs within the assembly

```
liftoff chm13.draft_v1.1.fasta KY962518.1.fasta -g
KY962518.1.gff3 -copies -sc 0.95 -mm2_options=”-N 300”
```

All annotations that have been lifted over and that overlapped the newly added rDNA regions were removed. The rDNA annotations (876 new genes) were added to create a final annotation set.

In the final annotation set, 63,494 genes were annotated (19,969 protein-coding, 27,026 non-coding RNA, 15,799 pseudogenes, 652 Immunoglobulin/T-cell receptor segments, and 48 novel genes without a Biotype from StringTie2). In terms of transcripts, 233,615 transcripts were annotated (86,245 protein-coding, 57,196 non-coding RNA, 15,997 pseudogenes, 674 Immunoglobulin/ T-cell receptor segments, 56 from StringTie2, and 73,447 other types).

Further breakdown by biotype is provided by Table S5: “Gene Counts” and Table S6: “Transcript Counts”.

Through this process 469 frameshifts in 387 genes were identified in CHM13 versus GENCODE v35. IDs of the frameshifted genes and transcripts can be found in Table S7: “Frameshifts”. An example is shown in Fig. S27.

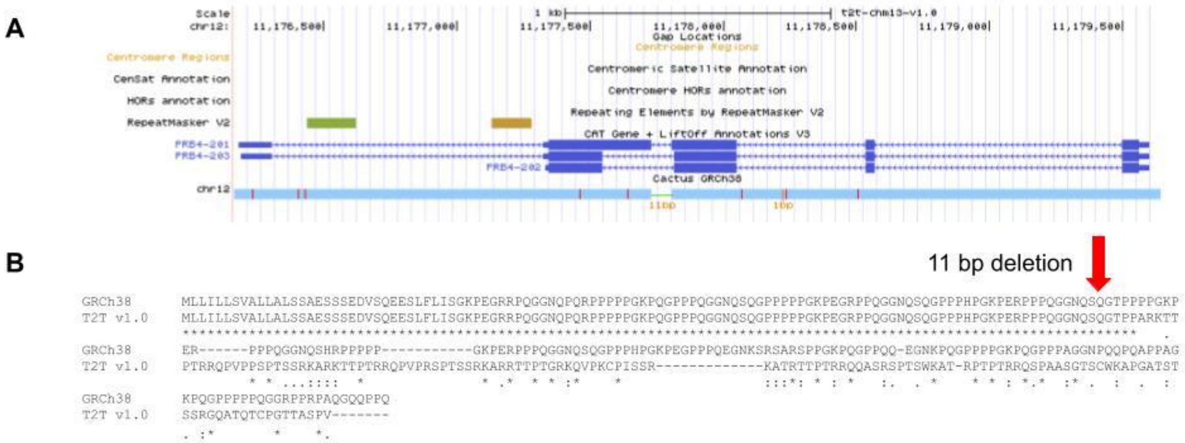


**Fig. S27: Frameshift in PRB4**. (A) Browser screenshot of the region. (B) Alignment of the predicted protein sequences.

From the GENCODE v35 annotation, 263 genes were missing from the v1.1 assembly gene set (63 protein-coding, 67 non-coding RNA, 94 pseudogenes, and 39 Immunoglobulin/ T-cell receptor gene segments). In terms of transcripts, 1,708 transcripts from GENCODE v35 annotation were missing from the v1.1 assembly annotation (829 protein-coding, 256 non-coding RNA, 109 pseudogenes, 39 Immunoglobulin/ T-cell receptor gene segments, and 475 other). IDs of the missing genes and transcripts can be found in, respectively, Table S8:

“Missing GENCODE genes” and Table S9: “Missing GENCODE transcripts”. There are a number of reasons why genes may not have been annotated by either the CAT or Liftoff pipelines. First, the gene may truly be absent in the v1.1 assembly (and potentially the CHM13 genome). Fig. S28 gives an example of *ORM1* gene, falling within a 6,762 bp deletion on Chr9 in the v1.1 assembly relative to GRCh38.

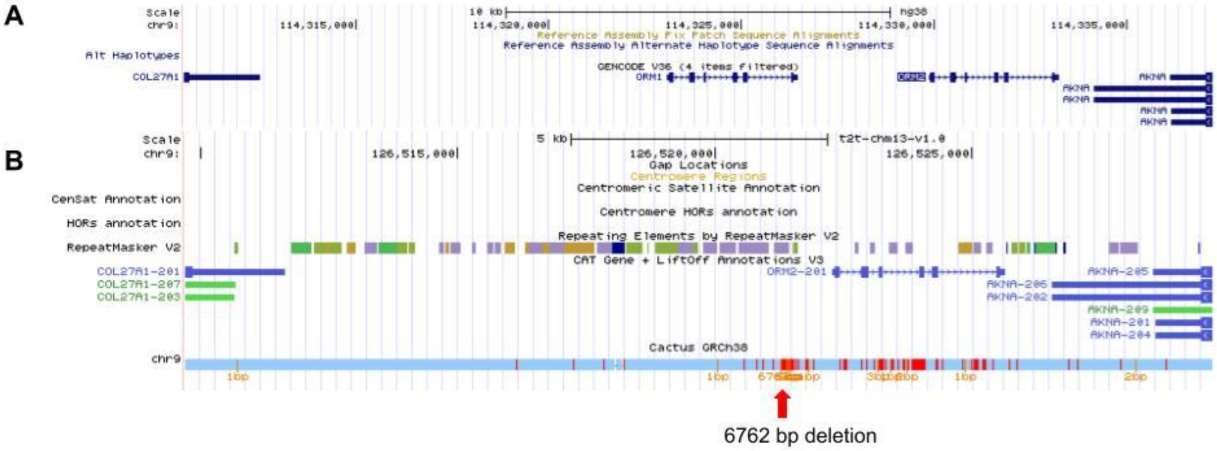


**Fig. S28: ORM1 gene deletion.** (A) A browser screenshot of GRCh38 showing a region on Chr9, with annotations for *ORM1* and *ORM2* genes. Upstream and downstream genes (*COL27A1* and *AKNA*) are included for context. (B) Browser screenshot of the corresponding region on CHM13 v1.0 assembly. *ORM2* is annotated; ORM1 falls within the 6,762 bp deletion, shown in the alignment track, and is thus missing.

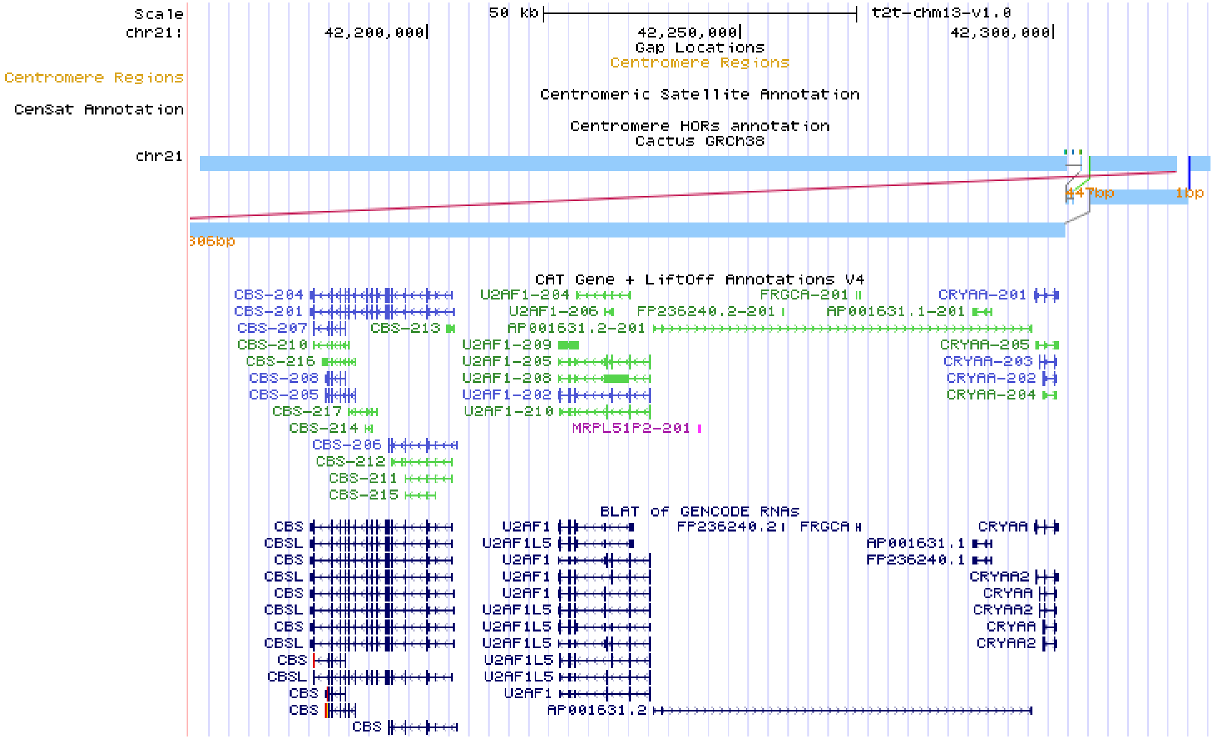

**Fig. S29**: **Gene annotation in a region of false duplication on GRCh38 in chr21 (coordinates chr21:42172000-42320000)**. The Cactus GRCh38 tracks show the alignment of the duplicated regions in GRCh38. The GENCODE BLAT track shows the multiple copies of the genes that all map to the same region in CHM13. The CAT+Liftoff annotation track shows how the annotation is resolved so that only one copy of each gene is included in the final annotation.

This is likely to happen with genes associated with segmental duplications within copy number variable regions, such as CT45/47 on Chromosome X which have previously been validated as correct in the CHM13 assembly (*23*). GENCODE genes may also not be annotated if alignments fail. With CAT, this can happen when the genomic sequence of CHM13 is unable to align (or aligns poorly) to GRCh38. The GENCODE transcripts may also fail to align to the assembly (done by TransMap in CAT, and Minimap2 in Liftoff). Lastly, GRCh38 may have assembly errors that create false duplications and 'missing' copies. For example, the region on Chromosome 21 shown in Fig. S29, has been flagged by prior studies (*68*, *69*) and is not consistent with copy-number variation seen in the human population (*8*).

Genes and transcripts exclusive to CHM13 are in Table S10: "Extra CHM13 Genes" and Table S11: "Extra CHM13 transcripts". The identity of the protein-coding genes are in Table S12: "Extra Paralog Identities".

### 10. Acrocentrics and rDNA analysis

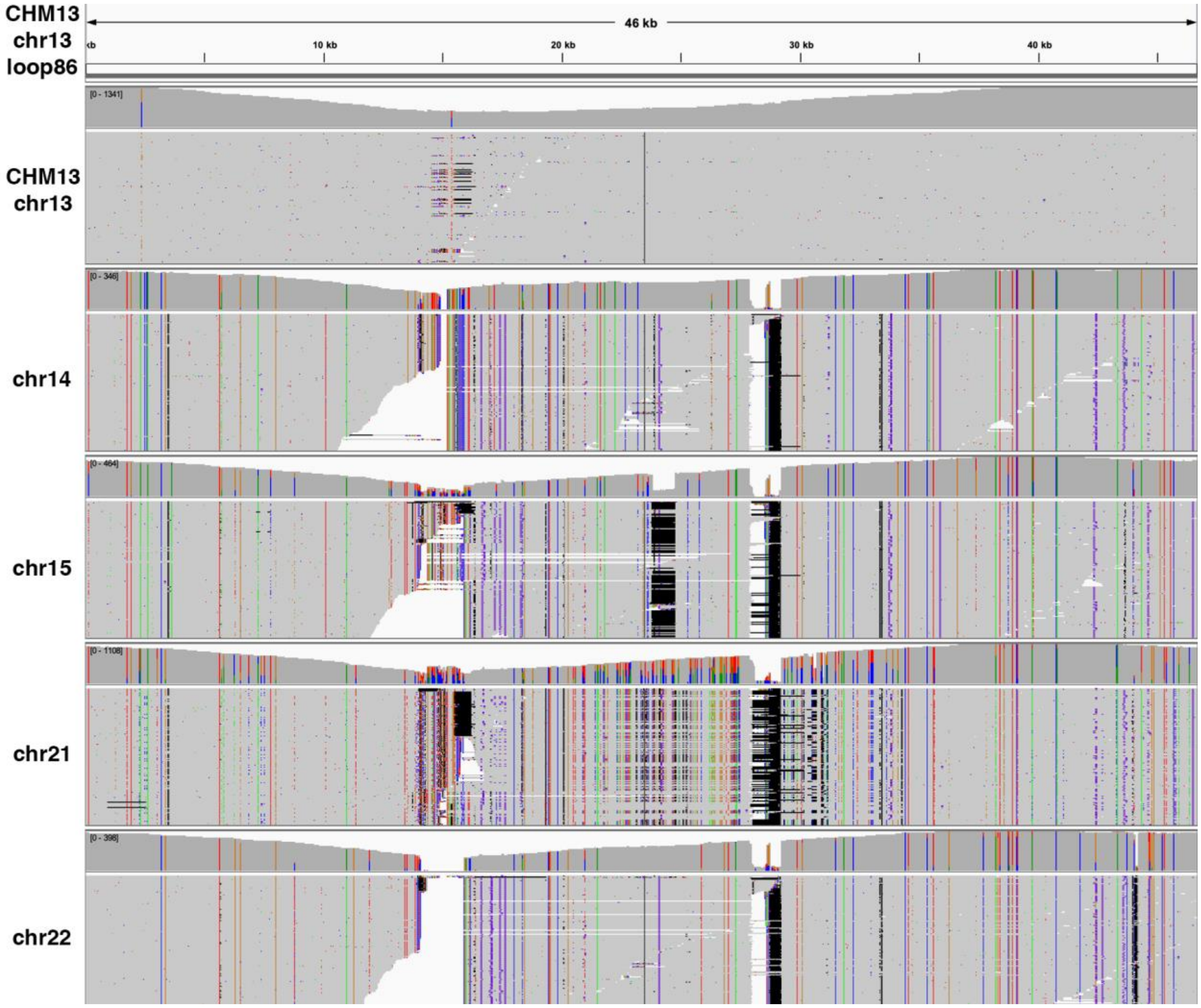


**Fig. S30: More variation exists between rDNA units on different chromosomes than within chromosomes.** CHM13 HiFi reads were binned by chromosome using the method described above in the section "Resolution of the rDNA arrays". Chromosome-specific HiFi reads were then aligned to the major morph on chromosome 13 ("loop86") using minimap2 with the `-ax map-pb` option and visualized in IGV. The alignments of chr13 reads versus the chr13 major morph are much cleaner than the alignments of reads from other chromosomes to chr13, demonstrating that the individual rDNA units on chr13 are more similar to one another than to those on other chromosomes. The major morphs on chromosomes 14, 15, 21, 22 show a similar pattern when this experiment is repeated.

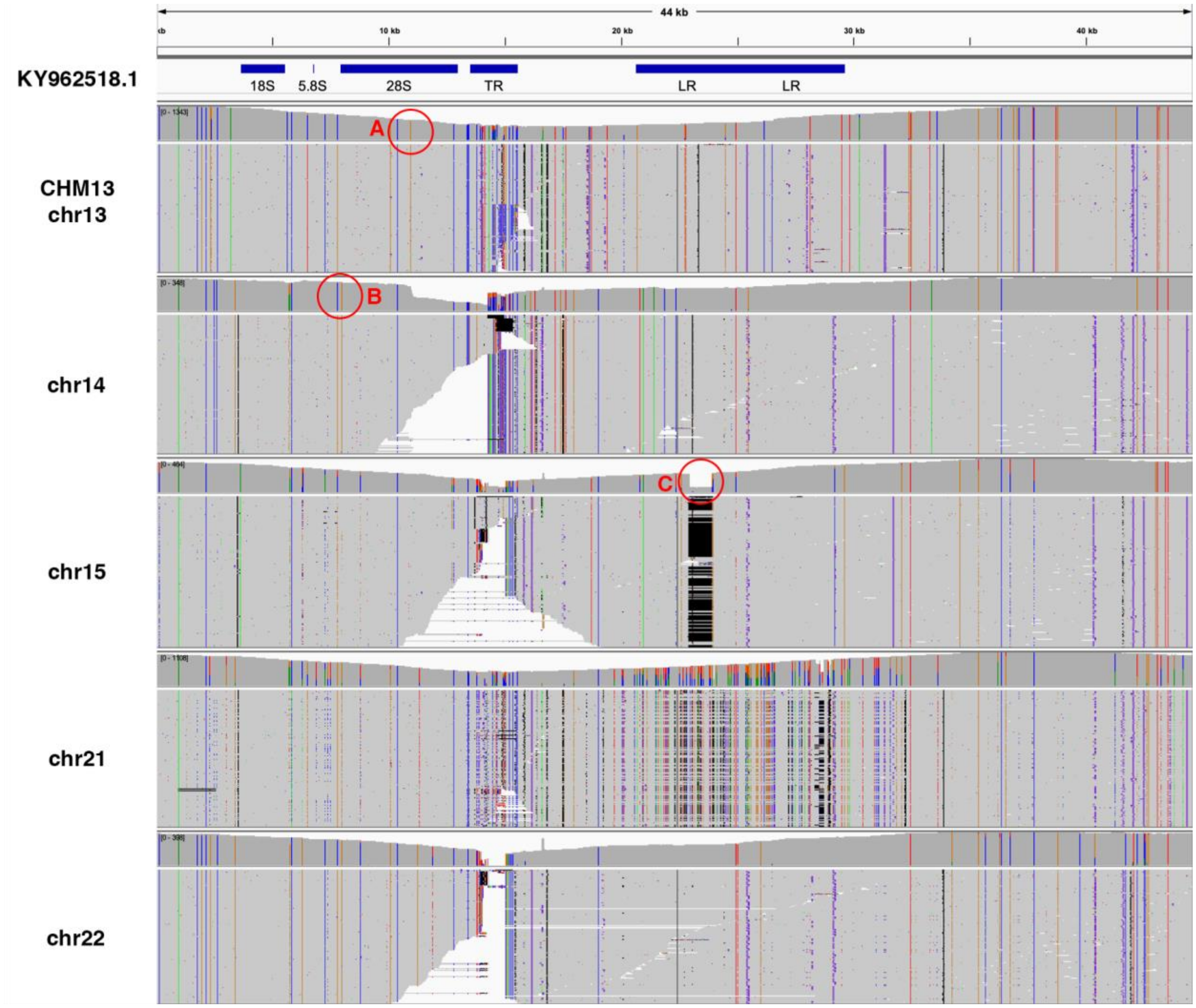


**Fig. S31: Some rDNA variants appear chromosome specific in CHM13.** CHM13 HiFi reads were binned by chromosome using the method described above in the section "Resolution of the rDNA arrays". Chromosome-specific HiFi reads were then aligned to a reference rDNA unit from GenBank (KY962518.1) using minimap2 with the `-ax map-pb` option and visualized in IGV. Key features of the rDNA unit, including the 18S, 5.8S, and 28S genes and the "TR" and "LR" repeats are annotated at the top. (A) An example SNP within the 28S is unique to CHM13 chr13 (brown variant). (B) Another example SNP is shared by chr14 and chr22 in CHM13, but absent from the reference and other CHM13 chromosomes. (C) A deletion within the LR is unique to the major morph on chr15 in CHM13, but the reference variant is also present at lower frequency in a minor morph.

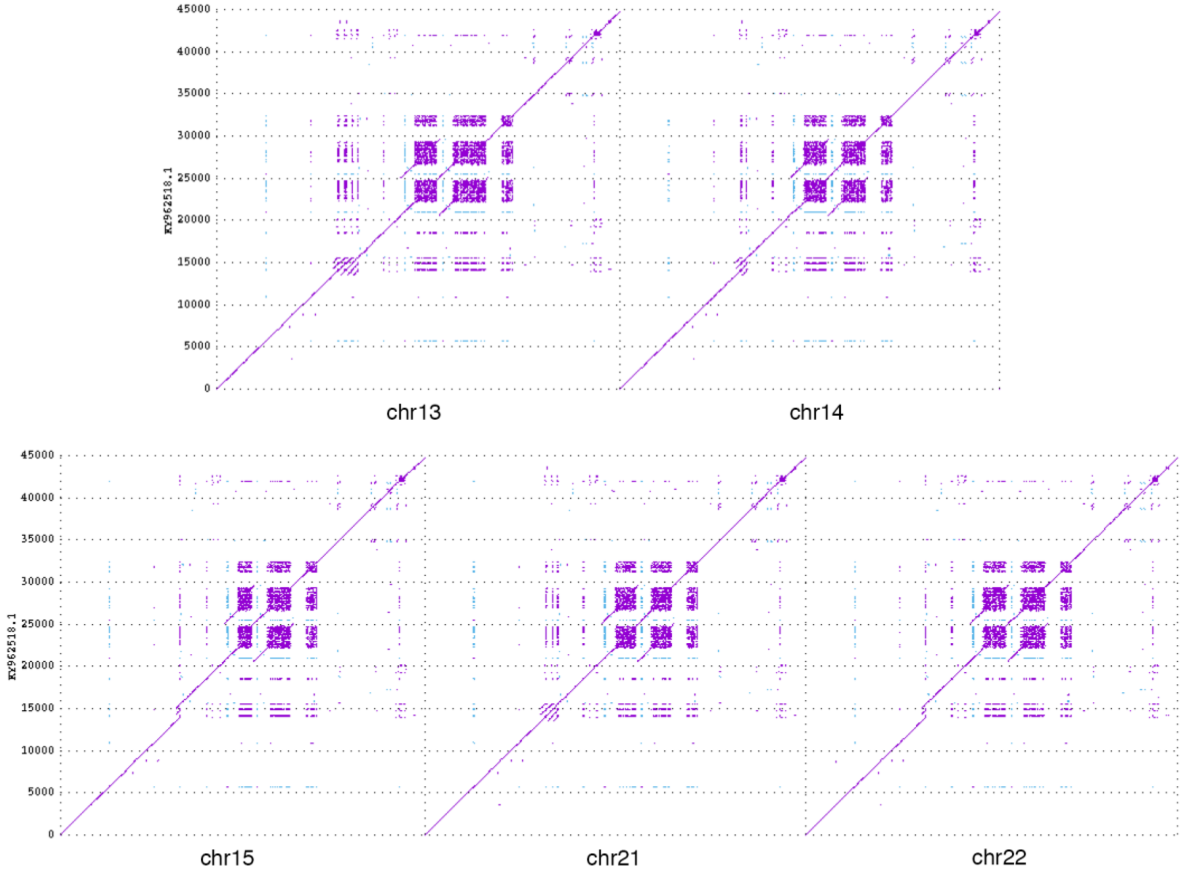


**Fig. S32: The major rDNA morphs on each chromosome are structurally unique.** MUMmer doplots are shown for exact matches of length 20 (forward matches in purple, reverse complement in blue). Chromosome morphs are on the x-axis and the KY962518.1 reference is on the y-axis. The “TR” tandem repeat at position 15 kbp is frequently variable between morphs, with a copy number of 4, 2, 1, 3, and 1 for the major morph on chromosomes 13, 14, 15, 21, and 22, respectively (the reference has 3 copies). The “LR” long repeat between positions 20 kbp and 30 kbp is another frequent source of variation between morphs, with the second copy expanded in chromosome 13 and the first copy collapsed in chromosome 15, relative to the reference. Most morphs can be distinguished based on the size of their TR and LR repeats.

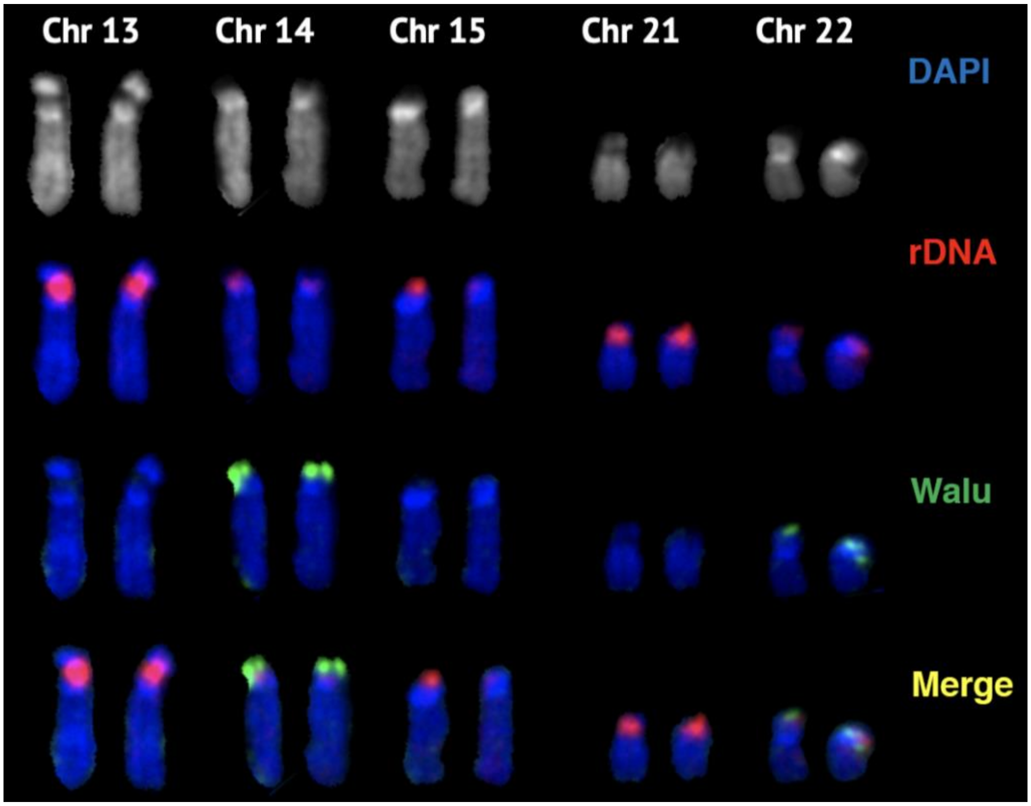


**Fig. S33: Repeat element FISH on acrocentric chromosomes.** Karyogram of acrocentric chromosomes from a CHM13 chromosome spread labeled by FISH with rDNA probe (orange) and the novel WaluSat probe (green). All acrocentric chromosomes were identified by morphology. WaluSat signal was detected on chromosomes 14 and 22, as predicted by the assembly.

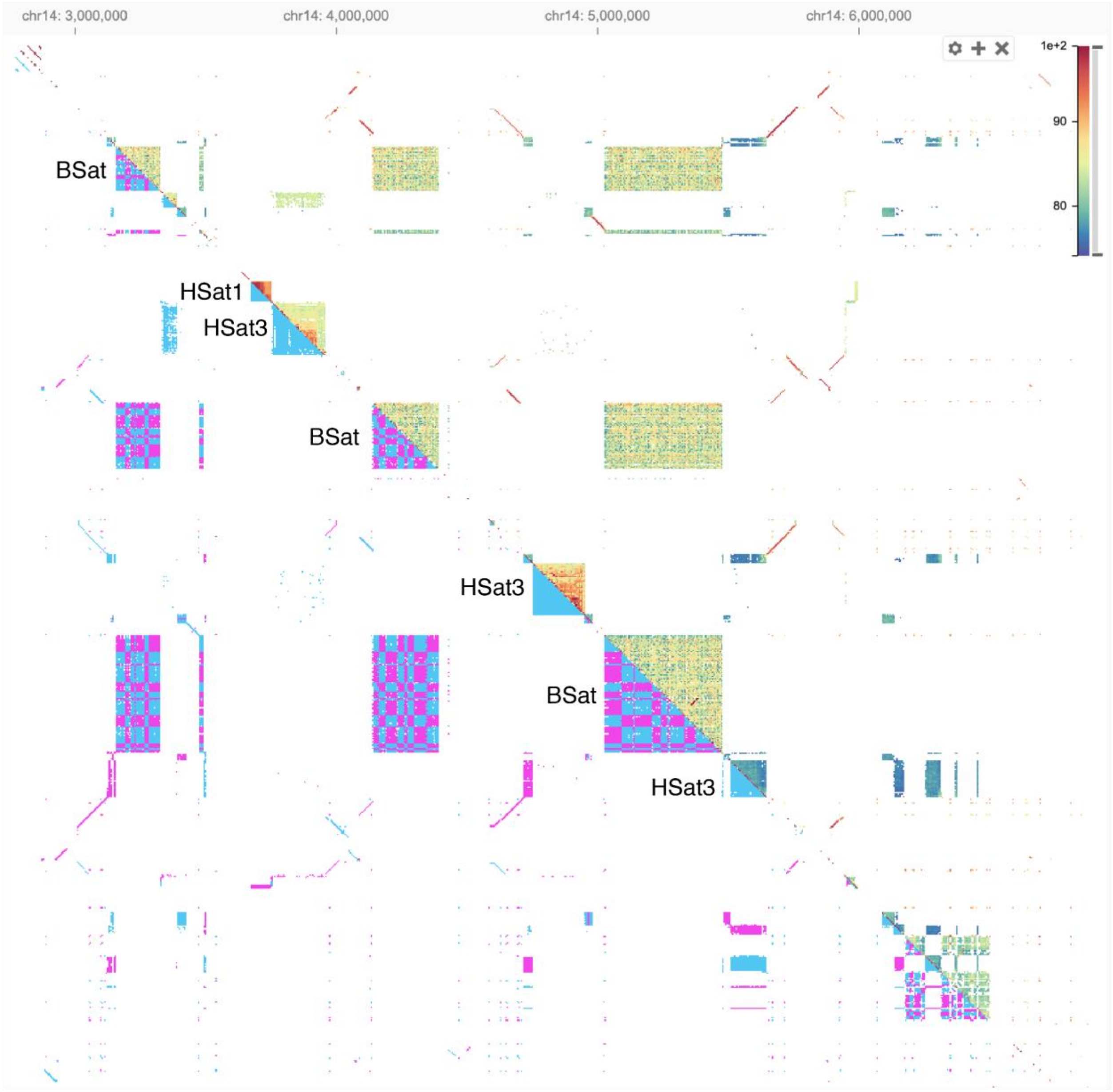


**Fig. S34: Beta satellite arrays on the acrocentric proximal short arms show a mosaic inversion structure.** A self-alignment dotplot of CHM13 Chromosome 14 (position 3–7 Mbp) is shown with the upper triangle displaying alignment similarity (as noted by the color scale) and the lower triangle displaying alignment orientation (forward in purple, reverse complement in blue). While the HSat1 and HSat3 tandem units only align in a forward orientation, the BSat arrays show a checkerboard pattern that is indicative of large inverted blocks of satellite sequence. Inversions are also evident within the alpha satellite monomer repeats at the bottom right of the plot. A similar pattern appears on the other acrocentric chromosomes, but only within the proximal satellite arrays and not the distal arrays.

### 11. Mappability

In an effort to compare the mappability of this assembly to the GRCh38 reference both a variety of sequencing technologies were compared (differing by read lengths and accuracy) and an aligner-independent strategy was implemented. First, alignments of different CHM13 sequencing read sets to v1.0 assembly (filtered by MAPQ >=60) were used to estimate the average length of perfectly matching segments for each sequencing technology. These characteristic perfect runs were 129 bp, 500 bp, and 33 bp for Illumina, HiFi, and ONT (Guppy v3.6.0 base-called), respectively. To assess the mappability of a genome by a particular read type we used GenMap (*70*) to find positions of unique k-mers in the genome, with k set to the technology's characteristic perfect run size. A sliding window equal to the read size was used to determine if a genomic region was "mappable". Any window with at least 1 kbp, or 33% of the read length for reads < 1 kbp, was considered "mappable". The resulting mappability statistics across different technologies and read lengths is summarized in Table S13.

**Table S13: Total length of unmappable regions across assemblies.**

| | GRCh38p13 | GRCh38p13 Q40 | CHM13 v1.0 |
|---|---|---|---|
| **Illumina 250 bp** | 223.01 (7.19%) | 220.21 (7.10%) | 200.35 (6.46%) |
| **HiFi 10 kbp** | 141.31 (4.56%) | 136.86 (4.41%) | 63.52 (2.05%) |
| **HiFi 20 kbp** | 136.34 (4.39%) | 128.17 (4.13%) | 35.12 (1.13%) |
| **HiFI 25 kbp** | 134.88 (4.35%) | 124.81 (4.03%) | 28.49 (0.92%) |
| **ONT 100 kbp** | 136.78 (4.41%) | 133.25 (4.30%) | 109.55 (3.53%) |
| **ONT 150 kbp** | 129.52 (4.18%) | 127.13 (4.10%) | 88.01 (2.84%) |
| **ONT 200 kbp** | 126.16 (4.07%) | 112.54 (3.95%) | 73.64 (2.38%) |

Total length of 'unmappable' regions (Mbp) and the fraction of the assembly they represent (using 3.1 Gbp as genome size). A region is considered unmappable by a particular technology if less than 1 kbp or 33% of the read length (whichever is smaller) of sequence is represented by unique k-mers (where k equals the technology's characteristic perfect run size). Note that regions with gaps can be classified as mappable given a unique sequence at their boundaries, allowing reads to extend into the gap. To estimate the potential effects of consensus quality on mappability, we added random errors to GRCh38 at the frequency of 0.01% (Q40) (with 45% of the errors being insertion, 45% -- deletions, and 10% -- substitutions). We observe that CHM13 v1.0 assembly is expected to be more mappable with long read technologies than GRCh38, despite having a higher repeat content. The increased mappability is not due to consensus quality as GRCh38 with added errors is still less mappable than CHM13 v1.0.

To find positions of multi-copy *k*-mers in the CHM13 v1.0 and GRCh38 for a wide range of *k* (powers of 2+1 from 33 to 131,073 bp) GenMap was used. At *k*=131,073 bp, all CHM13v1.0 *k*-mers were unique and at *k*=65,537 the only repetitive *k*-mers were located within the Chromosome 9 HSat3 array draft reconstruction (Fig. S24). Contrastingly, GRCh38 had megabases of non-unique regions spread across all chromosomes at both *k*=65,537 bp and *k*=131,037 bp, even when gap bases were not considered. Fig. S35 presents several *k*-mer based repeat statistics across different genomes and values of *k*. Note that rDNA repeat units were excluded from *k*-mer analysis since these are model-based in the current assembly (see "Resolution of the rDNA arrays").

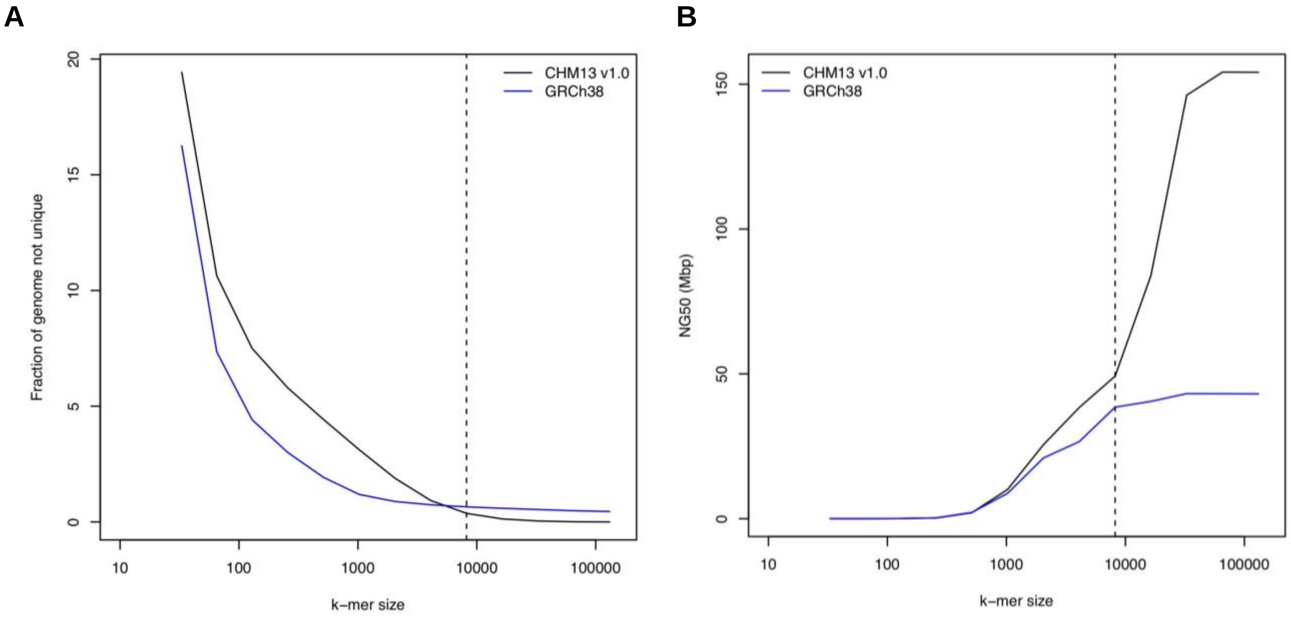


**Fig. S35: *k*-mer analysis of CHM13v1.0 and GRCh38.** (A) Fraction of genome (in percent) spanned by multi-copy *k*-mers (y axis) vs. *k*-mer size (x axis). CHM13v1.0 has higher repeat content at lower *k*-mer sizes, but as *k* exceeds 8 kbp (vertical line) GRCh38 overtakes CHM13v1.0 in repetitive sequence fraction. This indicates that individual instances of large genomic repeats might not have been accurately resolved within GRCh38 , but were rather copy/pasted or compiled from model sequence, effectively homogenizing repeat units. (B) NG50 of genomic fragments after breaking at the start/end of every stretch of non-unique *k*-mers (y axis) vs. *k*-mer size (x axis). Non-unique stretches formed their own contigs. Note that due to the co-localization of the repeats within the genome, CHM13v1.0 is more “assemblable” than GRCh38 at all *k*-mer sizes over 1 kbp despite a larger fraction of it being repetitive. Substantial difference between CHM13v1.0 and GRCh38 statistics at higher values of *k* (over 8 kbp) indicates that a haploid human genome is more easily assembled by highly accurate sequencing reads (of 10 kbp and longer) than one would presume from GRCh38 reference analysis. This is consistent with the observation that projections relying on GRCh38 substantially under-estimated continuity of recent HiFi-based assemblies (*13*, *71*).

### 12. *FRG1* Analysis

All *FRG1* paralogs were identified in CHM13v1.0 and GRCh38 and are listed in Table S14.

**Table S14: Identified *FRG1* SD paralogs in CHM13 v1.0 and GRCh38.**

| Reference | Chr. | Start | End | Gene in SD | Strand |
|---|---|---|---|---|---|
| CHM13v1.0 | chr4 | 193304806 | 193328432 | *FRG1* | + |
| CHM13v1.0 | chr14 | 9482463 | 9564392 | *FRG1GP 2* | + |
| CHM13v1.0 | chr15 | 15030674 | 15117873 | *FRG1BP 9* | + |
| CHM13v1.0 | chr22 | 12204955 | 12286842 | *FRG1GP* | + |
| CHM13v1.0 | chr9 | 42281940 | 42361070 | *FRG1HP* | - |
| CHM13v1.0 | chr9 | 77150969 | 77230137 | *AL353763.2* | - |
| CHM13v1.0 | chr13 | 13539901 | 13624992 | *FRG1BP 5* | - |
| CHM13v1.0 | chr14 | 6505624 | 6592011 | *FRG1BP 6* | + |
| CHM13v1.0 | chr20 | 31531577 | 31613298 | *FRG1DP* | - |
| CHM13v1.0 | chr21 | 10617841 | 10702911 | *FRG1BP 4* | - |
| CHM13v1.0 | chr21 | 2600799 | 2689988 | *FRG1BP 3* | - |
| CHM13v1.0 | chr13 | 12171267 | 12264520 | *FRG1BP 8* | + |
| CHM13v1.0 | chr14 | 7817618 | 7910951 | *FRG1BP 10* | - |
| CHM13v1.0 | chr15 | 15404707 | 15484609 | *FRG1KP 4* | + |
| CHM13v1.0 | chr21 | 9297091 | 9390323 | *FRG1BP 7* | + |
| CHM13v1.0 | chr9 | 47641268 | 47724493 | *FRG1JP* | + |
| CHM13v1.0 | chr22 | 4339765 | 4408405 | *FRG1BP 2* | - |
| CHM13v1.0 | chr14 | 10057678 | 10125075 | *FRG1FP 2* | - |
| CHM13v1.0 | chr20 | 29521044 | 29589686 | *FRG1CP* | + |
| CHM13v1.0 | chr22 | 12726527 | 12793932 | *FRG1FP* | - |
| CHM13v1.0 | chr20 | 31098413 | 31169152 | *FRG1EP* | - |
| CHM13v1.0 | chr20 | 30385905 | 30449933 | *FRG1BP* | - |
| CHM13v1.0 | chr15 | 17211534 | 17232403 | no transcript | - |
| CHM13v1.0 | chr9 | 40738195 | 40747139 | *FRG1KP* | + |
| CHM13v1.0 | chr9 | 80141785 | 80150732 | *FRG1KP 2* | + |
| CHM13v1.0 | chr9 | 42074670 | 42083617 | *FRG1HP* | - |
| CHM13v1.0 | chr9 | 80385461 | 80394406 | *FRG1KP 3* | - |
| GRCh38 | chr4 | 189940872 | 189963192 | *FRG1* | + |
| GRCh38 | chrUn_GL000219v1 | 71519 | 160661 | AL592183.1 | - |
| GRCh38 | chr9 | 40933942 | 41013034 | *FRG1HP* | + |
| GRCh38 | chr22 | 12550526 | 12632365 | *FRG1GP* | + |
| GRCh38 | chr9 | 63831480 | 63916428 | *FRG1JP* | - |
| GRCh38 | chr20 | 28575029 | 28643629 | *FRG1CP* | - |
| GRCh38 | chr20 | 29045065 | 29108496 | *FRG1DP* | + |
| GRCh38 | chr20 | 29493951 | 29536889 | *FRG1EP* | - |
| GRCh38 | chr20 | 30346078 | 30399907 | *FRG1BP* | + |
| GRCh38 | chr22 | 10933716 | 10966724 | *FRG1FP* | - |
| GRCh38 | chr9_KI270720v1_random | 14808 | 39050 | no transcript | - |
| GRCh38 | chr20 | 29474039 | 29493657 | *FRG1EP* | - |
| GRCh38 | chr9_KI270718v1_random | 22210 | 31135 | no transcript | + |
| GRCh38 | chr9 | 65646335 | 65653950 | *FRG1KP* | - |
| GRCh38 | chr9 | 41211431 | 41219057 | *FRG1HP* | + |

Annotated segmental duplications of the ancestral *FRG1* locus (chr4:189940872-189963192, GRCh38, “Segmental Dups” track (*72*); chr4:193304806-193328432, CHM13v1.0, SEDEF track (*59*)) were intersected with annotated genes (GENCODEv36 (*66*) for GRCh38 and CAT+Liftoff v4 for CHM13v1.0) to identify *FRG1* paralogs.

For CHM13v1.0 paralogs, sequences were aligned using MAFFT v7 (*73*) and analyzed using MEGA X (*74*). A phylogenetic tree was inferred using the Neighbor-Joining method (*75*) with 500 bootstrap replicates. The evolutionary distances were computed using the Kimura 2-parameter method (*76*). This analysis involved 27 nucleotide sequences. All ambiguous positions were removed for each sequence pair (pairwise deletion option). There were a total of 128,204 positions in the final dataset (Fig. S36A). Additionally, we applied a maximum likelihood method using RAxML (*77*) and the general time reversible+G model (*78*) as matched by jModelTest2. RAxML was run on the PTHREADS version and rapid bootstrap approximation to generate 100 bootstrap replicates. Consensus trees were then generated in Geneious3 and visualized in FigTree4 (Fig. S36B). The tree is drawn to scale, with branch lengths measured in the number of substitutions per site. Despite some of the paralog branches differing between approaches, both trees strongly supported that the *FRG1BP4~10* paralogs shared the same ancestral origin with *FRG1DP* (Fig. S36) and share identity >96.5% (Table S15).

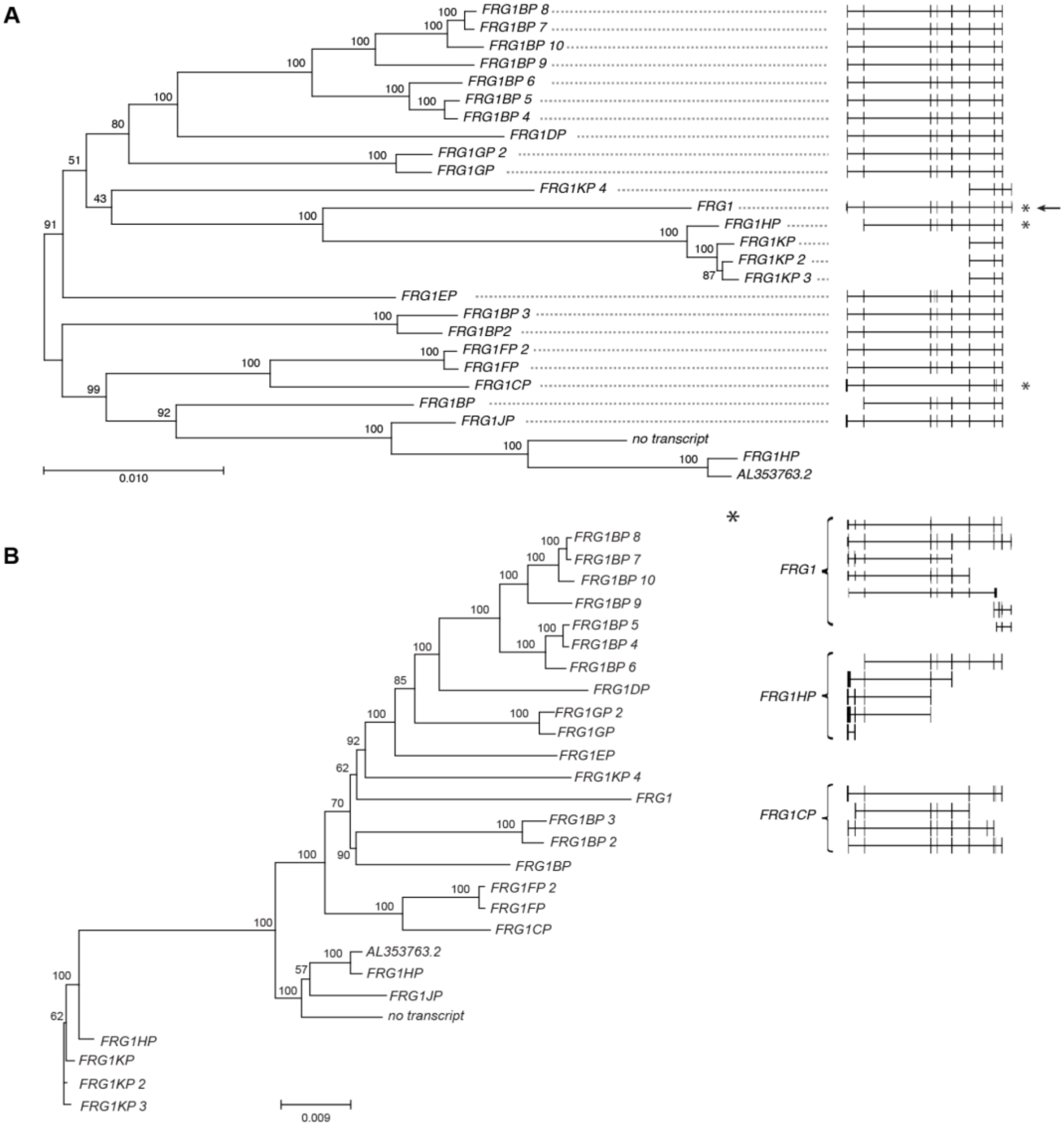


**Fig. S36. Phylogenetic tree of the *FRG1* paralogs.** (A) Tree inferred from the Neighbor-joining method. The tree is drawn to scale, with branch lengths in the same units as those of the evolutionary distances used to infer the phylogenetic tree. Units are the number of base substitutions per site. Numbers shown next to the branches are the percentage of replicate trees clustered together in the bootstrap test. Genes annotated with multiple transcripts are marked with *. (B) Tree inferred from the maximum likelihood comparison method with 100 bootstrap replicates. Both trees share similar insights on the branches close to *FRG1DP*.

**Table S15. Estimates of evolutionary divergence between sequences.**

| | FRG1 | FRG1 BP | FRG1 BP_2 | FRG1 BP_3 | FRG1 BP_4 | FRG1 BP_5 | FRG1 BP_6 | FRG1 BP_7 | FRG1 BP_8 | FRG1 BP_9 | FRG1 BP_10 | FRG1 CP | FRG1 DP | FRG1 EP | FRG1F P | FRG1F P_2 | FRG1 GP | FRG1 GP_2 | FRG1 HP | FRG1 HP (v2) | FRG1J P | FRG1 KP | FRG1 KP_2 | FRG1 KP_3 | FRG1 KP_4 | AL353 763.2 |
|---|---|---|---|---|---|---|---|---|---|---|---|---|---|---|---|---|---|---|---|---|---|---|---|---|---|---|
| FRG1 | - | | | | | | | | | | | | | | | | | | | | | | | | | |
| FRG1 BP | 94.7% | - | | | | | | | | | | | | | | | | | | | | | | | | |
| FRG1 BP_2 | 94.3% | 95.8% | - | | | | | | | | | | | | | | | | | | | | | | | |
| FRG1 BP_3 | 94.3% | 95.8% | 99.4% | - | | | | | | | | | | | | | | | | | | | | | | |
| FRG1 BP_4 | 94.3% | 95.8% | 95.4% | 95.2% | - | | | | | | | | | | | | | | | | | | | | | |
| FRG1 BP_5 | 94.3% | 95.8% | 95.4% | 95.2% | 99.9% | - | | | | | | | | | | | | | | | | | | | | |
| FRG1 BP_6 | 94.2% | 95.8% | 95.4% | 95.2% | 99.4% | 99.4% | - | | | | | | | | | | | | | | | | | | | |
| FRG1 BP_7 | 94.0% | 95.7% | 95.3% | 95.2% | 98.3% | 98.3% | 98.3% | - | | | | | | | | | | | | | | | | | | |
| FRG1 BP_8 | 94.0% | 95.7% | 95.3% | 95.1% | 98.3% | 98.3% | 98.3% | 99.9% | - | | | | | | | | | | | | | | | | | |
| FRG1 BP_9 | 94.1% | 95.7% | 95.3% | 95.1% | 98.2% | 98.2% | 98.3% | 98.9% | 98.9% | - | | | | | | | | | | | | | | | | |
| FRG1 BP_10 | 94.0% | 95.6% | 95.2% | 95.1% | 98.3% | 98.2% | 98.3% | 99.7% | 99.7% | 98.9% | - | | | | | | | | | | | | | | | |
| FRG1 CP | 94.4% | 95.8% | 95.4% | 95.3% | 95.4% | 95.4% | 95.4% | 95.3% | 95.2% | 95.3% | 95.2% | - | | | | | | | | | | | | | | |
| FRG1 DP | 93.9% | 95.6% | 95.0% | 94.9% | 96.7% | 96.7% | 96.6% | 96.5% | 96.5% | 96.5% | 96.5% | 94.9% | - | | | | | | | | | | | | | |
| FRG1 EP | 94.4% | 96.1% | 95.8% | 95.8% | 96.1% | 96.1% | 96.1% | 96.1% | 96.0% | 96.0% | 96.0% | 95.5% | 96.1% | - | | | | | | | | | | | | |
| FRG1F P | 94.4% | 95.8% | 95.5% | 95.4% | 95.5% | 95.5% | 95.5% | 95.3% | 95.3% | 95.4% | 95.3% | 97.9% | 95.0% | 95.5% | - | | | | | | | | | | | |
| FRG1F P_2 | 94.4% | 95.8% | 95.5% | 95.4% | 95.5% | 95.5% | 95.5% | 95.3% | 95.3% | 95.4% | 95.3% | 97.9% | 95.0% | 95.5% | 99.9% | - | | | | | | | | | | |
| FRG1 GP | 94.5% | 95.8% | 95.4% | 95.2% | 96.4% | 96.4% | 96.4% | 96.5% | 96.5% | 96.5% | 96.4% | 95.5% | 96.3% | 96.0% | 95.5% | 95.5% | - | | | | | | | | | |
| FRG1 GP_2 | 94.4% | 95.8% | 95.3% | 95.2% | 96.4% | 96.4% | 96.4% | 96.5% | 96.5% | 96.5% | 96.4% | 95.4% | 96.3% | 96.0% | 95.5% | 95.5% | 99.6% | - | | | | | | | | |
| FRG1 HP | 94.3% | 96.5% | 95.7% | 95.7% | 95.6% | 95.5% | 95.5% | 95.5% | 95.5% | 95.5% | 95.4% | 96.4% | 95.2% | 95.7% | 96.4% | 96.4% | 95.6% | 95.6% | - | | | | | | | |
| FRG1 HP(v2) | 95.7% | 94.8% | 94.5% | 94.5% | 94.8% | 94.7% | 94.7% | 94.7% | 94.6% | 94.6% | 94.6% | 94.7% | 94.2% | 95.3% | 94.6% | 94.7% | 94.5% | 94.6% | 72.0% | - | | | | | | |
| FRG1J P | 94.5% | 96.4% | 95.6% | 95.5% | 95.4% | 95.4% | 95.4% | 95.4% | 95.4% | 95.4% | 95.3% | 96.1% | 95.1% | 95.7% | 96.2% | 96.2% | 95.5% | 95.5% | 98.3% | 96.2% | - | | | | | |
| FRG1 KP | 95.7% | 94.8% | 94.4% | 94.3% | 94.7% | 94.7% | 94.6% | 94.6% | 94.6% | 94.6% | 94.5% | 94.5% | 94.1% | 95.2% | 94.5% | 94.5% | 94.5% | 94.5% | 71.4% | 99.5% | 96.2% | - | | | | |
| FRG1 KP_2 | 95.8% | 94.8% | 94.4% | 94.4% | 94.8% | 94.7% | 94.6% | 94.6% | 94.6% | 94.6% | 94.5% | 94.6% | 94.1% | 95.2% | 94.5% | 94.6% | 94.5% | 94.5% | 71.2% | 99.6% | 96.2% | 99.8% | - | | | |
| FRG1 KP_3 | 95.7% | 94.7% | 94.4% | 94.3% | 94.7% | 94.6% | 94.6% | 94.6% | 94.6% | 94.6% | 94.5% | 94.5% | 94.1% | 95.2% | 94.5% | 94.5% | 94.4% | 94.5% | 71.4% | 99.6% | 96.2% | 99.8% | 99.9% | - | | |
| FRG1 KP_4 | 94.1% | 95.5% | 95.1% | 95.0% | 95.2% | 95.2% | 95.2% | 95.1% | 95.1% | 95.1% | 95.1% | 95.2% | 95.1% | 95.2% | 95.2% | 95.2% | 96.0% | 96.0% | 95.4% | 94.6% | 95.3% | 94.6% | 94.6% | 94.5% | - | |
| AL353 763.2 | 94.3% | 96.5% | 95.7% | 95.7% | 95.5% | 95.5% | 95.5% | 95.5% | 95.5% | 95.5% | 95.5% | 96.4% | 95.2% | 95.7% | 96.4% | 96.4% | 95.6% | 95.6% | 99.7% | 72.2% | 98.3% | 71.6% | 71.5% | 71.6% | 95.4% | - |
| no_tra nscript | 93.6% | 96.8% | 95.6% | 95.6% | 95.4% | 95.4% | 95.4% | 95.2% | 95.2% | 95.3% | 95.2% | 95.3% | 95.1% | 95.4% | 95.2% | 95.2% | 95.5% | 95.5% | 98.3% | 77.3% | 97.8% | 76.9% | 76.6% | 76.9% | 95.1% | 98.3% |

Pairwise identity for all *FRG1* paralogs in the CHM13 assembly (see Table S14 for gene locations). The number of base substitutions per site from between sequences was calculated and percent identity calculated by taking one minus this value. Analyses were conducted using the Kimura 2-parameter model and MEGA X as for the tree in Fig. S36A. Similarity between *FRG1DP* and *FRG1BP4~10* is highlighted in blue.

Transcript abundance was estimated in transcripts per million (TPM) with Salmon v1.3.0 (*79*), using the CHM13 transcriptome and the v1.0 assembly as a “decoy” sequence to account for reads mapping to unannotated sequences. TPM values were aggregated to the gene level using the R package tximport (*80*). In addition, TPM values per gene were calculated from alignments of RNA-seq and Iso-seq data, filtered on singly unique markers in the CHM13 assembly. Full details are in (*44*). The *FRG1* paralogs show varying levels of expression (Table S16).

#### Table S16: Expression analysis.

| Gene | Cen/ Acro | Chr | Start | End | Gene (GENCODEv35) | Ensembl ID | T2T Annotation ID | Salmon | RNA-seq (k=21) | RNA-seq (k=51) | PRO-seq (k=21) | PRO-seq (k=51) | Iso-seq |
|---|---|---|---|---|---|---|---|---|---|---|---|---|---|
| FRG1 | - | chr4 | 193305949 | 193328281 | FRG1 | ENSG00000109536 | CHM13_G0043626 | 31.98 | 7.95 | 7.47 | 10.14 | 10.09 | 4.43 |
| FRG1CP | Cen | chr20 | 29561831 | 29600874 | FRG1CP | ENSG00000282826 | CHM13_G0034693 | 30.94 | 12.65 | 13.45 | 9.03 | 13.61 | 24.19 |
| FRG1HP | Cen | chr9 | 42076623 | 42302738 | FRG1HP | ENSG00000276291 | CHM13_G0055936 | 12.02 | 0.31 | 0.37 | 0.39 | 0.57 | 0.49 |
| FRG1BP | Cen | chr20 | 30394345 | 30416241 | FRG1BP | ENSG00000149531 | CHM13_G0034700 | 10.22 | 3.61 | 3.78 | 4.95 | 6.35 | 4.51 |
| FRG1BP 2 | Acro | chr22 | 4345924 | 4367793 | FRG1BP | ENSG00000149531 | LOFF_G0001769 | 4.75 | 2.58 | 2.67 | 1.67 | 2.25 | 13.06 |
| FRG1FP | Cen | chr22 | 12732209 | 12754159 | FRG1FP | ENSG00000283047 | CHM13_G0036432 | 3.08 | 0.22 | 0.21 | 0.26 | 0.42 | 5.50 |
| FRG1BP 4 | Cen | chr21 | 10623578 | 10645664 | FRG1BP | ENSG00000149531 | LOFF_G0001736 | 1.21 | 0.55 | 0.54 | 0.65 | 1.08 | 2.98 |
| FRG1GP 2 | Cen | chr14 | 9534454 | 9558648 | FRG1GP | ENSG00000283023 | CHM13_G0015651 | 1.18 | 0.27 | 0.27 | 0.22 | 0.58 | 0 |
| FRG1BP 3 | Acro | chr21 | 2606958 | 2628801 | FRG1BP | ENSG00000149531 | LOFF_G0001639 | 1.04 | 0.70 | 0.66 | 1.99 | 2.47 | 6.03 |
| FRG1KP 2 | Cen | chr9 | 80143074 | 80148774 | FRG1KP | ENSG00000280286 | LOFF_G0002852 | 0.99 | 0 | 0 | 0 | 0 | 0 |
| FRG1BP 10 | Cen | chr14 | 7824728 | 7846791 | FRG1BP | ENSG00000149531 | LOFF_G0000730 | 0.78 | 0.15 | 0.27 | 0.50 | 0.54 | 1.49 |
| FRG1JP | Cen | chr9 | 47695018 | 47722534 | FRG1JP | ENSG00000215548 | CHM13_G0056023 | 0.78 | 0.90 | 0.91 | 3.79 | 2.69 | 1.60 |
| FRG1BP 5 | Cen | chr13 | 13545638 | 13567729 | FRG1BP | ENSG00000149531 | LOFF_G0000580 | 0.65 | 0.15 | 0.15 | 0.50 | 1.00 | 5.47 |
| FRG1EP | Cen | chr20 | 31112410 | 31129441 | FRG1EP | ENSG00000282995 | CHM13_G0034716 | 0.57 | 0.79 | 0.81 | 1.48 | 2.21 | 6.45 |
| FRG1BP 9 | Cen | chr15 | 15088679 | 15110769 | FRG1BP | ENSG00000149531 | LOFF_G0000878 | 0.54 | 0.92 | 1.26 | 0.23 | 0.52 | 0 |
| FRG1DP | Cen | chr20 | 31537341 | 31559718 | FRG1DP | ENSG00000282870 | CHM13_G0034724 | 0.33 | 0.82 | 0.81 | 3.16 | 3.79 | 4.42 |
| FRG1BP 8 | Cen | chr13 | 12235362 | 12257433 | FRG1BP | ENSG00000149531 | LOFF_G0000552 | 0.13 | 0.03 | 0.03 | 0.06 | 0.04 | 0 |
| FRG1KP 4 | Cen | chr15 | 15473166 | 15478976 | FRG1KP | ENSG00000280286 | LOFF_G0000886 | 0.09 | 0.07 | 0.07 | 0 | 0.28 | 0 |
| FRG1KP | Cen | chr9 | 40739484 | 40745181 | FRG1KP | ENSG00000280286 | CHM13_G0055909 | 0 | 0 | 0 | 0 | 0 | 0 |
| FRG1BP 6 | Cen | chr14 | 6562833 | 6584916 | FRG1BP | ENSG00000149531 | LOFF_G0000705 | 0 | 0 | 0 | 0 | 0 | 0 |
| FRG1FP 2 | Cen | chr14 | 10063361 | 10085312 | FRG1FP | ENSG00000283047 | LOFF_G0000759 | 0 | 0 | 0 | 0 | 0.09 | 0 |
| FRG1BP 7 | Cen | chr21 | 9361171 | 9383233 | FRG1BP | ENSG00000149531 | LOFF_G0001708 | 0 | 0 | 0 | 0.03 | 0 | 0.50 |
| FRG1GP | Cen | chr22 | 12256920 | 12281097 | FRG1GP | ENSG00000283023 | LOFF_G0001867 | 0 | 0.81 | 0.86 | 1.64 | 2.67 | 8.18 |
| FRG1KP 3 | Cen | chr9 | 80387420 | 80393118 | FRG1KP | ENSG00000280286 | LOFF_G0002871 | 0 | 0 | 0 | 0 | 0 | 0 |

The table lists TPM, colored from white to red by abundance. ‘Gene’: The gene name in Fig. 4. ‘Cen/Acro’, ‘Chr’, ‘Start’, ‘End’: the location of the gene (centromeric satellite or acrocentric) along with the chromosome and coordinates in the CHM13 v1.0 assembly. ‘Gene (GENCODEv35)’: The gene name of the closest GENCODEv35 gene. ‘Ensembl ID’: closest ID in Ensembl. ‘T2T Annotation ID’: the ID of the annotated gene in the CHM13 v1.0 assembly. The remaining columns show abundance estimated from Salmon and short-reads aligned with unique 21 and 51-mer markers along with IsoSeq reads.

### 13. HG002 ChrX Assembly and Validation

#### Whole genome assembly

The same assembly pipeline was used (see “Whole genome assembly” section) to reconstruct Chromosome X from the GIAB (*81*) HG002 cell line (*82*). The HPRC (https://humanpangenome.org/hg002/) HiFi libraries (see “Data Availability”) were used to build a string graph and ONT UL data (*52*) to manually resolve tangles. There is a GA-rich region in the p-arm pseudoautosomal region (PAR) which we have not yet patched and so the current assembly ends approximately 2 Mbp short of the p-arm telomeric sequence.

#### Molecular Methods

##### Pulsed field gel electrophoresis

Alpha satellite array sizes were estimated by PFGE and Southern blotting using established methods (*23*, *83*). High molecular weight DNA from $10^7$-$10^8$ was embedded in 1% low melting point agarose plugs and digested with restriction enzymes that cut infrequently within alpha satellite DNA, releasing the DXZ1 array as one or a few large fragments. HMW DNA in one-half of an agarose plug was digested overnight with 20U of enzyme and run on a 1% agarose gel. Saccharomyces cerevisiae and Hansenula wingei chromosomes embedded in agarose were used as size standards (Bio-Rad CHEF DNA Size Markers). Gels were run at 3V/cm for 48-51 hours at 14°C in 1X TAE buffer, using switch times of 250 seconds (initial) – 900 seconds (final). CHM13, containing a previously sized DXZ1 array, was used as a control (*23*) (Fig. S37).

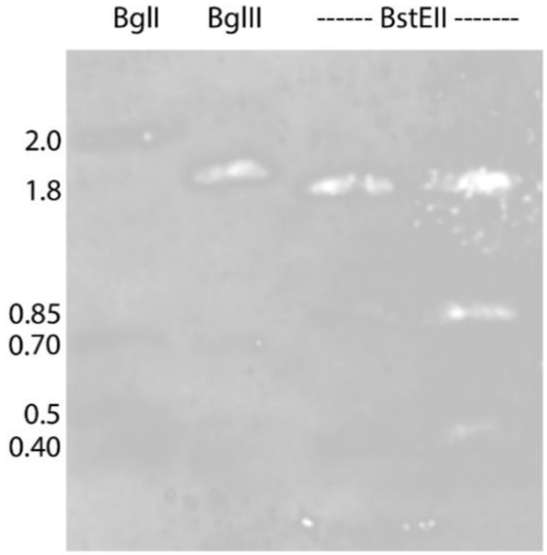


**Fig. S37: PFGE digestion of DXZ1 confirms assembly size.** Average estimates were 3.2 Mbp for BglI (2.0 + 0.7 + 0.5, n=3); 3.05 Mbp for BglII (1.9 + 0.7 + 0.45, n=1); and 3.05 Mbp for BstEII (1.8 + 0.85 + 0.40, n=4). The digest sizes were consistent with an in-silico digest of the assembly.

#### Southern blotting

After electrophoresis, gels were stained with ethidium bromide and imaged using a UV light source. Gels were depurinated with 0.25 M HCl for 12 minutes at room temperature, then washed twice for 15 minutes each in denaturing buffer (1.5 M NaCl, 0.5 M NaOH). DNA was transferred to HyBond-N+ membrane (GE Healthcare/Amersham) for 48 hours in fresh denaturing buffer. DNA was UV-crosslinked to membranes using the auto-crosslink setting on a Stratagene Stratalinker prior to hybridization.

A 500bp fragment spanning monomers 9-12 of DXZ1 was generated by PCR (*84*) and labeled overnight at 37°C with digoxygenin-11-dUTP using DIG High Prime (Sigma-Aldrich). Labeling reactions were purified using the High Pure PCR purification kit (Roche).

Membranes were pre-hybridized for 45-60 minutes in glass hybridization bottles containing 20mL ExpressHyb buffer (Clontech) at 63°C. Pre-hybridization buffer was replaced with 20mL of fresh ExpressHyb containing 300-350ng of labeled probe that has been denatured at 95°C for 10 minutes. The probe was allowed to hybridize to the membrane at 63°C overnight in a hybridization oven. Membranes were washed at 68°C twice for 20 minutes in 2X SSC/0.1% sodium dodecyl sulfate (SDS), followed by a single high stringency wash in 0.2X SSC/0.1% SDS for 15 minutes at 68°C. Membranes were blocked in 1X Western blocking reagent (Roche) in maleic acid buffer (0.1 M maleic acid, 0.15 M NaCl, pH 7.5) for 60-90 minutes at room temperature, then incubated for 60 minutes in blocking buffer with anti-digoxygenin-alkaline phosphatase (Roche, 1:2000). Chemiluminescent detection was performed using 5mL of CDP-Star ready-to-use reagent (Tropix). Membranes were imaged on a G:Box using GeneSys software (Syngene) for direct image analysis. Images were adjusted (leveled to curves) and labeled in Adobe Photoshop.

#### ddPCR

Genomic DNA was isolated using the DNeasy Blood & Tissue Kit (Qiagen). DNA was quantified using Qubit Fluorometer with Qubit dsDNA HS Assay (Invitrogen). Primers, gDNA concentrations, and restriction enzymes are listed Table S1-2. EvaGreen ddPCR reactions were performed using the manufacturer’s protocol (Bio-Rad). Mastermixes were simultaneously prepared for HPRT1 and the gene of interest which were then incubated for 15 minutes to allow for enzymatic restriction digestion. Statistics were performed using the confidence interval calculated by the Quantasoft software and applying it to Taylor's expansion (Fig. S38).

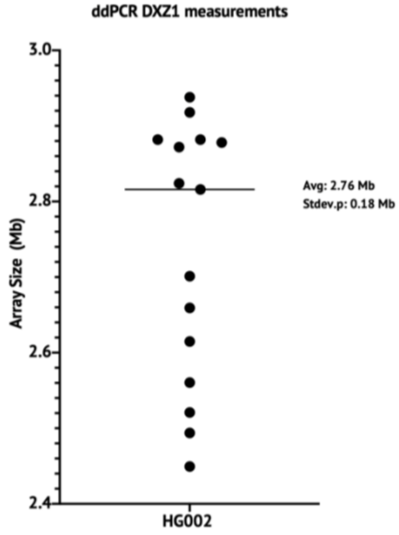


**Fig. S38: ddPCR estimate of 1N DXZ1 copy number.** Mean ± s.d. is shown. Droplet digital PCR rDNA copy number estimates for HG002 cell line calculated by DXZ1 measurements normalized to the single copy gene HPRT1 is consistent with the assembly size of 3.1 Mbp.

#### Alignment-based validation

**A**

**B**

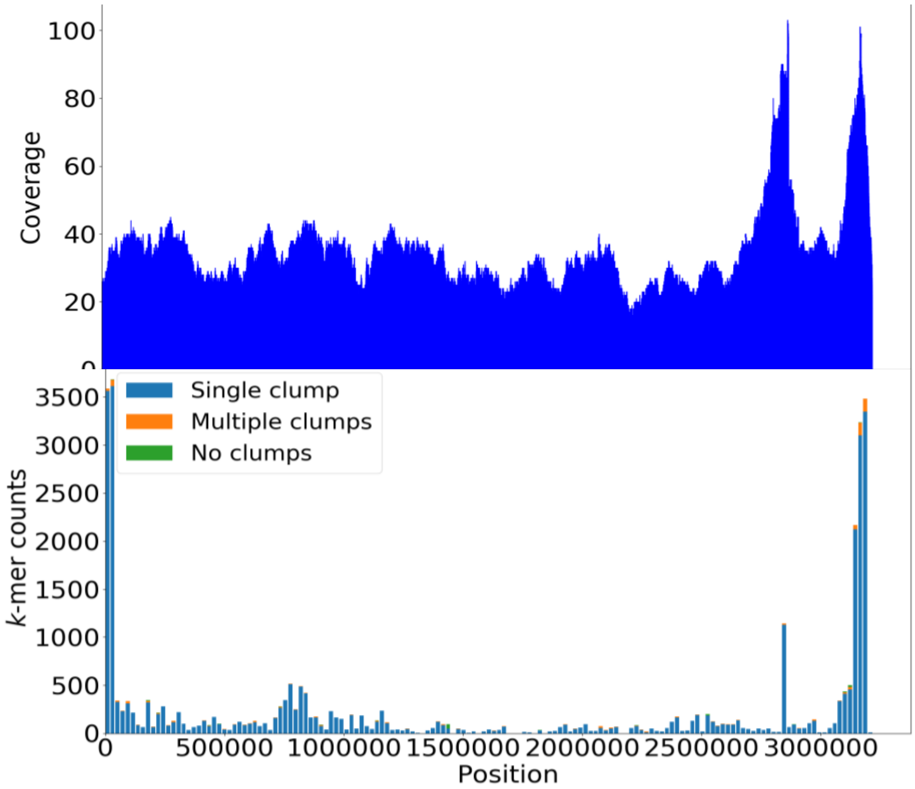

**Fig. S39: TandemTools validation of HG002 CenX.** (A) Coverage by ONT mapped reads to the assembly, with the position set to 0 at the start of the centromeric satellite. There are two coverage peaks at 2.8 and 3.1 Mbp corresponding to LINE elements but the rest of the array shows even coverage. (B) k-mer validation of the centromeric array. Pileups of multiple clumps, as defined by TandemTools (*30*), which would be indicative of mis-assemblies, are absent from the figure.

TandemTools (*30*) was run to validate the HG002 centromeric satellite assembly using ONT data. The coverage plot (Fig. S39A) showed no suspicious coverage dips or spikes, except for two peaks at 2.8 and 3.1 Mbp corresponding to LINE elements. The plot (Fig. S39B) showed high base-calling quality across the whole centromere (usually represented by k-mers forming no clumps) and suggested an absence of collapsed repeats (usually represented by k-mers forming multiple clumps). Neither this analysis nor comparison to CHM13 revealed any errors or high frequency heterozygous sites in the centromeric satellite assembly.

4. A. Auton, G. R. Abecasis, D. M. Altshuler, R. M. Durbin, G. R. Abecasis, D. R. Bentley, A. Chakravarti, A. G. Clark, P. Donnelly, E. E. Eichler, P. Flicek, S. B. Gabriel, R. A. Gibbs, E. D. Green, M. E. Hurles, B. M. Knoppers, J. O. Korbel, E. S. Lander, C. Lee, H. Lehrach, E. R. Mardis, G. T. Marth, G. A. McVean, D. A. Nickerson, J. P. Schmidt, S. T. Sherry, J. Wang, R. K. Wilson, R. A. Gibbs, E. Boerwinkle, H. Doddapaneni, Y. Han, V. Korchina, C. Kovar, S. Lee, D. Muzny, J. G. Reid, Y. Zhu, J. Wang, Y. Chang, Q. Feng, X. Fang, X. Guo, M. Jian, H. Jiang, X. Jin, T. Lan, G. Li, J. Li, Y. Li, S. Liu, X. Liu, Y. Lu, X. Ma, M. Tang, B. Wang, G. Wang, H. Wu, R. Wu, X. Xu, Y. Yin, D. Zhang, W. Zhang, J. Zhao, M. Zhao, X. Zheng, E. S. Lander, D. M. Altshuler, S. B. Gabriel, N. Gupta, N. Gharani, L. H. Toji, N. P. Gerry, A. M. Resch, P. Flicek, J. Barker, L. Clarke, L. Gil, S. E. Hunt, G. Kelman, E. Kulesha, R. Leinonen, W. M. McLaren, R. Radhakrishnan, A. Roa, D. Smirnov, R. E. Smith, I. Streeter, A. Thormann, I. Toneva, B. Vaughan, X. Zheng-Bradley, D. R. Bentley, R. Grocock, S. Humphray, T. James, Z. Kingsbury, H. Lehrach, R. Sudbrak, M. W. Albrecht, V. S. Amstislavskiy, T. A. Borodina, M. Lienhard, F. Mertes, M. Sultan, B. Timmermann, M.-L. Yaspo, E. R. Mardis, R. K. Wilson, L. Fulton, R. Fulton, S. T. Sherry, V. Ananiev, Z. Belaia, D. Beloslyudtsev, N. Bouk, C. Chen, D. Church, R. Cohen, C. Cook, J. Garner, T. Hefferon, M. Kimelman, C. Liu, J. Lopez, P. Meric, C. O'Sullivan, Y. Ostapchuk, L. Phan, S. Ponomarov, V. Schneider, E. Shekhtman, K. Sirotkin, D. Slotta, H. Zhang, G. A. McVean, R. M. Durbin, S. Balasubramaniam, J. Burton, P. Danecek, T. M. Keane, A. Kolb-Kokocinski, S. McCarthy, J. Stalker, M. Quail, J. P. Schmidt, C. J. Davies, J. Gollub, T. Webster, B. Wong, Y. Zhan, A. Auton, C. L. Campbell, Y. Kong, A. Marcketta, R. A. Gibbs, F. Yu, L. Antunes, M. Bainbridge, D. Muzny, A. Sabo, Z. Huang, J. Wang, L. J. M. Coin, L. Fang, X. Guo, X. Jin, G. Li, Q. Li, Y. Li, Z. Li, H. Lin, B. Liu, R. Luo, H. Shao, Y. Xie, C. Ye, C. Yu, F. Zhang, H. Zheng, H. Zhu, C. Alkan, E. Dal, F. Kahveci, G. T. Marth, E. P. Garrison, D. Kural, W.-P. Lee, W. Fung Leong, M. Stromberg, A. N. Ward, J. Wu, M. Zhang, M. J. Daly, M. A. DePristo, R. E. Handsaker, D. M. Altshuler, E. Banks, G. Bhatia, G. del Angel, S. B. Gabriel, G. Genovese, N. Gupta, H. Li, S. Kashin, E. S. Lander, S. A. McCarroll, J. C. Nemesh, R. E. Poplin, S. C. Yoon, J. Lihm, V. Makarov, A. G. Clark, S. Gottipati, A. Keinan, J. L. Rodriguez-Flores, J. O. Korbel, T. Rausch, M. H. Fritz, A. M. Stütz, P. Flicek, K. Beal, L. Clarke, A. Datta, J. Herrero, W. M. McLaren, G. R. S. Ritchie, R. E. Smith, D. Zerbino, X. Zheng-Bradley, P. C. Sabeti, I. Shlyakhter, S. F. Schaffner, J. Vitti, D. N. Cooper, E. V. Ball, P. D. Stenson, D. R. Bentley, B. Barnes, M. Bauer, R. Keira Cheetham, A. Cox, M. Eberle, S. Humphray, S. Kahn, L. Murray, J. Peden, R. Shaw, E. E. Kenny, M. A. Batzer, M. K. Konkel, J. A. Walker, D. G. MacArthur, M. Lek, R. Sudbrak, V. S. Amstislavskiy, R. Herwig, E. R. Mardis, L. Ding, D. C. Koboldt, D. Larson, K. Ye, S.

Gravel, The 1000 Genomes Project Consortium, Corresponding authors, Steering committee, Production group, Baylor College of Medicine, BGI-Shenzhen, Broad Institute of MIT and Harvard, Coriell Institute for Medical Research, E. B. I. European Molecular Biology Laboratory, Illumina, Max Planck Institute for Molecular Genetics, McDonnell Genome Institute at Washington University, US National Institutes of Health, University of Oxford, Wellcome Trust Sanger Institute, Analysis group, Affymetrix, Albert Einstein College of Medicine, Bilkent University, Boston College, Cold Spring Harbor Laboratory, Cornell University, European Molecular Biology Laboratory, Harvard University, Human Gene Mutation Database, Icahn School of Medicine at Mount Sinai, Louisiana State University, Massachusetts General Hospital, McGill University, N. National Eye Institute, A global reference for human genetic variation. *Nature*. **526**, 68–74 (2015).

5. E. Lowy-Gallego, S. Fairley, X. Zheng-Bradley, M. Ruffier, L. Clarke, P. Flicek, Variant calling on the GRCh38 assembly with the data from phase three of the 1000 Genomes Project [version 2; peer review: 2 approved]. *Wellcome Open Res.* **4** (2019), doi:10.12688/wellcomeopenres.15126.2.

6. K. A. Frazer, D. G. Ballinger, D. R. Cox, D. A. Hinds, L. L. Stuve, R. A. Gibbs, J. W. Belmont, A. Boudreau, P. Hardenbol, S. M. Leal, S. Pasternak, D. A. Wheeler, T. D. Willis, F. Yu, H. Yang, C. Zeng, Y. Gao, H. Hu, W. Hu, C. Li, W. Lin, S. Liu, H. Pan, X. Tang, J. Wang, W. Wang, J. Yu, B. Zhang, Q. Zhang, H. Zhao, H. Zhao, J. Zhou, S. B. Gabriel, R. Barry, B. Blumenstiel, A. Camargo, M. Defelice, M. Faggart, M. Goyette, S. Gupta, J. Moore, H. Nguyen, R. C. Onofrio, M. Parkin, J. Roy, E. Stahl, E. Winchester, L. Ziaugra, D. Altshuler, Y. Shen, Z. Yao, W. Huang, X. Chu, Y. He, L. Jin, Y. Liu, Y. Shen, W. Sun, H. Wang, Y. Wang, Y. Wang, X. Xiong, L. Xu, M. M. Y. Waye, S. K. W. Tsui, H. Xue, J. T.-F. Wong, L. M. Galver, J.-B. Fan, K. Gunderson, S. S. Murray, A. R. Oliphant, M. S. Chee, A. Montpetit, F. Chagnon, V. Ferretti, M. Leboeuf, J.-F. Olivier, M. S. Phillips, S. Roumy, C. Sallée, A. Verner, T. J. Hudson, P.-Y. Kwok, D. Cai, D. C. Koboldt, R. D. Miller, L. Pawlikowska, P. Taillon-Miller, M. Xiao, L.-C. Tsui, W. Mak, Y. Qiang Song, P. K. H. Tam, Y. Nakamura, T. Kawaguchi, T. Kitamoto, T. Morizono, A. Nagashima, Y. Ohnishi, A. Sekine, T. Tanaka, T. Tsunoda, P. Deloukas, C. P. Bird, M. Delgado, E. T. Dermitzakis, R. Gwilliam, S. Hunt, J. Morrison, D. Powell, B. E. Stranger, P. Whittaker, D. R. Bentley, M. J. Daly, P. I. W. de Bakker, J. Barrett, Y. R. Chretien, J. Maller, S. McCarroll, N. Patterson, I. Pe’er, A. Price, S. Purcell, D. J. Richter, P. Sabeti, R. Saxena, S. F. Schaffner, P. C. Sham, P. Varilly, D. Altshuler, L. D. Stein, L. Krishnan, A. Vernon Smith, M. K. Tello-Ruiz, G. A. Thorisson, A. Chakravarti, P. E. Chen, D. J. Cutler, C. S. Kashuk, S. Lin, G. R. Abecasis, W. Guan, Y. Li, H. M. Munro, Z. Steve Qin, D. J. Thomas, G. McVean, A. Auton, L. Bottolo, N. Cardin, S. Eyheramendy, C. Freeman, J. Marchini, S. Myers, C. Spencer, M. Stephens, P. Donnelly, L. R. Cardon, G. Clarke, D. M. Evans, A. P. Morris, B. S. Weir, T. Tsunoda, T. Johnson, J. C. Mullikin, S. T. Sherry, M. Feolo, A. Skol, H. Zhang, C. Zeng, H. Zhao, I. Matsuda, Y. Fukushima, D. R. Macer, E. Suda, C. N. Rotimi, C. A. Adebamowo, I. Ajayi, T. Aniagwu, P. A. Marshall, C. Nkwodimmah, C. D. M. Royal, M. F. Leppert, M. Dixon, A. Peiffer, R. Qiu, A. Kent, K. Kato, N. Niikawa, I. F. Adewole, B. M. Knoppers, M. W. Foster, E. Wright Clayton, J. Watkin, R. A. Gibbs, J. W. Belmont, D. Muzny, L. Nazareth, E. Sodergren, G. M. Weinstock, D. A. Wheeler, I. Yakub, S. B. Gabriel, R. C. Onofrio, D. J. Richter, L. Ziaugra, B. W. Birren, M. J. Daly, D. Altshuler, R. K. Wilson, L. L. Fulton, J. Rogers, J. Burton, N. P. Carter, C. M. Clee, M. Griffiths, M. C. Jones, K. McLay, R. W. Plumb, M. T. Ross, S. K. Sims, D. L. Willey, Z. Chen, H. Han, L. Kang, M. Godbout, J. C. Wallenburg, P. L’Archevêque, G. Bellemare, K. Saeki, H. Wang, D. An, H. Fu, Q. Li, Z. Wang, R. Wang, A. L. Holden, L. D. Brooks, J. E. McEwen, M. S. Guyer, V. Ota Wang, J. L. Peterson, M. Shi, J. Spiegel, L. M. Sung, L. F. Zacharia, F. S. Collins, K. Kennedy, R.

Jamieson, The International HapMap Consortium, Genotyping centres: Perlegen Sciences, Baylor College of Medicine and ParAllele BioScience, Beijing Genomics Institute, Broad Institute of Harvard and Massachusetts Institute of Technology, Chinese National Human Genome Center at Beijing, Chinese National Human Genome Center at Shanghai, Chinese University of Hong Kong, Hong Kong University of Science and Technology, Illumina, McGill University and Génome Québec Innovation Centre, University of California at San Francisco and Washington University, University of Hong Kong, University of Tokyo and RIKEN, Wellcome Trust Sanger Institute, Analysis groups: Broad Institute, Cold Spring Harbor Laboratory, Johns Hopkins University School of Medicine, University of Michigan, University of Oxford, W. T. C. for H. G. University of Oxford, RIKEN, US National Institutes of Health, US National Institutes of Health National Center for Biotechnology Information, Community engagement/public consultation and sample collection groups: Beijing Normal University and Beijing Genomics Institute, E. E. I. Health Sciences University of Hokkaido and Shinshu University, Howard University and University of Ibadan, University of Utah, legal and social issues: C. A. of S. S. Ethical, Genetic Interest Group, Kyoto University, Nagasaki University, University of Ibadan School of Medicine, University of Montréal, University of Oklahoma, Vanderbilt University, Wellcome Trust, SNP discovery: Baylor College of Medicine, Washington University, Scientific management: Chinese Academy of Sciences, Genome Canada, Génome Québec, C. Japanese Ministry of Education Sports, Science and Technology, Ministry of Science and Technology of the People's Republic of China, The Human Genetic Resource Administration of China, The SNP Consortium, A second generation human haplotype map of over 3.1 million SNPs. *Nature*. **449**, 851–861 (2007).

7. H. Li, J. M. Bloom, Y. Farjoun, M. Fleharty, L. Gauthier, B. Neale, D. MacArthur, A synthetic-diploid benchmark for accurate variant-calling evaluation. *Nat. Methods*. **15**, 595–597 (2018).

8. S. Aganezov, A complete human reference genome improves variant calling for population and clinical genomics. *Prep* (2021).

21. V. A. Schneider, T. Graves-Lindsay, K. Howe, N. Bouk, H. C. Chen, P. A. Kitts, T. D. Murphy, K. D. Pruitt, F. Thibaud-Nissen, D. Albracht, R. S. Fulton, M. Kremitzki, V. Magrini, C. Markovic, S. McGrath, K. M. Steinberg, K. Auger, W. Chow, J. Collins, G. Harden, T. Hubbard, S. Pelan, J. T. Simpson, G. Threadgold, J. Torrance, J. M. Wood, L. Clarke, S. Koren, M. Boitano, P. Peluso, H. Li, C. S. Chin, A. M. Phillippy, R. Durbin, R. K. Wilson, P. Flicek, E. E. Eichler, D. M. Church, Evaluation of GRCh38 and de novo haploid genome assemblies demonstrates the enduring quality of the reference assembly. *Genome Res*. **27** (2017), pp. 849–864.

22. C. Jain, S. Koren, A. Dilthey, A. M. Phillippy, S. Aluru, A fast adaptive algorithm for computing whole-genome homology maps. *Bioinformatics*. **34**, i748–i756 (2018).

23. K. H. Miga, S. Koren, A. Rhie, M. R. Vollger, A. Gershman, A. Bzikadze, S. Brooks, E. Howe, D. Porubsky, G. A. Logsdon, V. A. Schneider, T. Potapova, J. Wood, W. Chow, J. Armstrong, J. Fredrickson, E. Pak, K. Tigyi, M. Kremitzki, C. Markovic, V. Maduro, A. Dutra, G. G. Bouffard, A. M. Chang, N. F. Hansen, A. B. Wilfert, F. Thibaud-Nissen, A. D. Schmitt, J.-M. Belton, S. Selvaraj, M. Y. Dennis, D. C. Soto, R. Sahasrabudhe, G. Kaya, J. Quick, N. J. Loman, N. Holmes, M. Loose, U. Surti, R. ana Risques, T. A. G. Lindsay, R. Fulton, I. Hall, B. Paten, K. Howe, W. Timp, A. Young, J. C. Mullikin, P. A. Pevzner, J. L.

Gerton, B. A. Sullivan, E. E. Eichler, A. M. Phillippy, Telomere-to-telomere assembly of a complete human X chromosome. *Nature* (2020), doi:10.1038/s41586-020-2547-7.

33. A. Rhie, S. A. McCarthy, O. Fedrigo, J. Damas, G. Formenti, S. Koren, M. Uliano-Silva, W. Chow, A. Fungtammasan, J. Kim, C. Lee, B. J. Ko, M. Chaisson, G. L. Gedman, L. J. Cantin, F. Thibaud-Nissen, L. Haggerty, I. Bista, M. Smith, B. Haase, J. Mountcastle, S. Winkler, S. Paez, J. Howard, S. C. Vernes, T. M. Lama, F. Grutzner, W. C. Warren, C. N. Balakrishnan, D. Burt, J. M. George, M. T. Biegler, D. Iorns, A. Digby, D. Eason, B. Robertson, T. Edwards, M. Wilkinson, G. Turner, A. Meyer, A. F. Kautt, P. Franchini, H. W. Detrich, H. Svardal, M. Wagner, G. J. P. Naylor, M. Pippel, M. Malinsky, M. Mooney, M. Simbirsky, B. T. Hannigan, T. Pesout, M. Houck, A. Misuraca, S. B. Kingan, R. Hall, Z. Kronenberg, I. Sović, C. Dunn, Z. Ning, A. Hastie, J. Lee, S. Selvaraj, R. E. Green, N. H. Putnam, I. Gut, J. Ghurye, E. Garrison, Y. Sims, J. Collins, S. Pelan, J. Torrance, A. Tracey, J. Wood, R. E. Dagnew, D. Guan, S. E. London, D. F. Clayton, C. V. Mello, S. R. Friedrich, P. V. Lovell, E. Osipova, F. O. Al-Ajli, S. Secomandi, H. Kim, C. Theofanopoulou, M. Hiller, Y. Zhou, R. S. Harris, K. D. Makova, P. Medvedev, J. Hoffman, P. Masterson, K. Clark, F. Martin, K. Howe, P. Flicek, B. P. Walenz, W. Kwak, H. Clawson, M. Diekhans, L. Nassar, B. Paten, R. H. S. Kraus, A. J. Crawford, M. T. P. Gilbert, G. Zhang, B. Venkatesh,

R. W. Murphy, K.-P. Koepfli, B. Shapiro, W. E. Johnson, F. Di Palma, T. Marques-Bonet, E. C. Teeling, T. Warnow, J. M. Graves, O. A. Ryder, D. Haussler, S. J. O'Brien, J. Korlach, H. A. Lewin, K. Howe, E. W. Myers, R. Durbin, A. M. Phillippy, E. D. Jarvis, Towards complete and error-free genome assemblies of all vertebrate species. *Nature*. **592**, 737–746 (2021).

34. A. M. McCartney, Chasing perfection: validation and polishing strategies for telomere-to-telomere genome assemblies. *Prep* (2021).

35. H. Li, Aligning sequence reads, clone sequences and assembly contigs with BWA-MEM. *ArXiv Prepr. ArXiv13033997* (2013).

36. R. Poplin, P.-C. Chang, D. Alexander, S. Schwartz, T. Colthurst, A. Ku, D. Newburger, J. Dijamco, N. Nguyen, P. T. Afshar, S. S. Gross, L. Dorfman, C. Y. McLean, M. A. DePristo, A universal SNP and small-indel variant caller using deep neural networks. *Nat. Biotechnol.* **36**, 983–987 (2018).

37. K. Shafin, T. Pesout, P.-C. Chang, M. Nattestad, A. Kolesnikov, S. Goel, G. Baid, J. M. Eizenga, K. H. Miga, P. Carnevali, M. Jain, A. Carroll, B. Paten, Haplotype-aware variant calling enables high accuracy in nanopore long-reads using deep neural networks. *bioRxiv* (2021), doi:10.1101/2021.03.04.433952.

38. G. Formenti, Merfin: improved variant filtering and polishing via k-mer validation. *Prep* (2021).

39. S. Zarate, A. Carroll, O. Krashenina, F. J. Sedlazeck, G. Jun, W. Salerno, E. Boerwinkle, R. Gibbs, Parliament2: Fast Structural Variant Calling Using Optimized Combinations of Callers. *bioRxiv*, 424267 (2018).

40. F. J. Sedlazeck, P. Rescheneder, M. Smolka, H. Fang, M. Nattestad, A. von Haeseler, M. C. Schatz, Accurate detection of complex structural variations using single-molecule sequencing. *Nat. Methods*. **15**, 461–468 (2018).

41. M. Kirsche, Jasmine: Population-scale structural variant comparison and analysis. *Prep* (2021).

42. J. T. Robinson, H. Thorvaldsdóttir, W. Winckler, M. Guttman, E. S. Lander, G. Getz, J. P. Mesirov, Integrative genomics viewer. *Nat. Biotechnol.* **29**, 24–26 (2011).

43. H. Li, B. Handsaker, A. Wysoker, T. Fennell, J. Ruan, N. Homer, G. Marth, G. Abecasis, R. Durbin, S. Genome Project Data Processing, The Sequence Alignment/Map format and SAMtools. *Bioinformatics*. **25** (2009), pp. 2078–9.

44. N. Altemose, Genetic and epigenetic maps of endogenous human centromeres. *Prep* (2021).

59. M. R. Vollger, Segmental duplications and their variation in a complete human genome. *Prep* (2021).

60. B. Gel, E. Serra, karyoploteR: an R/Bioconductor package to plot customizable genomes displaying arbitrary data. *Bioinformatics*. **33**, 3088–3090 (2017).

61. R. S. Harris, thesis, Pennsylvania State University, USA (2007).

62. B. Paten, D. Earl, N. Nguyen, M. Diekhans, D. Zerbino, D. Haussler, Cactus: Algorithms for genome multiple sequence alignment. *Genome Res.* **21**, 1512–1528 (2011).

63. S. Kovaka, A. V. Zimin, G. M. Pertea, R. Razaghi, S. L. Salzberg, M. Pertea, Transcriptome assembly from long-read RNA-seq alignments with StringTie2. *Genome Biol.* **20**, 278 (2019).

64. I. T. Fiddes, J. Armstrong, M. Diekhans, S. Nachtweide, Z. N. Kronenberg, J. G. Underwood, D. Gordon, D. Earl, T. Keane, E. E. Eichler, D. Haussler, M. Stanke, B. Paten, Comparative Annotation Toolkit (CAT)—simultaneous clade and personal genome annotation. *Genome Res.* (2018), doi:10.1101/gr.233460.117.

65. M. Stanke, M. Diekhans, R. Baertsch, D. Haussler, Using native and syntenically mapped cDNA alignments to improve de novo gene finding. *Bioinformatics*. **24**, 637–644 (2008).

66. A. Frankish, M. Diekhans, A.-M. Ferreira, R. Johnson, I. Jungreis, J. Loveland, J. M. Mudge, C. Sisu, J. Wright, J. Armstrong, I. Barnes, A. Berry, A. Bignell, S. Carbonell Sala, J. Chrast, F. Cunningham, T. Di Domenico, S. Donaldson, I. T. Fiddes, C. García Girón, J. M. Gonzalez, T. Grego, M. Hardy, T. Hourlier, T. Hunt, O. G. Izuogu, J. Lagarde, F. J. Martin, L. Martínez, S. Mohanan, P. Muir, F. C. P. Navarro, A. Parker, B. Pei, F. Pozo, M. Ruffier, B. M. Schmitt, E. Stapleton, M.-M. Suner, I. Sycheva, B. Uszczynska-Ratajczak, J. Xu, A. Yates, D. Zerbino, Y. Zhang, B. Aken, J. S. Choudhary, M. Gerstein, R. Guigó, T. J. P. Hubbard, M. Kellis, B. Paten, A. Reymond, M. L. Tress, P. Flicek, GENCODE reference annotation for the human and mouse genomes. *Nucleic Acids Res.* **47**, D766–D773 (2019).

78. M. Nei, S. Kumar, *Molecular evolution and phylogenetics* (Oxford university press, 2000).

79. R. Patro, G. Duggal, M. I. Love, R. A. Irizarry, C. Kingsford, Salmon provides fast and bias-aware quantification of transcript expression. *Nat. Methods*. **14**, 417–419 (2017).

80. C. Soneson, M. Love, M. Robinson, Differential analyses for RNA-seq: transcript-level estimates improve gene-level inferences [version 2; peer review: 2 approved]. *F1000Research*. **4** (2016), doi:10.12688/f1000research.7563.2.

81. J. M. Zook, D. Catoe, J. McDaniel, L. Vang, N. Spies, A. Sidow, Z. Weng, Y. Liu, C. E. Mason, N. Alexander, E. Henaff, A. B. R. McIntyre, D. Chandramohan, F. Chen, E. Jaeger, A. Moshrefi, K. Pham, W. Stedman, T. Liang, M. Saghbini, Z. Dzakula, A. Hastie, H. Cao, G. Deikus, E. Schadt, R. Sebra, A. Bashir, R. M. Truty, C. C. Chang, N. Gulbahce, K. Zhao, S. Ghosh, F. Hyland, Y. Fu, M. Chaisson, C. Xiao, J. Trow, S. T. Sherry, A. W. Zaranek, M. Ball, J. Bobe, P. Estep, G. M. Church, P. Marks, S. Kyriazopoulou-Panagiotopoulou, G. X. Y. Zheng, M. Schnall-Levin, H. S. Ordonez, P. A. Mudivarti, K. Giorda, Y. Sheng, K. B. Rypdal, M. Salit, Extensive sequencing of seven human genomes to characterize benchmark reference materials. *Sci. Data*. **3**, 1–26 (2016).

82. A. M. Wenger, P. Peluso, W. J. Rowell, P.-C. Chang, R. J. Hall, G. T. Concepcion, J. Ebler, A. Fungtammasan, A. Kolesnikov, N. D. Olson, A. Töpfer, M. Alonge, M. Mahmoud,

Y. Qian, C.-S. Chin, A. M. Phillippy, M. C. Schatz, G. Myers, M. A. DePristo, J. Ruan, T. Marschall, F. J. Sedlazeck, J. M. Zook, H. Li, S. Koren, A. Carroll, D. R. Rank, M. W. Hunkapiller, Accurate circular consensus long-read sequencing improves variant detection and assembly of a human genome. *Nat. Biotechnol.* **37** (2019), pp. 1155–1162.

83. L. L. Sullivan, C. D. Boivin, B. Mravinac, I. Y. Song, B. A. Sullivan, Genomic size of CENP-A domain is proportional to total alpha satellite array size at human centromeres and expands in cancer cells. *Chromosome Res.* **19**, 457 (2011).

84. B. Mravinac, L. L. Sullivan, J. W. Reeves, C. M. Yan, K. S. Kopf, C. J. Farr, M. G. Schueler, B. A. Sullivan, Histone Modifications within the Human X Centromere Region. *PLOS ONE*. **4**, e6602 (2009).
